# ExpertTwins: Auditable Long Horizon Scientific Panels through Grounded Expertise

**DOI:** 10.64898/2026.10.06.757114

**Authors:** Ali Foroughi pour, Abbas Shirinifard

## Abstract

Multi-agent language-model systems increasingly assign roles to agents and let them deliberate. A role label, however, does not specify which evidence an agent can bring or which standard it applies before accepting a conclusion. Agents that read one another’s outputs tend to converge. We present ExpertTwins, a framework in which each seat of a scientific panel is defined by an inspectable evidence policy. Discipline seats retrieve under field-specific queries, and scientist-associated seats retrieve from attributed publications and a citation-derived scholarly neighborhood. Private initial packets, structured exchange, cited-source disclosure, deterministic quotation checks, and peer source review turn deliberation into an auditable record, which we also render as interactive idea-flow maps. Scientist associations denote publication-grounded perspectives, not replicas or endorsements of named researchers. In a nine-seat, four-turn neuroblastoma panel, 438 of 439 claims passed all attached quotation checks, and seats withdrew or defended positions when challenged on their evidence. A ten-turn panel sustained evidence acquisition and differentiated recommendations without conversational collapse. Across six base models given byte-identical initial packets, four sustained 97.9–99.8% quotation grounding, and at least 91% on every turn, whereas Haiku 4.5 failed by paraphrase and GPT-5 mini by silence. Agreement among strong models fell from 31% of claims at turn one to 5% at turn four. With follow-up questions held fixed, it stayed at 28–33% through turn three and fell to 20% only at the closing turn. Stress runs showed the collapse guardrail escalating through its full intervention ladder when consensus was forced, but missing deference to one unusually strong seat. On idea-flow maps, the forced consensus appeared as parallel lanes of identical ideas, although quotation grounding and participation balance stayed at their maxima. We also re-implemented two close published systems and ran them on matched models, seats, and questions. A multi-persona debate cited only real records, yet none of its cited sentences carried a checkable quotation and it had no way to abstain. An author-memory fusion system ran only after its fact checker was split by seat to fit in context. Its fused answer kept about two-thirds of the fact-checked specialist claims, answered 36 of 60 question parts on which our seats had abstained, and showed the reader 19% of its agents’ disagreements. In a computational pathology case study, six scientist-associated agents built distinct pipelines to predict MYCN amplification from whole-slide images. Across six base models, seat-by-model interaction explained 57% of internal variance, and internal rank poorly predicted external rank (*r* = 0.18 across 36 nominees). These results establish differentiated, source-traceable scientific reasoning as an operational capability and identify the base model as a parameter that must be reported with the protocol.

## 1. Introduction

The most useful product of a scientific discussion is often not its conclusion. It may be a confound that changes an analysis, a method no one had considered, or a distinction between what a result shows and what its authors would like it to mean. Such contributions come from differences among participants in the literature they command, the techniques they trust, and the evidence they require. An automated scientific panel should be judged on whether it supplies these differences and makes their consequences visible.

Language-model systems already support literature synthesis, hypothesis development, and experimental planning. Retrieval-augmented generation supplies evidence beyond a model’s parameters [1], and literature agents such as PaperQA2 and OpenScholar connect answers to sources [2,3]. Virtual Lab and AI co-scientist coordinate agents around scientific tasks [4,5]. Robin combines literature and data-analysis agents and produced an experimentally validated therapeutic candidate [87], and Kosmos sustains long autonomous sessions while citing each statement to code or primary literature [88]. Multi-agent debate improves factual accuracy on question-answering tasks [30], Tool-MAD gives each debater a distinct retrieval tool [43], and deliberation in which agents hold disjoint private evidence lowers correlated error relative to a shared pool [89].

Most of these systems start from a role description, which is a weak specification of expertise. Calling an agent an immunologist does not determine what it retrieves or which distinctions it preserves. A study of 162 roles, four model families, and 2,410 factual questions found no general benefit from personas [6]. Assigned personas drift in multi-agent settings [7], and when 6,749 scientists rated machine-generated ideas derived from their own papers, asking the model to adopt their persona gave only marginal gains [90]. Communication adds a second difficulty. Sharing complete solutions erodes diversity on verifier-scored tasks [28], whereas structured exchange can help integrate distributed evidence [27]. Human groups behave similarly: discussion favors shared over unshared information [8], social influence narrows estimates without improving them [9], and Delphi methods counter both with independent initial judgments and controlled feedback [10]. Language models conform to peer agents [31,32] and to the stated views of their interlocutors [33], and decisions that share one model are homogenized [34,35]. Shared weights, training data, retrieval material, and visible peer outputs can all correlate a panel, and changing the retrieved documents does not make agents statistically independent.

ExpertTwins makes expertise an inspectable combination of evidence access and methodological remit, and makes deliberation a record that can be verified rather than trusted. Expertise is specified as a *seat*: a literature-selection policy, a permitted evidence set, and explicit scientific responsibilities. Discipline seats are associated with a field or disease domain. Scientist-associated seats, which we call *ExpertTwins*, are built from attributed publications, a citation-derived *scholarly neighborhood*, and a written methodological rubric. Seats begin from private evidence, exchange structured positions rather than drafts, and attach source text to every claim. A deterministic quotation verifier, peer source review, and turn-level monitoring then check the record (Figure 1). Literature-based discovery, which connects literatures that no single reader holds, is the intellectual context for this design [29,45].

**Figure 1.**
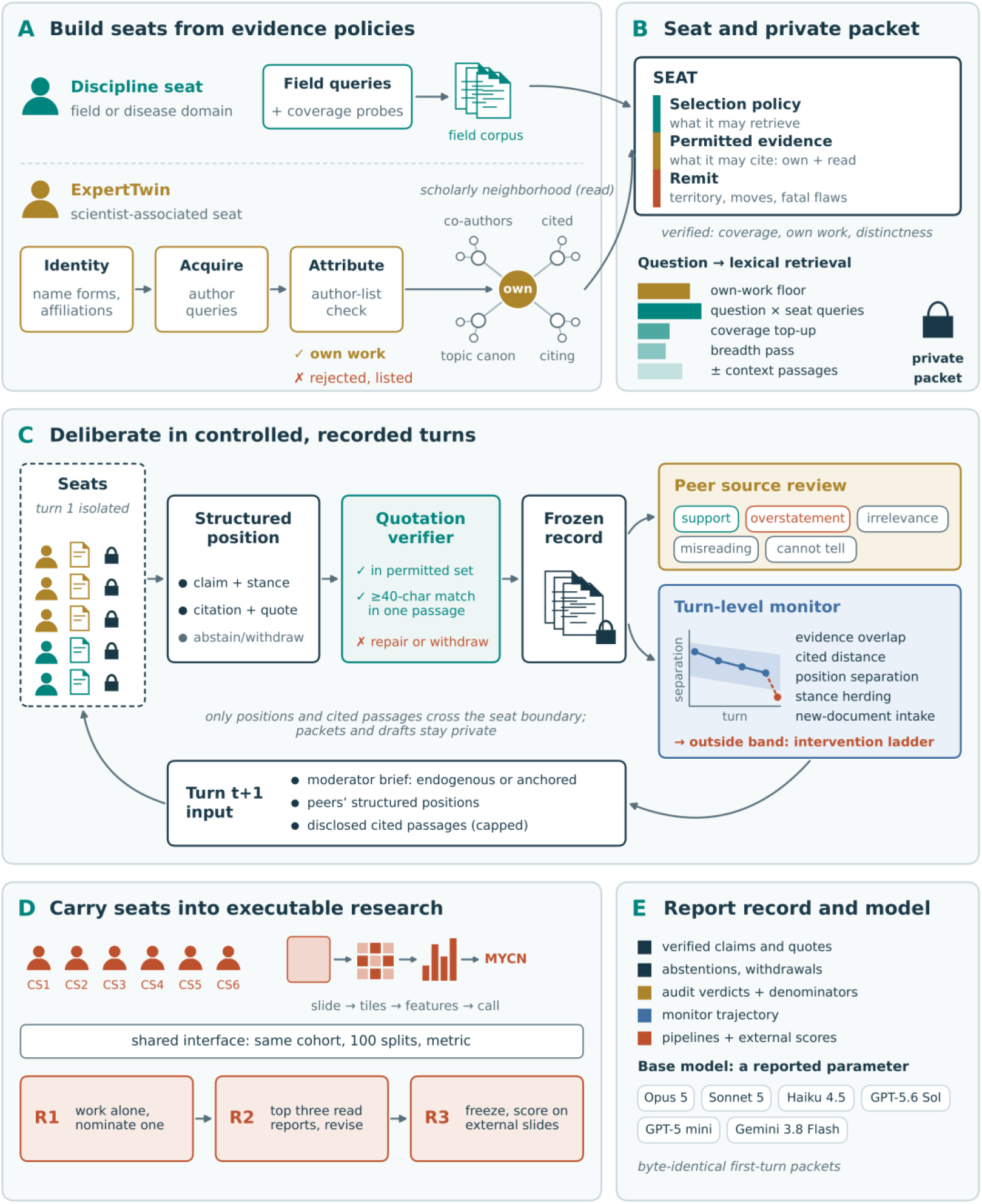
The ExpertTwins framework. (A) Seats are built from evidence policies. A discipline seat retrieves a field corpus through field queries checked by coverage probes. A scientist-associated seat (ExpertTwin) is built from an identity specification, author queries, and an author-list attribution check whose rejected works are reported. The verified own-work set (own) anchors a scholarly neighborhood (read) of most-cited references, collaborators’ works, most-cited citing works, and topic canon. (B) A seat combines a literature-selection policy, a permitted evidence set, and a written remit, and is verified for coverage, held own work, and distinctness. For each question, lexical retrieval assembles a private packet in strata: own-work floor, question-by-seat queries, coverage topup, breadth pass, and context passages. (C) Turn 1 is isolated. Each citation must pass a deterministic quotation verifier (document in the permitted set; quoted span matched in a single passage) or be repaired or withdrawn. The frozen turn is reviewed by peer seats, and a monitor compares evidence overlap, cited-evidence distance, position separation, stance herding, and new-document intake with bands derived from turn 1. A breached band makes the intervention ladder available (peer routing, differential retrieval, own-work reanchoring, anonymization, isolation, assigned dissent, termination, declaration of collapse). The next turn receives a moderator brief, written from the run’s own record (endogenous) or taken from a fixed question set (anchored), with peers’ structured positions and capped disclosed passages; packets and drafts never cross the seat boundary. (D) In a computational panel, agents CS1–CS6 build whole-slide-image pipelines for MYCN amplification under a shared cohort, split, and metric interface. Each nominates one configuration (R1), the top seats read one another’s reports and revise (R2), and frozen nominations are scored on external slides (R3). (E) The reported record: verified claims and quotations, abstentions and withdrawals, audit verdicts with denominators, the monitor trajectory, and pipelines with external scores. The base model is a reported parameter, varied across six models from byte-identical first-turn packets. The diagram is conceptual. Scientist-associated seats are not replicas or endorsements of named researchers.

Model development offers an execution test that complements deliberation. In computational pathology, tile-level classification, multiple-instance learning, and slide-level foundation models [11–14] offer several plausible choices of representation and aggregation. MYCN amplification is a clinically important marker in neuroblastoma and part of consensus risk stratification [15,46]. Predicting it from slides lets us test whether conditioning agents on individual scientists’ methods produces different pipelines.

Two published systems are closest to this work. The Multi-Persona Debate System (MPDS) induces personas from a retrieved corpus and gives each agent a role-specific pool of abstracts [24]. Its agents share one base model and follow a fixed three-round arc that is designed to converge and ends in a moderator’s consensus plan [24]. It is described as a synthesis engine, not a verification engine [24]. MirrorMind consolidates a researcher’s publications into episodic, semantic, and persona memory, evaluates the result as an author-level simulation, and delegates fact checking to reviewing agents rather than to a mechanical test against a permitted source [42]. Related systems give domain experts separate knowledge bases [25] or condition retrieval on a representation of an individual researcher [26]. Conditioning an agent on a scientist’s publications is therefore not new, and we adopt that premise rather than claim it. None of these systems, however, defines an agent by an attribution-checked evidence set whose retrieval, disclosure, and quotation can be verified after the fact. We make no claim of fidelity to the researcher.

Two of our design choices are improvements rather than new mechanisms. We keep evidence at the level of the manuscript and its passages instead of distilling it into a concept graph, which gives up abstraction but leaves every statement quotable. We also let peer seats read the source behind another seat’s citation, so a claim can be contested on its evidence rather than its phrasing. What we claim as new is treating a panel as a long-horizon process whose agreement is measured rather than produced. MirrorMind evaluates single responses [42] and MPDS stops at its third round by construction [24], so neither can **separate a consensus reached because the evidence supports it from a consensus reached because agents accommodated one another**. That distinction is our primary problem, and it becomes visible only over many turns.

We make six methodological contributions.

1. **We define expertise operationally.** A seat is a literature-selection policy, a permitted evidence set, and a methodological remit. Scientist-associated seats are built from attributed publications and their scholarly neighborhood, so differentiation enters retrieval, where a prompt alone cannot reach.
2. **We control and record the flow of information.** Seats start from private packets, exchange structured positions rather than drafts, and see each other’s sources only through mediated disclosure, so later agreement can be traced to shared inputs or to independent evidence. **An expert holding ten reasons for a position will usually offer the two that bear on what another seat has just said**, so a transcript is a selection made for an audience, not the whole reasoning. Keeping the private packet separate from the filed position preserves that difference.
3. **We make grounding and deliberation measurable.** A deterministic quotation verifier, restricted to each seat’s permitted set, establishes textual traceability, and peer source review separates a valid quotation from a supported claim. Turn-level statistics describe evidence intake, participation, stance movement, and position separation.
4. **We treat convergence as a failure mode to detect, not an outcome to deliver.** An intervention ladder escalates when separation collapses.
5. **We extend seats to executable research.** Scientist-associated agents implement, exchange, and externally evaluate computational pipelines under shared cohort, split, and metric interfaces.
6. **We render each panel as an interactive idea-flow map.** Claims are grouped into ideas, followed across turns, and linked by the stances seats filed on one another, so the course of a discussion can be traced back to its quoted evidence (Section 3.7).

The demonstrations support these claims, but do not establish an advantage. A nine-seat panel remained quote-grounded over four turns and revised an inherited consensus into one experimental design, and a ten-turn panel sustained differentiated inquiry. Six base models gave largely different but equally grounded conversations under the same corpus, roster, and rules. The moderator’s questions set the level of agreement as much as the model did. Against re-implementations of MPDS and MirrorMind, ExpertTwins was the only architecture whose claims routinely carried a checkable quoted span, and it kept abstentions and disagreements that debate could not record. Stress runs show the collapse guardrail escalating correctly when consensus is forced. The idea-flow maps make collapse visible at a glance, and reveal **a second failure mode that the guardrail misses: one unusually strong seat quietly dominating an asymmetric panel**. These results establish feasibility and characterize behavior. They do not show that more agents are intrinsically better; budget-matched comparisons remain the necessary next step.

## 2. The ExpertTwins framework

### 2.1 Design problem and principles

The framework answers three documented failures of role-based multi-agent systems: a role label changes the prompt but not the evidence [6], communication correlates agents’ outputs [28,31,32], and claims carry support that cannot be checked, of which fabricated references are the clearest case [36,37]. Four principles follow.

#### Differentiation –must enter retrieval

A role prompt that does not change the retrieval query leaves the retrieved documents unchanged (Section 3.1). A seat is therefore defined by which documents it may read and which it holds, not by how it is addressed.

#### Information flow must be controlled and recorded

The protocol fixes which fields and passages cross a seat boundary at each stage, so later agreement can be traced to shared inputs or to independent evidence.

#### Every claim must carry checkable source text

A claim is filed with quotations that a deterministic verifier can locate in a permitted document. A second, semantic layer asks whether that text warrants the claim.

#### Convergence is a trajectory to examine, not a failure to assume

A change of position can reflect evidence or conformity, and only the record can tell them apart [40]. Turn-level statistics describe the trajectory; an intervention policy exists but is not credited with observed behavior.

A further commitment governs representation: a scientist-associated seat is a publication-conditioned agent, and the framework makes no fidelity claim about the named researcher.

### 2.2 Seats: expertise as evidence access and methodological remit

A *seat* is the unit of expertise. It has a literature-selection policy, which determines what it may retrieve; a permitted evidence set, which bounds what it may cite; and a written remit, which states what it is responsible for and when it must refuse to agree. Two kinds of seat share this structure and differ in how the evidence set is built (Figures 1A–B and 2; Supplementary Methods S1 and S2).

A *discipline seat* is associated with a field or a disease domain. Its specification lists retrieval queries and coverage probes that must return hits before the seat is used. Disease-domain seats hold clinical stratification and disease-specific literature that a methodological seat would treat as peripheral; the distinction is operational and confers no validated clinical expertise. Verification also checks that each seat is distinct from those already at the table, because a seat whose retrievals largely overlap another’s is an existing seat wearing a second label.

A *scientist-associated seat*, or ExpertTwin, is built in four steps. An identity specification records the name forms, affiliations, and topics needed to recognize the scientist’s work. Author queries against literature services acquire candidate publications. *Attribution* checks each candidate against its stored author list, and rejected works, for example from common names, are reported. Finally, the verified own-work set anchors a citation-derived *scholarly neighborhood*: the works the scientist cites most often, their collaborators’ works, the most-cited works that cite them, and the most-cited works in their topics, plus curated landmark titles. The harness tags own-work sources own and neighborhood sources read; read denotes set membership, not a verified act of reading.

A seat is thus conditioned through three channels: the corpus it may retrieve, the identity named in its prompts, and the rubric it applies. Corpus access is not the only difference among agents. Publications are a record of research activity, not of a scientist. Attribution does not prove intellectual contribution, and name matching is vulnerable to incomplete metadata and common-name collisions. Cited works are not reading histories, collaborators’ papers are not shared knowledge, and later citing works are not evidence of earlier influence. The neighborhood is an explicit retrieval policy that approximates a scholarly environment, not a reconstruction of anyone’s intellectual formation. A fidelity claim would require held-out behavior from the real researcher, which we do not have. Systems that pursue fidelity fine-tune on an individual’s complete corpus to reproduce their linguistic and argumentative patterns [91], and predicting a researcher’s next papers is a weak proxy for fidelity in any case [26]. Scientist-associated seats are therefore *publication-conditioned research assistants* whose evidence can be inspected and challenged. Because the packet builder reserves space for own work, a packet’s own-work fraction is an exposure diagnostic, not a measure of expertise.

**Figure 2.**
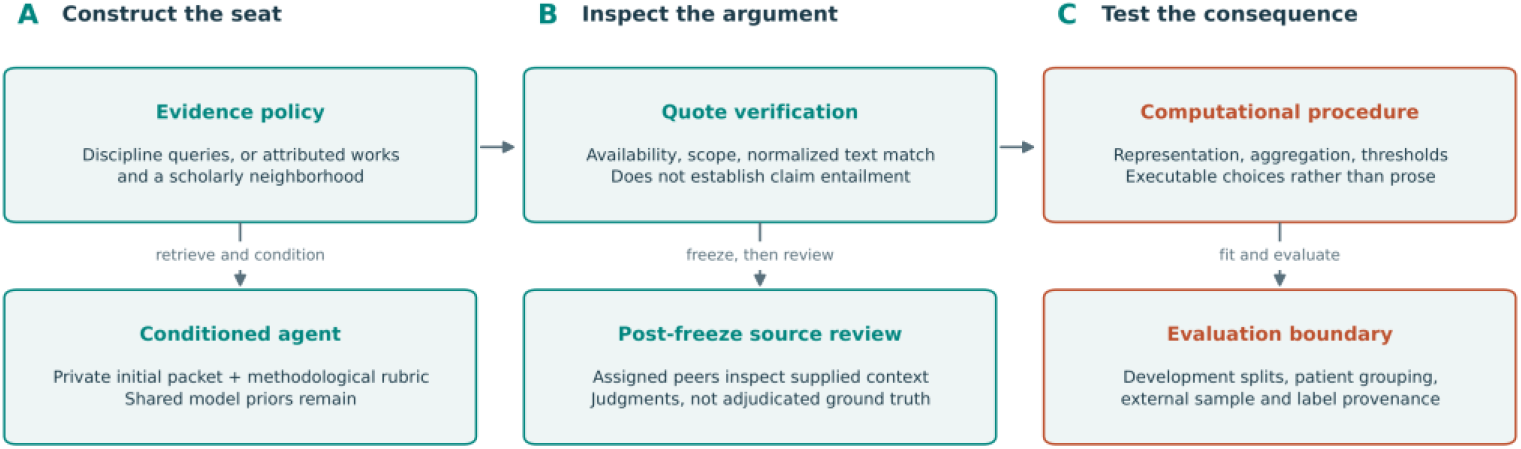
From an evidence policy to an auditable scientific result. A seat’s external evidence is selected through a discipline policy or a scientist-associated publication policy and combined with explicit instructions. Quote verification, peer source review, and computational evaluation test different properties of its outputs. Arrows show a conceptual workflow, not a causal pathway; the model’s parameterized knowledge remains available throughout. Initial packets are private; later structured exchange and cited-source disclosure are mediated separately. Scientist names identify corpus associations, not participants or endorsers.

### 2.3 Evidence packets and communication boundaries

Each seat answers a question from an *evidence packet* assembled under its policy (Figure 1B–C; Supplementary Methods S3 and S4). The packet builder retrieves lexically over the full text of the permitted corpus and keeps neighboring passages in source order. Thereby, a matched sentence is read in context. It fills the packet in strata: (a) question-conditioned retrieval inside the own-work set, up to an own-work floor; (b) retrieval across the rest of the permitted corpus with the question and the seat’s queries; (c) coverage top-ups for question terms that no retrieved passage contains; (d) a breadth pass that guarantees a minimum number of documents; and (e) context passages on both sides of each excerpt. Separate budgets for ranked hits and for the packet after context stop the hits from consuming everything, and round-robin allocation over queries stops one query from starving another. The own-work floor is a floor, not a quota: if the scientist wrote nothing the question can reach, the stratum is empty, and the emptiness is a finding. Before any model call, a packet audit reports documents, own-work documents, term coverage, and the largest single-document share, so a thin packet is visible in advance.

The protocol controls when each of three kinds of material crosses a seat boundary. A seat’s *private packet* is never broadcast. Its *structured position* (claims with stance labels, citations, and verified quotations, imported from its returned file rather than its prose) is what other seats see. *Disclosed source context* is passage text from a document another seat cited in the previous turn, anchored on the quoted passage, supplied outside the evidence budget, and capped per receiving seat. Turn one is isolated: every seat answers from its private packet alone. Later turns add the moderator’s brief, the other seats’ positions, and disclosed context. Abstentions and withdrawals are first-class outcomes, consistent with work that treats calibrated refusal as desirable [92]. Each turn is frozen before it is audited, so the audit never rewrites what it assesses, although it is not independent of model priors or shared context. The orchestration layer prepares packets, imports responses, runs deterministic checks, freezes turns, and renders reports, while model calls are dispatched outside it, so prompts, returned files, and verification decisions can all be inspected.

Follow-up questions come from one of two regimes. In the *endogenous* regime, a moderator model reads only the run’s own frozen record (verified claims, abstentions, disclosed sources, citation-check verdicts, and monitor values) and writes the next brief, so the panel evolves from wherever it went. In the *anchored* regime, questions written from another transcript are reused, which steers every panel through the same conversation and separates the moderator’s contribution from the seats’ (Section 3.8).

### 2.4 Layered verification: quotation checks and source review

Verification has two layers (Figure 1C; Supplementary Methods S5). The first is deterministic. A citation passes when the cited document lies within the seat’s permitted set and the quoted span is found in a single stored passage of that document. The check rejects nonexistent identifiers, unauthorized sources, spans that are too short, and unmatched text. “Exact” means exact after normalization, not byte-identical to PDF typography. A similar membership test for verbatim spans has served as a training reward that increases quoting [93]. Here it is an inference-time gate on each claim. Repair is limited to evidence the seat already has: it may correct a quotation, attach existing support, narrow the claim, or withdraw it, and both failure and repair stay in the record. A *citation case* (one claim–source–quote relationship) and a *claim* (which may carry several citations) have different denominators. *Claim-level quote grounding* means a claim has at least one citation and every attached citation passes; *citation verification* counts individual checks. Neither implies semantic entailment.

The second layer is semantic and is itself an agent judgment. After a turn is frozen, peer seats review claim–source cases with supplied context and return a verdict (support, overstatement, irrelevance, misreading, cannot tell) with a rationale. The target is a failure the verifier cannot see: a genuine quotation asked to carry more than it says, such as a multi-clause recommendation resting on a source that establishes one clause, or an association promoted to a mechanism. Outcomes depend on which cases are sampled, how much context is shown, and the rubric. When referees judge different cases, their verdict rates cannot be compared as strictness, and the overstatement rate does not bound a human-adjudicated rate. Peer review is a second reading, not ground truth, and its errors may correlate with the generators’ because model evaluators share their biases and favor their own model family [38,39].

### 2.5 Monitoring and mapping deliberation

Monitoring makes the trajectory of a discussion inspectable without asserting what it means. Four statistics are computed at every turn (Figure 1C; Supplementary Methods S6). *Supplied-evidence diversity* is the overlap coefficient between seats’ retrieved document sets. *Cited-evidence diversity* is the Jaccard distance between their cited sets. *Position separation* is a spectral effective rank [50] of a similarity matrix over seat-level claim text, scaled to the roster and related to the Vendi diversity measures [19]. It has been proposed independently as a diagnostic for collapse among role-prompted agents [94]. We use it as a signal, not a decision rule. *Stance movement* is a directional herding statistic over transitions into and out of agreement. A negative value means movement away from agreement, not better quality. We also record new-document intake, participation, abstention, and quotation-check outcomes. All are operational descriptors, not measures of truth.

The isolated first turn sets a within-run reference. Later turns are compared with retention bands adjusted for packet separation and for a lexical proxy of how contested the question is. The bands are engineering choices, not parameters learned from labeled panels. A permutation diagnostic compares observed separation with separation after randomly redistributing the same material among seats. A breached band makes an intervention ladder available: peer routing, differential retrieval, own-work reanchoring, anonymization, isolation, evidence-constrained assigned dissent, termination, and declaration of collapse. An intervention can mechanically move the statistic it targets, so an improved value afterward does not show restored independence. The sequential monitor accumulates likelihood-ratio-shaped evidence toward an action boundary, as in a sequential probability ratio test [51]. However, it has not been calibrated to a false-positive rate or power. We therefore call it a monitor, not a validated test. The ladder did not fire in any study panel, so we exercised it in dedicated stress runs (Appendices H and I).

Statistics summarize a turn but do not show where an idea came from or where it went. We therefore also render each run as an interactive *idea-flow map*, in which claims are clustered into ideas and linked across turns by the stances and withdrawals seats filed (Section 3.7; Supplementary Methods S9). Building the map calls no model: every node, edge, and position is a projection of the record. Ideas and themes are titled by short topic labels that a language model writes from each idea’s own claims, and every idea opens onto its verbatim claims and verified quotations.

### 2.6 From discussion to executable research

The seat abstraction transfers to computational work unchanged. A scientist-associated agent receives the same corpus, identity, and rubric, and is asked to implement an analysis instead of arguing a position. In a *computational panel*, each agent builds and runs a prediction pipeline under shared cohort, split, and metric interfaces (Figure 1D), so its choices of representation, aggregation, effective training-set size, operating point, and model selection become inspectable outputs. In round one, each seat works alone and nominates one configuration by mean internal performance. In round two, the highest-ranked seats read one another’s reports and revise within their own identity, a controlled analogue of structured discussion. In the final round, frozen nominations are scored on an external collection. The resulting leaderboard tests methodological differentiation under combined conditioning; it is neither an ablation of corpus effects nor a validation of any scientist’s behavior.

### 2.7 Evaluation questions and the base-model variation

We evaluate four questions that need different evidence. *Evidence differentiation* asks whether seats receive different material. It depends only on acquisition, attribution, retrieval, and packing, and low overlap shows different exposure, not independent evidence: two non-overlapping reviews may summarize the same experiment. *Quote validity* and *claim support* are answered by the two layers of Section 2.4. A grounded claim requires every attached citation to pass. *Deliberation dynamics* asks whether further discussion changes the record, through new-document intake, participation and abstention, stance movement, position separation, and the idea-flow maps. *Scientific utility* asks whether the workflow makes research better. The present studies do not answer it directly: they show that the workflow runs, describe its behavior, and test its dependence on the base model. We count concrete products, such as a new control, a claim revised in light of evidence, or an implemented analysis, but these do not necessarily show an advantage over a simpler alternative (Appendix E).

**Figure 3.**
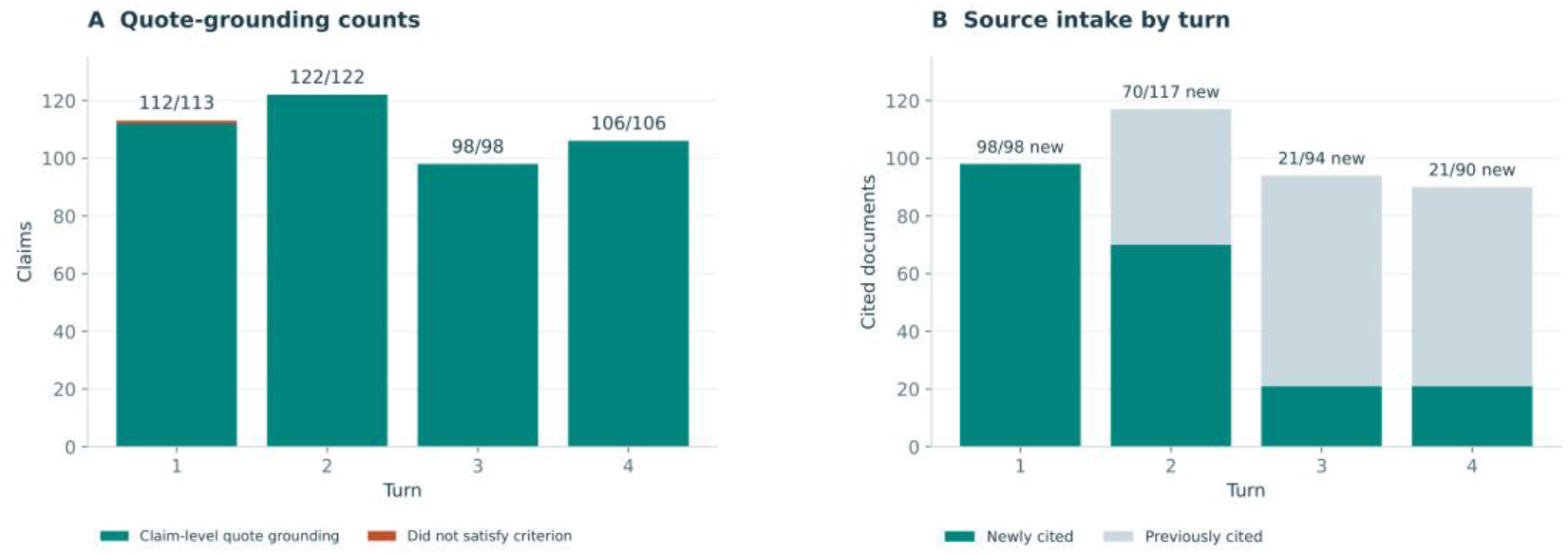
Primary-panel output remains quote-grounded while new-document intake declines. (A) Claims that meet the harness criterion and claims that do not; annotations give qualifying over total claims per turn. (B) Newly and previously cited documents within each turn. Document counts are not independent experiments, and claim-level grounding is not semantic support. The trajectories describe one run. Source: NBRT1 trajectory, ledgers, and per-turn reports; the anchored-run counterpart is `fig02_panel_dynamics_nb01`.

**Figure 4.**
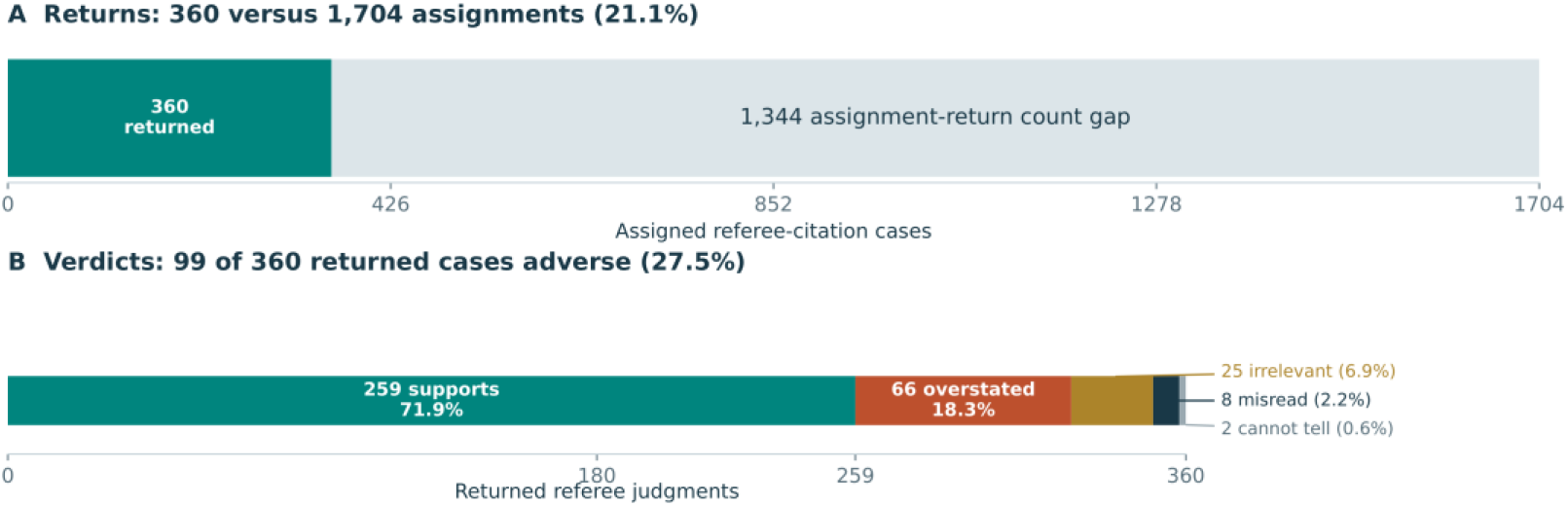
Audit coverage limits interpretation of the support judgments. (A) 360 returned judgments against 1,704 referee-citation assignments. The remaining 1,344 is a workload gap, not a count of uniquely missing assignments, because stable case IDs were not retained. (B) Verdicts among the 360 returned judgments: 259 support, 66 overstatement, 25 irrelevance, eight misreading, two cannot-tell. The returns are overlapping reviews of 34 claims, mainly by three authors, and are not established as representative. Source: NBRT1 turn-four audit assignments and responses; the anchored-run counterpart (385 of 2,080; 82 support, 297 overstated, six irrelevant) is `fig03_audit_coverage_nb01`.

Because the base model is part of every seat, we vary it with everything else fixed (Figure 1E). Six models (Opus 5, Sonnet 5, Haiku 4.5, GPT-5.6 Sol, GPT-5 mini, and Gemini 3.8 Flash), all at medium reasoning effort (Haiku 4.5 does not accept the setting), ran the primary roundtable from byte-identical first-turn packets in both regimes. The computational panel was repeated with each model, with the moderator and operator agent swapped to the same model and the data, partitions, and scoring code linked to the original run. This measures sensitivity to the model, not added value over simpler workflows. To compare architectures rather than models, we re-implemented the two closest published systems and ran them at fixed base model on our seats and questions: MPDS from its released code (Section 3.13, Appendix J), and MirrorMind, which has no released code [42], from its preprint at three base models (Section 3.14, Appendix K). Configurations, cohorts, and statistics are in Supplementary Methods S7 and S8.

## 3. Results

### 3.1 Seat policies produce differentiated evidence access

The inspected library states hold 32,488 documents and 921,895 indexed passages in the discipline-oriented implementation, and 42,016 documents and 1,148,025 passages in the scientist-associated implementation. They are distinct, overlapping archived states, so their counts must not be added. The twelve-seat discipline roster comprises nine disciplines, two disease domains, and a seat associated with vascular-normalization literature that has no single disciplinary label. The scientist-oriented roster has 25 seats: 14 scientist-associated seats, the nine disciplines, and the two disease domains (attribution counts in Appendix F).

An eight-seat retrieval diagnostic on a 28,214-document, 824,502-passage snapshot gave a mean overlap coefficient of 0.8% across 28 seat pairs, so the configured policies retrieve substantially different documents. By contrast, four role prompts that did not change the retrieval query returned the same 33 documents. Differentiation must enter the retrieval procedure to change its deterministic output.

### 3.2 A nine-seat neuroblastoma panel remained quote-grounded over four turns

The primary run, NBRT1, had four turns and nine seats: seven scientist-associated immunology seats (IMM1–IMM7) and the immunology and neuroblastoma discipline seats. Scientist-associated seats are reported, and referred to below, by codes (IMM*n* for immunology, CS*n* for computational scientists) that are stable across the paper, the appendices, and the supplementary examples. The researchers did not participate. All seats used Opus 5. A fixed operator-composed question opened turn one, and a moderator that read only this run’s record opened turns two to four. Section 3.8 applies the same procedure to every model.

The question concerned heterogeneity in the neuroblastoma immune landscape and its implications for experimental and trial design. Across 36 seat-turns, the panel filed 439 claims and 236 abstentions. Of the claims, 438 (99.8%) met the criterion that every attached citation verifies. Of 851 citation instances, 850 verified and one failed with `quote_not_found`; there were zero invented and zero out-of-scope citations (citation-level verification 99.9%). Neither percentage measures semantic support.

**Table 1.**
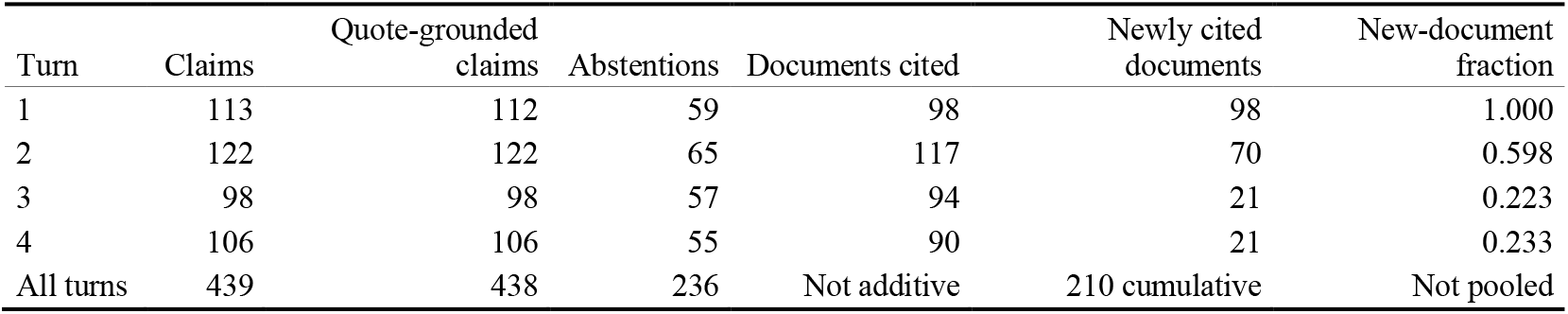
Output and evidence intake in the primary panel. Document counts are distinct verified-citation documents within each turn; newly cited documents exclude documents cited in earlier turns.

| Turn | Claims | Quote-grounded<br>claims | Abstentions | Documents cited | Newly cited<br>documents | New-document<br>fraction |
| --- | --- | --- | --- | --- | --- | --- |
| 1 | 113 | 112 | 59 | 98 | 98 | 1.000 |
| 2 | 122 | 122 | 65 | 117 | 70 | 0.598 |
| 3 | 98 | 98 | 57 | 94 | 21 | 0.223 |
| 4 | 106 | 106 | 55 | 90 | 21 | 0.233 |
| All turns | 439 | 438 | 236 | Not additive | 210 cumulative | Not pooled |

### 3.3 Seats withdrew inherited positions and narrowed the MYCN question to one design

Newly cited documents fell from 98 to 70, 21, and 21 across turns. At turn four, 21 of 90 cited documents (0.233) were new, and 210 distinct documents were cited over the run. The decline does not show that the literature was exhausted. Packet limits, accumulated disclosure, a narrowing question, and the role of a closing turn are alternative explanations. The herding statistic was 0.000, +0.273, and 0.000 after turn one, so movement toward agreement was recorded only at turn three.

The informative behavior was scope revision, elicited directly by the moderator’s second question. Turn one had been unanimous on four points, among them that MYCN amplification is an immune correlate in human disease, not a demonstrated causal determinant (T1: IMM2/C3, IMM4/C8, IMM5/C7, IMM1/C12, IMM7/C5, immunology/C5). The moderator then asked seats to argue the opposing side and to state whether each agreement was independent or inherited. The immunology seat withdrew the mechanistic half of its MYCN claim, because MYCN appeared in its evidence only as a risk classifier (immunology/C9). The neuroblastoma seat withdrew the basis of its claim, because the document it had relied on was absent from its current packet (neuroblastoma/C8). IMM1 withdrew its endorsement of the innate-effector strategy as inherited (IMM1/C3), while IMM4 declared its position independent and kept it (IMM4/C2). At turn three, IMM4 withdrew its use of du2026cells [54] and its load-bearing use of danilenko2025internationa [55] for a disease-wide reading of circulating tumor DNA (IMM4/C10, C11). IMM2 and IMM7 withdrew CSF-1R and CXCR1/2 inhibition from their trial queues on their own evidence (IMM2/C2, IMM7/C4). IMM4 agreed with IMM5, from its own systems work, that leukocyte density is the wrong in-situ quantity (IMM4/C8).

By the closing turn, the MYCN discussion had contracted to one design: compare MYCN-amplified and non-amplified regions from the same operation, tracking HLA class I and CD8 infiltration within patient (T4: IMM4/C10). IMM2 accepted it as the only MYCN comparison it would sign, and IMM5 required both regions on one section with one panel (IMM2/C9, IMM5/C12). Two seats rejected the moderator’s framing that no seat retained evidence for MYCN even as a correlate. The immunology seat held that its packet still supported the correlate (immunology/C13), and the neuroblastoma seat argued that the lack of an established correlate was a fact about the disease, not an artifact of its packet (neuroblastoma/C10). The panel thus closed with a shared design and an unresolved disagreement about what the existing evidence shows.

Asked whether the first randomization should intensify the existing anti-GD2 antibody with no new agent, all nine seats agreed from their own evidence. They differed on what to intensify (dose, schedule, or timing) and on which observation would refute the choice. Two seats declared their agent queues empty rather than fill them (IMM6/C2, IMM7/C3). This is agreement by elimination, and the record shows the eliminations. Clinical evidence for anti-GD2-containing therapy exists [20], but it does not validate an agent-generated extension. The example shows a panel recognizing the boundary between established evidence and a proposed extension. How often this happens, and whether it is clinically correct, has not been measured by blinded human review.

### 3.4 The monitor recorded a healthy trajectory and never fired

The primary run used the `tfidf-cosine` text backend. The isolated first turn set a position-separation reference of 0.77, with a floor of about 0.57 and a ceiling of about 0.93. Every follow-up turn returned a `healthy` verdict, no breached axis, and none as the applied move.

**Table 2.**
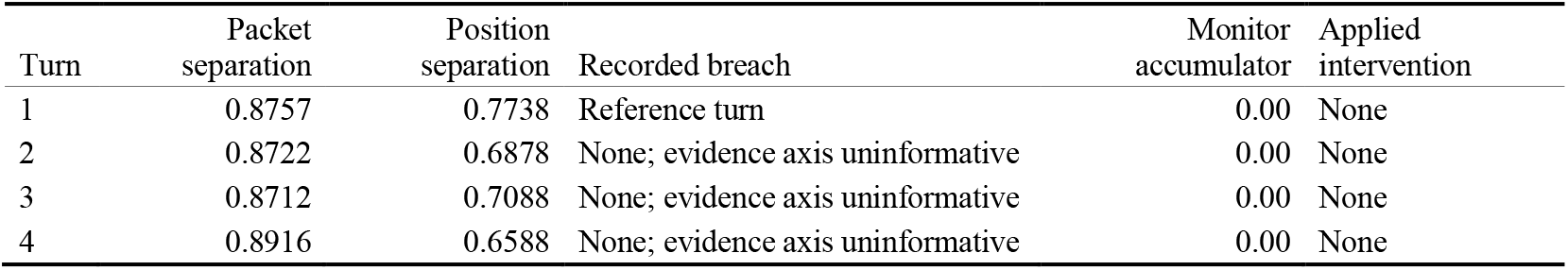
Recorded primary-panel monitoring trajectory. Values are operational scores for this run, not calibrated probabilities of collapse.

At every follow-up turn, the evidence-axis permutation null saturated, so that axis was excluded and the monitor ran on position separation alone. A saturated null means that observed separation did not exceed the permutation distribution, not that seats cited identical documents. Position separation stayed within 0.659–0.709, and the per-turn log-likelihood-ratio increments (−8.32, −9.56, and −6.98) all fell below the reset boundary.

### 3.5 Peer source review covered a narrow sample and mostly supported it

The turn-four audit covered 213 eligible citation instances from 106 grounded claims across 90 documents, each assigned to the eight nonauthor referees (1,704 referee-citation assignments). The archive holds 360 returned judgments (21.1% of the workload), exactly 40 per referee: 259 `supports, 66 overstated, 25 irrelevant, eight misread`, and `two cannot_tell`. Of these, 99 (27.5%) are adverse.

**Table 3.**
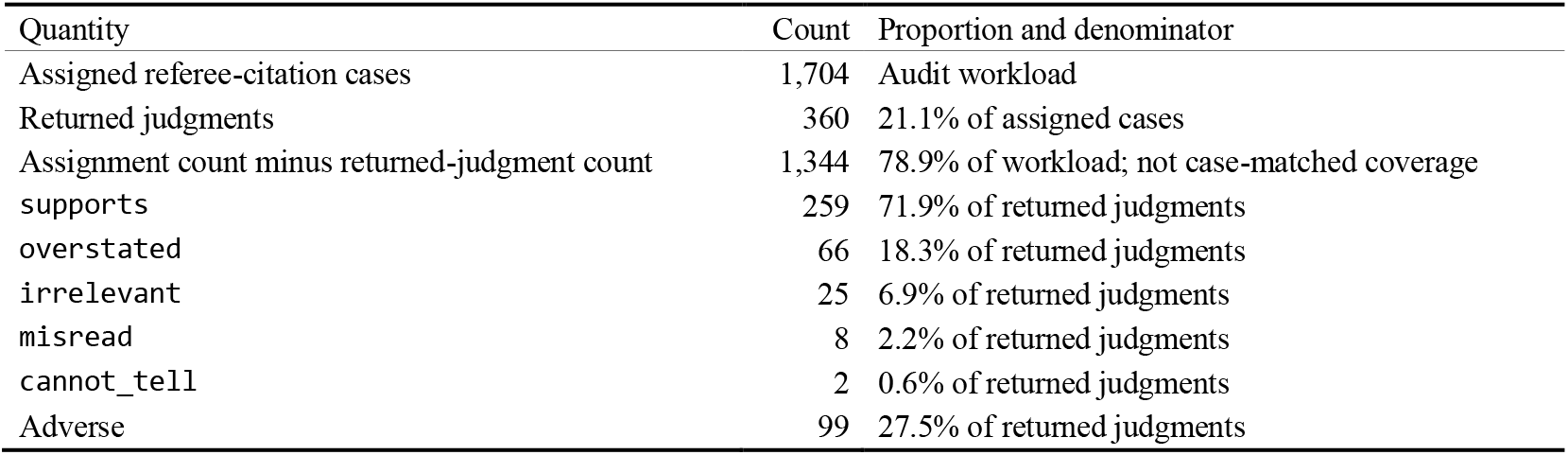
Final-turn audit coverage and returned outcomes. Assignments and returned judgments are different denominators. Adverse verdicts are `overstated, irrelevant, or misread`.

| Quantity | Count | Proportion and denominator |
| --- | --- | --- |
| Assigned referee-citation cases | 1,704 | Audit workload |
| Returned judgments | 360 | 21.1% of assigned cases |
| Assignment count minus returned-judgment count | 1,344 | 78.9% of workload; not case-matched coverage |
| supports | 259 | 71.9% of returned judgments |
| overstated | 66 | 18.3% of returned judgments |
| irrelevant | 25 | 6.9% of returned judgments |
| misread | 8 | 2.2% of returned judgments |
| cannot_tell | 2 | 0.6% of returned judgments |
| Adverse | 99 | 27.5% of returned judgments |

Coverage was narrow and nonrandom. The 360 returns concern 34 of 106 grounded claims and 29 of 90 documents, and 354 concern claims by the first three authors in packet order. The uniform 40 returns per referee reflect seats working down the case list and stopping at a round number, not truncation. Yet, the instructions set no case limit and the harness imposes no output cap.

The review layer nonetheless did what quotation matching cannot: it located where a source’s reach stopped short of the claim. IMM2’s closing claim that no myeloid-directed agent in this class was useful without a partner cited two papers from its permitted corpus. For `holmgaard2016oncoimmunolo` [56], which reports that CSF-1R blockade alone neither increased T-cell infiltration nor delayed growth, six of eight referees returned supports. For `holmgaard2016ebiomedicine` [57], which reports single-agent activity in MDSC-high tumors, seven of eight returned overstated or misread and quoted the contradicting passage (T4: IMM2/C1). The referees narrowed the wording without disputing the recommendation. Conversely, IMM4’s within-patient MYCN design cited a mouse microbiota study [58] for the premise that hosts of one genotype differ in immune composition with environment. Referees returned supports because they judged the premise, not the design (IMM4/C10). A rubric that scores empirical premises, inferential links, and proposed experiments separately would make these distinctions explicit (Appendix E).

### 3.6 A ten-round panel sustained differentiated inquiry

A separate six-seat roundtable (NBRT18) completed ten turns without observed conversational collapse. It kept introducing evidence and specifying different experiments through its closing turn, filing 436 claims, 426 of them quote-grounded, and citing 213 distinct documents. The seats agreed that the original pediatric treatment proposal was not trial-ready, yet they still differed at the end on the evidence required to advance it. Appendix D gives the trajectory, scientific progression, source audit, and the adaptive protocol under which the result was obtained.

### 3.7 Idea-flow maps make the course of a panel discussion inspectable

The statistics above summarize each turn, and a transcript lists what each seat said, but neither shows how positions met, merged, split, or were abandoned. Argument-mapping systems capture this by asking participants to file each contribution into a typed graph as they speak [99,100], and bipolar argumentation formalizes support alongside attack [101]. Visual analytics follows evolving topics as streams that merge and split [102–104], and recent tools display the event logs of LLM agents [105]. An ExpertTwins record already holds what such a map needs: every claim carries its seat, turn, and verified quotations, and every stance names a target seat and a verdict. The map is therefore drawn after the fact, without changing how the panel runs.

Our renderer turns one run record into a self-contained interactive HTML page (Supplementary Methods S9). Claims are clustered by semantic similarity into *ideas* within each turn, and into *themes*, each with a fixed color, across the run. Turns are columns, ideas are boxes titled by a short topic label, and seats are roots on the left. Edges join an idea to its uptake in the next turn and are colored by the dominant stance: green for support, red for contradiction, purple for mixed, amber for a request for other evidence, dashed black for a withdrawal, and dotted gray for a continuation inferred from similarity alone. Bars under each idea show which seats voiced it, and their borders show how each seat received it in the next turn. Broadly voiced, supported ideas are bold; withdrawn or contradicted ones are pale. Hovering highlights an idea’s lineage, and clicking opens its claims, quotations, cited documents, peer-review verdicts, and reception. The structure of the map involves no model call, and its only added judgment is the clustering. Topic titles and one-sentence gists are written by a language model from each idea’s claims alone and checked automatically. The idea’s most central claim is shown verbatim beside them (S9).

#### The map recovers the narrative of Section 3.3 as structure

The counts between columns alone show the panel moving from assent to contest. Out of turn one, the map drew 112 support links and 7 contradiction or mixed links; out of turn two, it drew 95 and 39. The stances beneath these links moved the same way: contradicting stances rose from 9 of 862 resolved stances on turn-one claims to 74 of 1,540 on turn-two claims. Over the run, 28 claims announced a withdrawal. Pinning one idea shows the events behind such counts (Figure 5B). The pinned turn-three idea, on what archival imaging can show of fresh myeloid function, has as its central claim IMM4’s withdrawal of a single-cell source [54] and also holds IMM7’s withdrawal of CXCR1/2 inhibition (Section 3.3). It descends by support from the immunology seat’s turn-two withdrawal of the mechanistic half of its MYCN claim, and by withdrawal edges from the turn-two positions it retracts. In turn four, six seats received it with agreement and IMM5 with mixed stances. Downstream, it feeds a cluster in which seats name the only MYCN comparison they will accept, a within-patient one. It also draws one contradiction, from a turn-four idea disputing IMM7’s view that an archival multiplex panel cannot see the granulocytic compartment. A reader can start from the narrowed design of turn four, walk upstream to the withdrawals that produced it, and descend from any idea to the quotations it rests on. **The map shows a collapse that grounding and participation statistics score as healthy.** Figure 6 draws two stress runs from Appendix H identically. In NBRT13, a negative control under neutral moderation, ideas branch, cross themes, and draw contradiction as well as support. In NBRT14, a moderator prompt dictated six claims verbatim from turn three onward, and the map turns into six parallel lanes. Every later turn holds the same six ideas, each voiced by all four seats and linked only to its own predecessor, with no contradiction after turn two and no stance at all in the last two transitions. Grounding stayed at 100% and participation entropy at its maximum throughout NBRT14. Therefore, neither statistic separates the two panels, but the map separates them at a glance. Position separation did detect this collapse, and the monitor escalated (Appendix H). The map adds what a statistic cannot: which ideas were frozen, and that every seat voiced them. Deference under asymmetric seat power (Appendix I) leaves a subtler mark, in the composition of ideas rather than the shape of the graph (Appendix L.5). Appendix L documents the viewer and maps four further runs.

**Figure 5.**
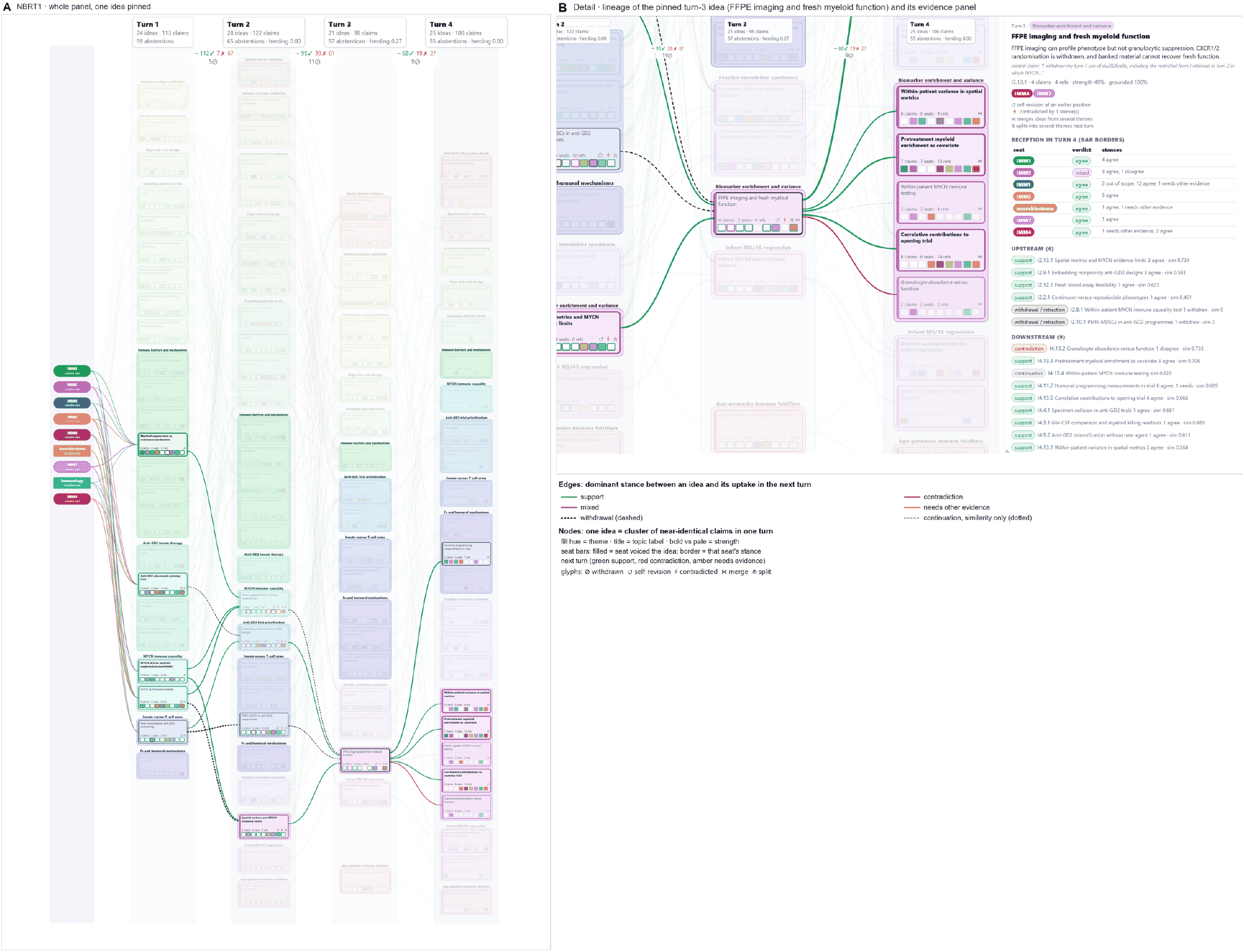
The primary panel as an idea-flow map. (A) NBRT1, nine seats over four turns: 98 ideas in 16 themes. Seats are the colored roots on the left, and each turn is a column. The counts between columns give the links drawn out of the previous turn at the default filter (support, contradiction or mixed, and needs other evidence) and the withdrawal links. One turn-three idea is pinned, so only its lineage stays at full opacity. (B) Detail of the pinned idea, whose central claim is IMM4’s withdrawal of its turn-one use of a single-cell source [54], with its six upstream and nine downstream ideas and the evidence panel. Bar borders give the turn-four reception: six seats agreed and IMM5 was mixed. Seats are shown by code; the viewer’s tags marking a cited document as a seat’s own work are hidden to preserve anonymization.

**Figure 6.**
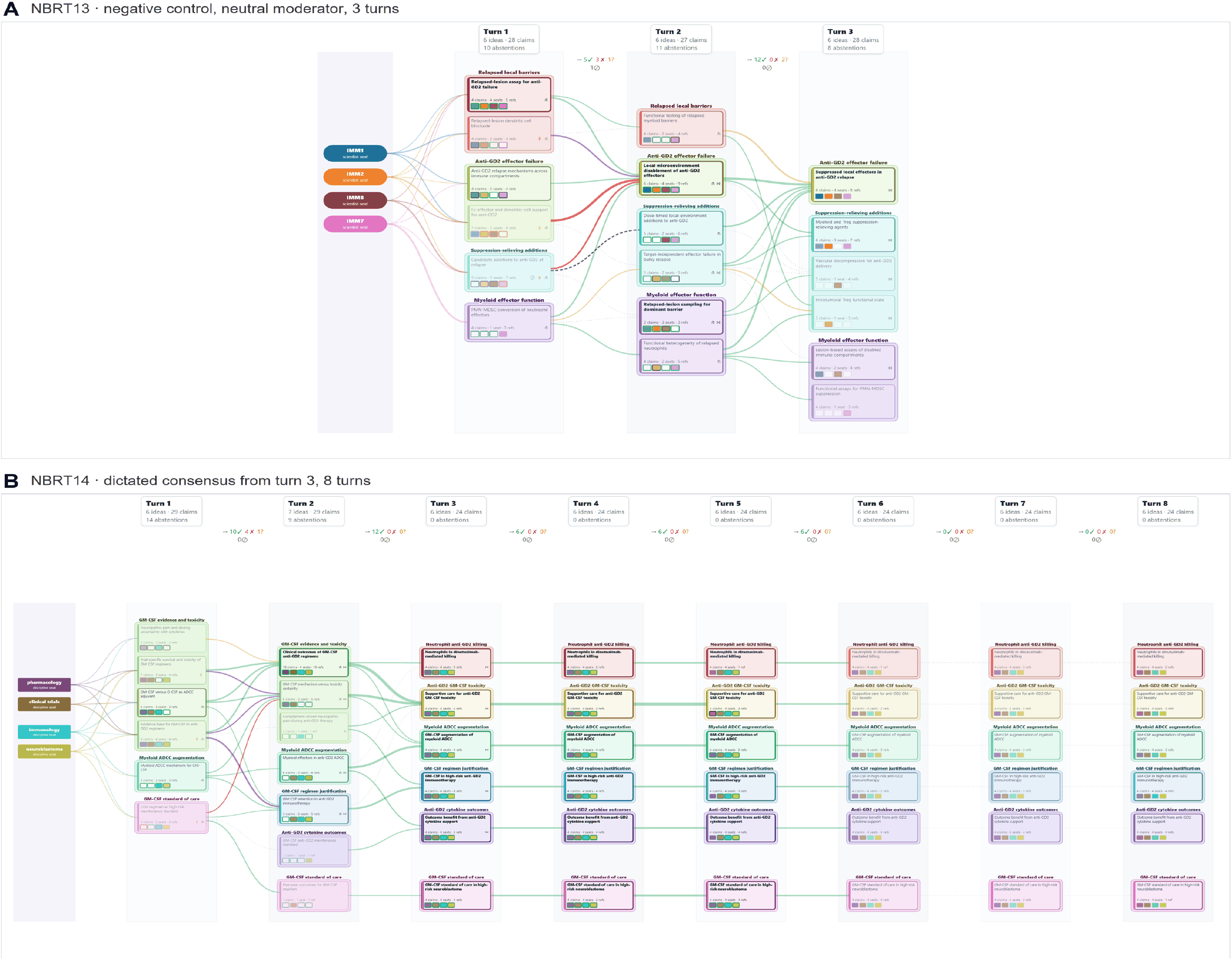
Free and dictated discussion, drawn identically. (A) NBRT13, the negative control: four scientist-associated seats, neutral moderation, three turns. (B) NBRT14: four discipline seats over eight turns, with a turn-three moderator prompt that dictated six claims verbatim and was reused through turn eight (Appendix H). From turn three the ideas form six parallel lanes, each voiced by all four seats, with no contradiction after turn two and no stances in the last two transitions. Ideas from turn six onward are pale because they drew no stances. Encodings as in Figure 5.

**Figure 7.**
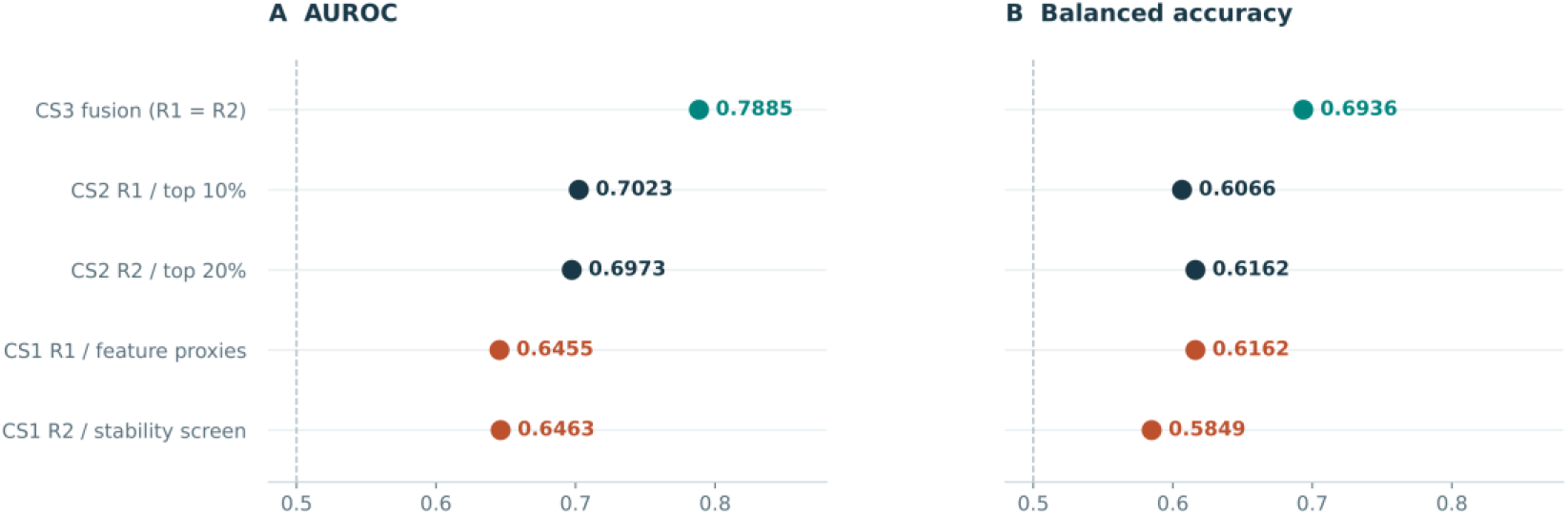
Full-cohort external results use a consistent evaluation unit. Each row is one distinct nominated configuration on the same 75 slides. (A) AUROC. (B) Balanced accuracy at internally derived thresholds. CS3’s duplicate entries are collapsed, and CS1’s first-round feature-proxy procedure is labeled explicitly. Points are descriptive estimates without patient-population confidence intervals. The reported baseline is omitted because its exact prediction artifact was not reconciled; the dashed 0.5 references are not significance thresholds. Source: the external comparison and saved-prediction records from aimycn-01 and its second-round extension (Appendix C).

### 3.8 Two arms separate the model’s contribution from the moderator’s

We next held the question, seats, and initial evidence fixed and changed only the language model. Six models (Opus 5, Sonnet 5, Haiku 4.5, GPT-5.6 Sol, GPT-5 mini, and Gemini 3.8 Flash), all at the provider’s medium reasoning-effort setting (Haiku 4.5 does not accept it), used the nine-seat roster, retrieval settings, four-turn structure, verification rule, and final-turn source review of Section 3.2. All nine first-turn packet files are byte-identical across runs, which establishes identical instructions and evidence, not merely a shared corpus path.

We ran the comparison in two arms. In the **primary (endogenous) arm**, turns two to four were opened by a moderator that read only that run’s own record (verified claims, abstentions, disclosed sources, citation-check verdicts, and monitor values) and wrote a fresh 1,300–1,700-word followup. The seat model is then the only variable, and each conversation evolves from wherever its panel went. In the **anchored arm**, every panel received the same four follow-up questions. The operator composed them from the Opus transcript of a separate run, including that panel’s summaries, audit counts, and monitor values. Opus is therefore the only model whose anchored follow-ups describe its own conversation; for the other five, they describe a conversation their seats never had. We keep the anchored arm as a supplementary analysis because it answers a legitimate question: how do models differ when steered through the same conversation? Appendix G documents both arms’ controls and the anchored-arm thematic analysis.

### 3.9 Model capability determined whether evidence survived the workflow

All six primary-arm runs reached four ingested turns: 216 planned seat-turns, 1,259 parsed claims, and 1,917 parsed citation instances, of which 1,852 verified. Opus, Sonnet, Sol, and Gemini kept high quotation grounding on every turn and near-full participation. Haiku and mini did not, and they failed differently (Table 4).

**Table 4.** Final archived output in the primary arm. A grounded claim has at least one citation and every attached citation verifies. Abstention entries are on-the-record declinations to claim on a topic and are distinct from seat-turns with no output.

| Reported model | Parsed claims | Quote-grounded claims | Claim-producing seat-turns / 36 | Abstention entries | Verified citation instances / parsed instances | Distinct documents with a verified citation |
| --- | --- | --- | --- | --- | --- | --- |
| Opus 5 | 439 | 438 (99.8%) | 36 | 236 | 850 / 851 | 210 |
| Sonnet 5 | 241 | 236 (97.9%) | 36 | 187 | 334 / 339 | 122 |
| Haiku 4.5 | 86 | 32 (37.2%) | 29 | 162 | 57 / 104 | 37 |
| GPT-5.6 Sol | 202 | 201 (99.5%) | 32 | 211 | 292 / 293 | 87 |
| GPT-5 mini | 30 | 29 (96.7%) | 13 | 30 | 30 / 31 | 16 |
| Gemini 3.8 Flash | 261 | 256 (98.1%) | 35 | 185 | 289 / 299 | 115 |

Grounding did not decay with conversational depth. The four high-grounding models held 91–100% on every turn, and none declined monotonically (Appendix G, Table G3). **Haiku failed by paraphrase.** Its claims are structurally valid and cite real, topically appropriate documents, but 40 of 104 recognized citation instances quoted text absent from the source, 15 claims carried no recognized citation list, and its grounded fraction fell from 60% at turn one to 9% at turn four. The transcript nonetheless reads as a confident, densely cited discussion. **GPT-5 mini failed by absence**: only 13 of 36 seat-turns produced any claim. Two Haiku instances and one mini instance cited identifiers absent from the permitted corpus.

Among the four functioning models, the base model alone produced a 2.2-fold range in claim yield (202–439), a 2.4-fold range in distinct verified documents (87–210), and a three-fold range in the turn-level disclosure-check adverse share (20% Gemini, 41% Opus, 59% Sonnet, 63% Sol; Appendix G, Table G4).

### 3.10 Conversations diverged after turn one, and agreement fell with them

Because each primary-arm moderator wrote from its own run’s record, the six conversations went to different places (Table 5).

**Table 5.** Closing-turn threads raised by the moderator in each primary-arm run. Condensed from the section headings of each run’s turn-four follow-up (`runs\<RUN>\moderator\q-04.md`). The moderator model is Opus 5 throughout; only the record it read differs.

| Run | Closing-turn threads |
| --- | --- |
| Opus 5 | The trial the panel agreed to open contains no new agent; the spatial axis has become a declared null, and whether that is honest or an exit; the agreed measurement may not be measurable in the agreed trial; the panel’s own evidentiary rules may forbid the trials it proposed; three threads that arrived late. |
| Sonnet 5 | Seven incompatible experiments proposed for one MYCN claim; the measurement missing under all of them; the axes no seat could answer; a specimen collision between imaging and functional assays; what is still hanging. |
| GPT-5.6 Sol | A survivorship audit of turn-one and turn-two claims; circularity in the trial ranking; the innate bet and a GD2-low tripwire; a loss function no seat will own; the three shared abstentions, classified; MYCN as a rule a seat could sign; a stress test of the one thing everyone agreed to. |
| Gemini 3.8 Flash | Innate primacy: the panel’s own data contradict its consensus; an IL-2 and GM-CSF contradiction the panel walked past; a unanimous conclusion built on challenged foreign evidence; GD2 loss under three incompatible readings with no assay; a definition that cannot be executed in children. |
| Haiku 4.5 | The rate-limiting barrier as the one live disagreement, and whether it is decidable; the gap three seats named and nobody filled; claims that took adverse citation checks; three axes on which every seat abstained every turn; where the panel never went. |
| GPT-5 mini | Why the table went silent, and whether the silence is a finding or a failure; the two live contested positions and the adverse checks against them; MYCN, and the one abstention that did its job. |

To measure divergence, we matched claims across runs on a shared cited document plus lexical similarity, or on high lexical similarity alone (Appendix G.3). We restricted the analysis to the three models that completed the protocol with full grounding (Opus, Sonnet, and Sol), so that weak-model failures could not drive the result. Agreement concentrated where inputs were identical (Table 6).

**Table 6.** Strong-model claims with a counterpart in another strong model’s run, by turn and arm. Turn-one responses are shared between arms. Pooled figures use a matching model fitted jointly across both arms; per-turn figures use arm-specific fits and differ from the pooled values by at most one point.

|  | Turn 1 | Turn 2 | Turn 3 | Turn 4 | Pooled | Unshared claims grounded | Documents cited by all three | Documents cited, all six runs |
| --- | --- | --- | --- | --- | --- | --- | --- | --- |
| Primary (follow-ups from own record) | 31% | 9% | 6% | 5% | 11% (96 / 882) | 99.1% | 41 | 299 |
| Anchored (follow-ups held fixed) | 28% | 30% | 33% | 20% | 26% (265 / 1,012) | 97.7% | 59 | 282 |

Three results follow. **First, most of what a strong model says under the protocol, no other strong model says.** In the primary arm, 11% of strong-model claims had a counterpart elsewhere, and 63% of those shared claims were filed at turn one, although turn one supplied only 27% of claims. Five claim clusters were shared by all three strong models and 33 by two. Across all six runs, 11 clusters reached four or more models (7 reached four and 4 reached five), and none reached all six. **Second, replication perfectly predicted passing the quotation check, but few claims replicated.** Every shared strong-model claim verified, and all seven ungrounded claims in the strong-model subset were unshared. **Third, after turn one, the moderator’s questions, more than the model, set the level of agreement.** Under fixed follow-ups, agreement stayed at 28–33% through turn three and fell only at the closing turn. Under endogenous follow-ups, it reached single digits by turn three. The control is each model against its own run in the other arm: when only the follow-ups changed, Opus reproduced 29–35% of its own claims, Sonnet 42–51%, and Sol 30–34% (Appendix G, Table G4).

### 3.11 Shared conclusions were restraints; distinctive developments and repetition differed by model

The claims replicated across all three strong models in the primary arm form a short list, all from turn one: (1) MYCN amplification is an immune correlate whose causal role in human disease is not established [59,60]; (2) a marker-defined myeloid population may not be called MDSC without demonstrated suppressive function [61]; (3) the “cold” description of neuroblastoma is directionally correct but incomplete [62]; (4) the association of MYCN and 11q with an M2-skewed macrophage phenotype is correlative [63]; and (5) PMN-MDSC ferroptosis is ready for neuroblastoma-specific functional testing, not clinical translation [64,65]. Each marks a boundary of what the evidence shows. The shared core is a set of restraints, not of mechanisms.

What each model built on that core differed, as one shared source shows. The CSF-1R/CTLA-4 scheduling result (`holmgaard2016oncoimmunolo`) [56] was claimed in four of six runs. Opus’s IMM2 turned it into a requirement that any myeloid agent added to an anti-GD2 backbone be randomized across schedule, not simply added (NBRT1, T2/IMM2/C1). Sonnet’s IMM2 connected it to the fact that the high-risk maintenance block delivers antibody, cytokine, and differentiating agent together [66,67] (NBRT2, T2/IMM2/C3). Gemini’s filed a near-verbatim restatement of the source sentence at turns one through four (NBRT6, T1–T4/IMM2), and Haiku’s restated it at turn three without a citation. Re-filing a claim the seat had already made (same seat, later turn, lexical similarity ≥ 0.80) accounted for 14 of Gemini’s 261 claims, 4 of Haiku’s 86, 4 of mini’s 30, and none of the claims of Opus, Sonnet, or Sol. Raw claim counts therefore overstate Gemini’s contribution relative to the strong three.

Justified revision also occurred without human steering. Gemini’s neuroblastoma seat asserted at turn one, on an association quotation [59], that MYCN amplification “drives” a cold microenvironment (NBRT6, `T1/neuroblastoma/C7`). The run’s own moderator raised the causal-versus-correlative question at turns two and three, and at turn three the seats corrected their own turn-one phrasing. The neuroblastoma seat restated the relationship as correlative (T3/neuroblastoma/C3), and IMM5 formally withdrew its causal phrasing on the CCR2/CCL2 axis (T3/IMM5/C1). Sol’s IMM3 withdrew vaccine readiness in favor of an ex-vivo gate, citing dendritic-cell-targeting vaccine studies [68,69] (NBRT4, T3/IMM3/C4). Distinctive unshared contributions included Opus’s paired quantity and per-cell suppression readout [57,70] (T2/IMM2/C2), Sonnet’s innate-arm ceiling signature [71–73] (T2/IMM2/C1), and Sol’s requirement that MYCN causality be shown under inducible perturbation in three models [57,74] (T2/IMM2/C4). Locators and sources are in Appendix G, Table G7.

### 3.12 Model-dependent participation changes the meaning of convergence

No intervention ladder fired in any of the twelve base-model runs. That is a property of these runs, not of the instrument. In the induced-collapse run of Appendix H (Figure 6B), the ladder detected the breach and escalated through five interventions before halting the panel. In the primary arm, the moderator’s guardrail verdict was healthy at turns three and four for Opus, Sonnet, Sol, and Gemini. It was undetermined for Haiku at turns three and four and for mini at every turn, because too few grounded contributions remained to evaluate a reference band. In the anchored arm, the participation guardrail fired on Haiku at every turn and on mini at turns two and three. Both instruments flagged exactly the two degraded runs, and neither could repair them.

Among the strong models, position separation declined with depth in the primary arm (Opus 0.77 to 0.66, Sonnet 0.86 to 0.76, Sol 0.86 to 0.69), while the newly cited fraction fell to 0.18–0.37 by turn four. The seats were engaging, converging, and consolidating a shared evidence base. Gemini’s separation rose (0.85 to 0.90) despite its low adverse volume. Haiku and mini sat at 0.92–1.00 on every turn and drew 80–100% new documents at turn four. This is incoherence, not viewpoint diversity: seats restarted from fresh evidence without engaging one another. A diversity metric applied without grounding and contestation covariates would rank the two failed runs as the most intellectually diverse of the six. A usable panel must preserve enough well-structured, source-traceable contributions for exchange to occur. And if a result is to be attributed to the model, the questions that drive the exchange must come from the panel’s own record.

### 3.13 A head-to-head comparison with multi-persona debate separates substrate, architecture, and claim unit

The base-model study varied the model within one architecture. Here we hold the model fixed and vary the architecture. The closest published alternative is MPDS [24]. Its personas argue in a round robin over a retrieved paper set, and a turn is accepted only if it carries at least three distinct identifiers that exist in that persona’s database. Our harness imports the authors’ prompt construction and turn validation from their released code [98] and extends the debate from two personas to nine, to match our panel. Four arms received the byte-identical opening question of NBRT7 and ran on Opus 5 at medium reasoning effort. MPDS1 runs the published system faithfully on its own abstract retrieval. MPDS2 runs the same architecture on our corpus, which separates the effect of the evidence substrate from that of the architecture. NBRT19 runs ExpertTwins with the same nine discipline seats, spanning T-cell, myeloid, antigen-presentation, general immunology, and neuroblastoma expertise (Appendix J.1). We used discipline seats because MPDS personas are disciplinary. NBRT7 runs ExpertTwins with nine scientist-associated seats, so it differs from NBRT19 in seat construction, not protocol. Both debate arms completed 27 turns (nine seats over three rounds) and a final judge synthesis, and all 54 debate turns passed the published validator on the first attempt.

#### Identifier validation is not quotation validation

Both debate arms passed their own validator perfectly: none of their 1,371 and 1,160 citation instances pointed to a record absent from the database. But neither architecture attaches source text to a claim, so none of their 852 and 745 cited sentences can be checked against what the source says. The two ExpertTwins arms attach 832 and 920 verified quotations that cover all of their claims. The debate architecture also has no abstention path, so a persona with no relevant evidence must still produce three pointers. On the other hand, the ExpertTwins arms recorded 293 declinations in NBRT19 (40.5% of seat-question opportunities) and 163 in NBRT7 (27.3%). These are differences in what can be audited, not in measured accuracy.

#### The evidence substrate and the architecture each account for part of the on-topic gap

The share of distinct cited sources that are neuroblastoma-specific rises from 6.9% (MPDS1) to 21.9% (MPDS2), 40.8% (NBRT19), and 45.4% (NBRT7). MPDS1’s first speaker named the problem itself: its snapshot “is a general TIL literature, not a neuroblastoma literature” and “contains exactly two neuroblastoma-specific records”. Changing only the corpus adds 15.0 points, roughly two-fifths of the gap to our runs, and the downstream text changes with it, from a 25.5 KB generic judge synthesis in MPDS1 to a 42.3 KB neuroblastoma-specific one in MPDS2. Changing the architecture on the same corpus and seats adds another 18.9 points (MPDS2 to NBRT19). Here, the architecture includes how each seat’s evidence is selected from the corpus. Changing from discipline to scientist-associated seats adds 4.6 more. A comparison that varied corpus and architecture together could not tell these contributions apart.

**Table 7.** Head-to-head comparison at fixed base model (Opus 5, medium effort) and identical initial question. MPDS1 and MPDS2 emit cited sentences; NBRT19 and NBRT7 emit claims with attached citations, whose citation-instance counts are verified quotations. Adjudicated support is the blinded set-level scoring described below and in Appendix J; the permissive and strict rubrics scored identical sets against identical, complete source text.

| Property | MPDS1 | MPDS2 | NBRT19 | NBRT7 |
| --- | --- | --- | --- | --- |
| Architecture | Debate [24] | Debate [24] | ExpertTwins | ExpertTwins |
| Evidence substrate | OpenAlex abstracts | Our corpus | Our corpus | Our corpus |
| Seats | 9 disciplinary | 9 disciplinary | 9 disciplinary | 9 scientist-associated |
| Seat-turns | 27 | 27 | 36 | 36 |
| Claims or cited sentences | 852 | 745 | 431 | 434 |
| Citation instances | 1,371 | 1,160 | 832 | 920 |
| Distinct cited sources | 635 | 494 | 238 | 183 |
| Neuroblastoma-specific sources | 6.9% | 21.9% | 40.8% | 45.4% |
| Pointers to absent records | 0 | 0 | not applicable | not applicable |
| Quote-verifiable claims | 0% | 0% | 100% | 100% |
| Abstentions | none possible | none possible | 293 | 163 |
| Adjudicated support, permissive | 72.9% | 75.7% | 58.6% | 47.1% |
| Adjudicated support, strict | 67.1% | 71.4% | 32.9% | 15.7% |
| Normative or design claims | 7.5% | 7.0% | 41.1% | 57.4% |

#### Blinded adjudication of claim support measures the claim unit as much as the evidence

We sampled 70 claim–source sets per arm (280 in total), stratified by seat, removed arm-identifying markers, and had Opus 5 adjudicators score each set against complete source text under two rubrics (Appendix J). Under the permissive rubric (does any source bear on the claim?), support was 72.9%, 75.7%, 58.6%, and 47.1%. Under the strict rubric (does the set establish every component of the claim?), it was 67.1%, 71.4%, 32.9%, and 15.7% for MPDS1, MPDS2, NBRT19, and NBRT7, respectively. The strict rubric cost the debate arms 5.7 and 4.3 points (4 and 3 of 70 sets) and the ExpertTwins arms 25.7 and 31.4 (18 and 22 sets). This asymmetry reflects the claim unit more than the evidence. A debate turn emits cited descriptive sentences averaging 1.83 finite verbs, whereas NBRT19’s seats emit compound positions averaging 3.26. Moreover, 41.1% of NBRT19’s claims and 57.4% of NBRT7’s are normative or design statements (proposed trials, triage rules, and funding priorities), against 7.5% and 7.0% in the debate arms. No literature source can establish a proposed trial, so the strict rubric scores as unsupported a behavior the framework is built to produce (Section 2.6). Restricted to descriptive empirical statements of at most 60 words, support was 75.2% (95% CI 66.5 to 82.3) for the pooled debate arms and 54.8% (37.8 to 70.8) for the pooled ExpertTwins arms. The intervals overlap, and this sample does not resolve the residual difference. The support rates therefore describe what each architecture emits; they do not rank grounding quality.

#### Seat construction leaves its own mark on the record

With question and model fixed, NBRT19 placed 165 of 431 claims outside its seats’ home territory, against 23 of 434 in NBRT7, whose scientist-associated seats stayed closer to their publication records. NBRT7 was correspondingly more speculative (40.3% novel syntheses against 14.8%) and NBRT19 more descriptive (43.9% against 30.4%). The monitor separated them too. Position separation stayed between 0.73 and 0.81 in all four turns of NBRT19 but fell to 0.51 twice in NBRT7, and new-document intake fell below the 0.15 convention at turn four in NBRT7 (0.116) while NBRT19 stayed just above it (0.156). Neither run triggered an intervention. The idea-flow maps show the same contrast as shape: a few broad currents in NBRT7 and many narrower, contested ones in NBRT19 (Appendix L.4). The debate arms expose no comparable trajectory, because they record no abstentions, no stances, and no per-turn monitoring. Against the closest published alternative, then, ExpertTwins retrieved more on-topic material and made more of its record checkable. Whether its claims are better supported, once the claim unit is matched, remains open.

### 3.14 A comparison with author-memory fusion measures what a single answer keeps from its specialists

MPDS differentiates agents by discipline. MirrorMind [42] is the published system closest to our scientist-associated seats. Each author agent holds an episodic memory of the author’s papers, a semantic memory of how the work developed, and a persona schema. A coordinator decomposes a query and routes sub-tasks to author and domain agents. A fact checker and a consistency checker review their outputs, and a knowledge integrator fuses everything into one answer. No code has been released, so we re-implemented the pipeline from the preprint (Appendix K).

We matched everything but the architecture. MirrorMind answers a question once instead of deliberating over turns, so we gave it the questions the ExpertTwins moderator had actually asked: the shared turn-one question and each run’s own turn-two to turn-four questions. Its nine agents were built over the NBRT1 roster from our corpus: The seven scientist-associated seats from their attributed publications and the two discipline seats from field corpora. Each MirrorMind stack ran wholly on Opus 5 (MM1), Sonnet 5 (MM2), or GPT-5.6 Sol (MM3) at medium reasoning effort and is compared with the ExpertTwins run on the same model (NBRT1, NBRT2, NBRT4). The moderator’s follow-up questions refer to a transcript MirrorMind has not seen, so a context-augmented control supplied the prior-turn record with the turn-two question (MM1c, MM2c, MM3c), a concession in MirrorMind’s favor. All 15 runs completed: four single-shot queries in each of the three arms, and the three context-augmented queries.

**Table 8.**
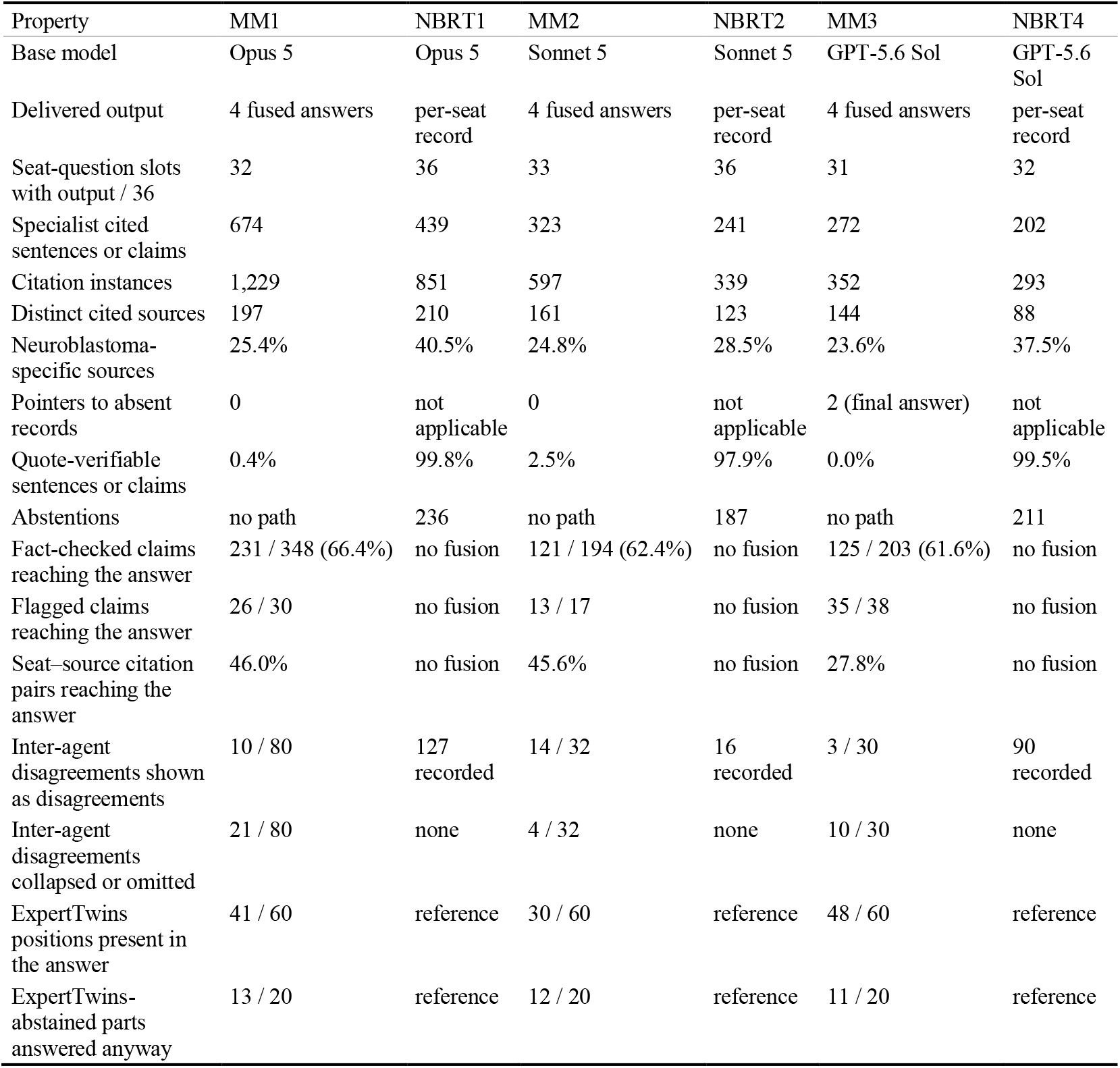
MirrorMind re-implementation against ExpertTwins at matched base model, roster, corpus, and questions. Each column pair sums four queries (MirrorMind) or four turns (ExpertTwins). MirrorMind specialists emit cited sentences. ExpertTwins seats emit claims with attached quotations. Survival rows are Opus 5 judgments of the final answer against the fact checker’s claim list and the consistency checker’s disagreement list. Content rows compare the final answer with the ExpertTwins per-seat record (Appendix K). Quote-verifiable means a quoted span of at least 40 characters found, after normalization, in the evidence retrieved for that seat (MirrorMind), or a claim whose attached quotations all pass verification (ExpertTwins, Table 4). Distinct cited sources count every cited document, so ExpertTwins values can exceed Table 4’s count of documents with a verified citation. In the last two rows, the ExpertTwins record is the reference against which each MirrorMind answer was judged.

#### Context management is part of the architecture

MirrorMind’s fact checker cross-references every specialist’s output against that specialist’s evidence in a single prompt. On nine-seat queries, that prompt reached 782–838 KB, roughly 200,000 tokens, and the agent stalled on context overflow. A long-context model did not fail visibly: it read about a third of the evidence, checked 38 claims, and returned output that looked complete. Splitting the check by seat checked 101 claims from every seat (Appendix K.2). We therefore ran the fact checker as seat-wise shards, 66 in all. ExpertTwins never asks one agent to hold the panel’s evidence: seats work from their own packets, quotation checks are string matches, and source review is assigned per citation, so verification cost grows with the number of citations rather than with any single context. This is a design property, not an implementation detail. The re-implementation needed a structural change before it could run at all.

#### The fused answer keeps about two-thirds of what the specialists said, including most of what was flagged

Of 745 fact-checked specialist claims across the three single-shot arms, the final answers retained 47.0%, qualified 17.0%, and dropped 36.0%. Fewer than half of the distinct seat–source citation pairs reached the answer (46.0%, 45.6%, 27.8%), and the fused text was 16–22% as long as the specialist outputs it consolidated. The fact checker’s verdicts did not gate integration: 74 of the 85 claims it marked unsupported or unverifiable (87.1%) still appear in a final answer. Attribution also thinned with the model: surviving claims were tied to their seat in 81.8%, 96.7%, and 58.4% of cases. ExpertTwins fuses nothing. Every claim stays in the per-seat record with its verification outcome, so a failed quotation is a visible property of the claim, not an input to an editor’s discretion.

#### Disagreement between agents was mostly resolved, sometimes erased, and rarely shown

The consistency checker identified 142 contradictions and tensions between agents across the twelve single-shot runs. The integrator presented 19.0% of them to the reader as disagreements, resolved 56.3% in favor of one side or a synthesis with a stated reason, collapsed 22.5% to one side without indicating any disagreement, and left 2.1% unaddressed. Resolution with a stated reason is the integrator’s designed behavior [42] and not a defect, but collapse is a loss. It varied by model (collapsed or omitted in 26.3%, 12.5%, and 33.3% of cases for MM1, MM2, and MM3, respectively), and MM3’s integrator named the seats on each side for only 4 of the 20 disagreements it presented or resolved. ExpertTwins keeps every stance it records, including 127, 16, and 90 explicit disagreements in NBRT1, NBRT2, and NBRT4, attributed to the seats that held them.

#### The fused answer answers what the panel declined to

A second judge compared each final answer with the ExpertTwins record for the same question and model. Of 60 question parts on which ExpertTwins seats had abstained, MirrorMind answered 36, flagged 22 as unresolved, and omitted 2. Of 96 points on which ExpertTwins seats visibly disagreed, the MirrorMind answer showed the disagreement for 24, resolved it for 40, and hid or omitted it for 32. Substantive overlap was nonetheless considerable. Of 180 principal ExpertTwins positions, 119 (66.1%; 68.3%, 50.0%, and 80.0% by model) had a counterpart in the MirrorMind answer. Of 144 MirrorMind positions, 73 were shared, 58 differed in a material detail such as a threshold, ranking, population, or comparator, and 13 had no counterpart. The most consequential differences ran in one direction. At turn four, MM1 rejected the anti-GD2 intensification on which all nine NBRT1 seats had converged, and substituted a cytokine-partner randomization that one seat had mentioned and two would have excluded. At turn two, MM2 credited a seat with a myeloid combination that the seat had explicitly rejected. At turn three, MM3 supplied biopsy thresholds and a trial budget of roughly 3–7 million US dollars, where nearly all NBRT4 seats had withdrawn their numbers or abstained on cost as unestimable from their evidence. In each case, the fused answer turned abstention or dissent into an apparent panel conclusion.

#### Two results from the debate comparison recur

MirrorMind pointed at real records: none of 2,178 specialist citation instances referenced an absent passage, and the only bad pointers were two misspelled seat names in one MM3 final answer. Yet it carried almost no checkable span, because its specialists are not asked to quote: 3 of 674, 8 of 323, and 0 of 272 cited specialist sentences contained a quotation found verbatim in the retrieved text. Its author agents draw only on the author’s own papers, as specified, and 23.6–25.4% of their distinct cited sources were neuroblastoma-specific, against 28.5–40.5% in the matched ExpertTwins runs. The coordinator’s expert-localization step recruited six of nine seats for the opening question in every arm and nearly all seats thereafter. ExpertTwins instead polls every seat on every turn, and a seat with no relevant evidence abstains on the record. Supplying the prior-turn record at turn two did not change the pattern (Appendix K.2).

#### The re-implementation itself is an open challenge

MirrorMind could be run here only after changes that the preprint does not specify or that the run forced (Appendix K): a dense encoder chosen because none is named, a hand-built sparse index, a seat-sharded fact checker, persona and semantic memories distilled once with Opus 5, and our own prompt wording at every stage. Any of these may have changed MirrorMind’s output in either direction. The comparison is therefore between ExpertTwins and our reading of MirrorMind. The least dependent results on those choices are structural ones: a single fused answer carries abstentions and attributed disagreement only if its integrator elects to, and a fact checker that must see every agent’s evidence does not fit in one context. The comparison does not establish whose conclusions are scientifically better. The content judge could not be blinded, because the output formats identify the systems, and neither output was scored against an external standard.

### 3.15 Six computational agents implemented different MYCN prediction pipelines

We now turn from deliberation to execution. Six scientist-associated agents (CS3, CS4, CS2, CS5, CS6, and CS1) were asked to predict a MYCN-related label from neuroblastoma whole-slide images. Opus 5 was the common base model, and each seat was initially allocated two A100-80GB GPUs and 40 pinned CPU cores (a documented allocation, not a measurement of equal realized compute). Seats wrote analysis plans, implemented pipelines, and returned results under shared cohort, split, and metric interfaces. The top three round-one seats read the leading first-round reports, revised, and underwent external evaluation (Section 2.6; Supplementary Methods S8). The internal cohort, from the COMET patient-matched pediatric solid-tumor resource [96], comprised 306 slides from 232 patients (75 positive, 231 negative). The external collection, from the Childhood Cancer Data Initiative Molecular Characterization Initiative [97], comprised 75 slides (23 positive, 52 negative). The exercise is a second case study of the same approach, not a prospectively traced test of a particular roundtable objection.

All agents were given tile features of UNI-v2 [12], GigaPath [13], Virchow2 [115], CTransPath [116], and CONCH [14], provided by the COMET project [96]. The first round completed 235 configuration entries across six seats, and the second added 129 across three seats. These 364 archived entries are not independent designs. For instance, eight of CS3’s second-round entries rematerialized first-round anchors from existing predictions. Search breadth differed across seats, which precludes a budget-matched comparison of problem-solving quality.

**Table 9.**
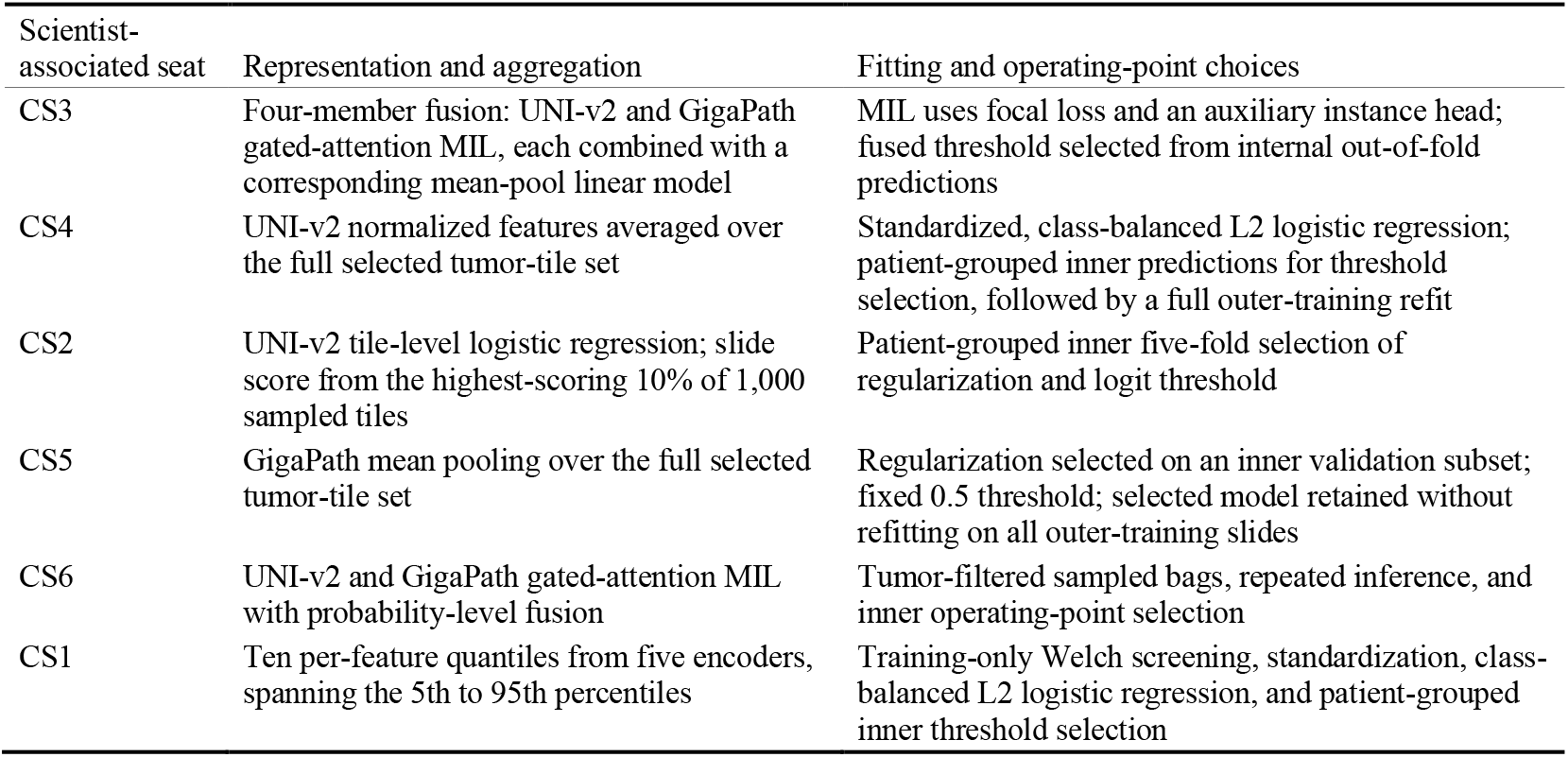
Implemented first-round nominees. All rows describe agents, not methods chosen by the named researchers. Encoder names follow the archived configuration labels (UNI-v2 [12], GigaPath [13]); exact feature-cache and checkpoint provenance remains necessary for full reproduction.

| Scientist-associated seat | Representation and aggregation | Fitting and operating-point choices |
| --- | --- | --- |
| CS3 | Four-member fusion: UNI-v2 and GigaPath gated-attention MIL, each combined with a corresponding mean-pool linear model | MIL uses focal loss and an auxiliary instance head; fused threshold selected from internal out-of-fold predictions |
| CS4 | UNI-v2 normalized features averaged over the full selected tumor-tile set | Standardized, class-balanced L2 logistic regression; patient-grouped inner predictions for threshold selection, followed by a full outer-training refit |
| CS2 | UNI-v2 tile-level logistic regression; slide score from the highest-scoring 10% of 1,000 sampled tiles | Patient-grouped inner five-fold selection of regularization and logit threshold |
| CS5 | GigaPath mean pooling over the full selected tumor-tile set | Regularization selected on an inner validation subset; fixed 0.5 threshold; selected model retained without refitting on all outer-training slides |
| CS6 | UNI-v2 and GigaPath gated-attention MIL with probability-level fusion | Tumor-filtered sampled bags, repeated inference, and inner operating-point selection |
| CS1 | Ten per-feature quantiles from five encoders, spanning the 5th to 95th percentiles | Training-only Welch screening, standardization, class-balanced L2 logistic regression, and patient-grouped inner threshold selection |

The implementations differ in aggregator, representation, effective training-set size, operating-point policy, and model-selection procedure. Some inherited labels did not describe executed behavior. CS4’s mean-pool path averaged all selected tiles rather than a fixed 1,024-tile bag, and refitted a final linear model rather than averaging five inner-model predictions. CS2’s top10 and top20 denote percentages of tiles, not counts, and CS1’s displayed first-round identifier truncates a five-encoder configuration. This breadth is evidence of methodological differentiation under combined corpus-and-rubric conditioning, not an ablation of corpus effects or a validation of the named scientists’ behavior. Attention-based MIL, for example, is an established method [22]; selecting or declining it is not a discovery.

### 3.16 Different methods reached competitive internal performance, and report exchange exposed partition sensitivity

**Table 10.** First-round internal nominations. Balanced accuracy (BA) is the mean over 100 official repeated slide-level holdouts. Percentile ranges describe split-to-split dispersion, not population confidence intervals.

| Seat | Completed configuration entries | Mean balanced accuracy | 2.5th–97.5th split percentiles |
| --- | --- | --- | --- |
| CS3 | 41 | 0.8025 | 0.6929–0.8857 |
| CS2 | 38 | 0.8000 | 0.6948–0.8908 |
| CS1 | 54 | 0.7986 | 0.7107–0.8985 |
| CS4 | 35 | 0.7964 | 0.6806–0.8857 |
| CS5 | 47 | 0.7758 | 0.6397–0.8695 |
| CS6 | 20 | 0.7717 | 0.6524–0.8637 |

The leading four means lie within 0.0061 BA. This is a close descriptive grouping, not a demonstrated equivalence class. Nominations were selected on the same split bank used to report their means, and the comparisons span many configurations with dependent outcomes, so selection bias can be substantial even when training and testing appear well separated [18,23]. The results do not show that pooling universally beats attention. Several linear or pooled models were competitive with several MIL implementations, yet the best model was an ensemble that included attention-based MIL. Furthermore, within some seats the attention variant beat the pooled one. Each score also reflects a combination of architecture, feature encoder, decision threshold, and tuning procedure, so the effect of any single choice cannot be isolated. Second-round work drew deliberately on first-round findings. CS1 moved to three large encoders, L1 regularization, a stability screen, training-score rescaling, and a shrunk operating threshold. CS2 changed its top-tail aggregation, and CS3 kept its first-round fusion as its best nomination.

**Table 11.** Official and alternate-partition second-round summaries. Alternate-partition values are supported by the shared comparison artifact; their primary prediction files were not available within the inspected analysis root. All columns concern the same internal cohort.

| Seat | Round-1 official BA | Round-2 official BA | Round-2 alternate-partition BA |
| --- | --- | --- | --- |
| CS3 | 0.8025 | 0.8025 | 0.8088 |
| CS2 | 0.8000 | 0.8025 | 0.8072 |
| CS1 | 0.7986 | 0.8087 | 0.8038 |

CS1’s second-round internal lead did not persist under an alternate bank of new train/test splits. Small development-score differences were sensitive to the evaluation setting, as expected when a few hundred slides are split repeatedly [52]. The agents also ran label permutations, simpler predictors, patient grouping, tumor filtering, feature perturbations, and operating-point changes. Their permutation procedures differed (whole-cohort label replacement, outer-training permutation with original test labels, and inner-training-only permutation). Near-chance results exclude specific gross failures, but not patient overlap, scanner effects, or adaptive selection. Patient overlap was instead tested directly. Four seats reran their nominated configurations on shared patient-grouped splits at a cost of 0.007–0.015 balanced accuracy (for example, 0.796 slide-level to 0.786 patient-level), so slide-level splitting is mildly optimistic by a margin the panel measured rather than assumed (Supplementary Methods S8). CS3’s internal rule-out analysis reported sensitivity 0.9733 and NPV 0.9791 while assigning 23.7% of slides to the negative group, and CS5 evaluated a sensitivity-targeted variant. These are supplementary internal experiments, not externally established policies for omitting the assay.

### 3.17 External evaluation reordered small internal differences and changed with the threshold

Only three seats were evaluated externally. CS3’s first- and second-round nominations were identical, so the six round-specific rows represent five distinct configurations.

**Table 12.** Primary external point estimates on the complete 75-slide sample. All values use the same full-cohort estimand, recomputed from the canonical saved predictions, with thresholds derived from internal data. No population confidence interval is inferred from the repeated-subset summaries.

| Distinct nominated configuration | AUROC | Balanced accuracy | Sensitivity | Specificity | TP / FP / FN / TN |
| --- | --- | --- | --- | --- | --- |
| CS3 R1 = R2, mixed MIL/linear fusion | 0.7885 | 0.6936 | 0.5217 | 0.8654 | 12 / 7 / 11 / 45 |
| CS2 R1, top 10% | 0.7023 | 0.6066 | 0.3478 | 0.8654 | 8 / 7 / 15 / 45 |
| CS2 R2, top 20% | 0.6973 | 0.6162 | 0.3478 | 0.8846 | 8 / 6 / 15 / 46 |
| CS1 R1, five-encoder proxy procedure | 0.6455 | 0.6162 | 0.3478 | 0.8846 | 8 / 6 / 15 / 46 |
| CS1 R2, three encoders and stability screen | 0.6463 | 0.5849 | 0.3043 | 0.8654 | 7 / 7 / 16 / 45 |

CS3’s fusion had the highest external estimates, yet it missed 11 of 23 positive slides (PPV 0.6316, NPV 0.8036). The predictions carry information, but they do not support replacing or omitting a molecular assay. Rank did not fully invert: CS3 led both the first-round internal and the external tables. What failed to transfer was CS1’s small second-round internal advantage. Round-two effects depended on the metric. CS2’s BA rose by 0.0096 (one fewer false positive) while its AUROC fell by about 0.0050, and CS1’s AUROC changed little relative to the proxy first-round procedure while its BA fell by about 0.0314.

The finite-sample summaries in Table 13 come from 100 unstratified subsets of 67 of the 75 slides (Supplementary Methods S8). CS3’s canonical result averages five seeds and uses a threshold of 0.265598, selected from averaged internal out-of-fold probabilities. A fixed 0.5 threshold instead gives BA 0.5773 (four true positives, one false positive, sensitivity 0.1739), because the fusion’s external probabilities run low (mean 0.221 against a prevalence of 0.307) while still separating the classes. The choice of threshold thus changes the clinical meaning of the same score ordering.

**Table 13.** Secondary external finite-sample stability summaries. The SD is across overlapping subsets.

| Configuration | Mean subset BA | Subset BA SD |
| --- | --- | --- |
| CS3 R1 = R2 | 0.6941 | 0.0192 |
| CS2 R1 | 0.6074 | 0.0178 |
| CS2 R2 | 0.6170 | 0.0176 |
| CS1 R1 | 0.6153 | 0.0190 |
| CS1 R2 | 0.5835 | 0.0182 |

### 3.18 The base model shaped the computational panel more than the persona did

We repeated the Opus 5 exercise five times, with every seat, the moderator, and the operator agent swapped to Sonnet 5, Haiku 4.5, GPT-5.6 Sol, GPT-5 mini, or Gemini 3.8 Flash. The 100 slide-level internal splits, the evaluation harness, and the external cohort were unchanged.

**Table 14.**
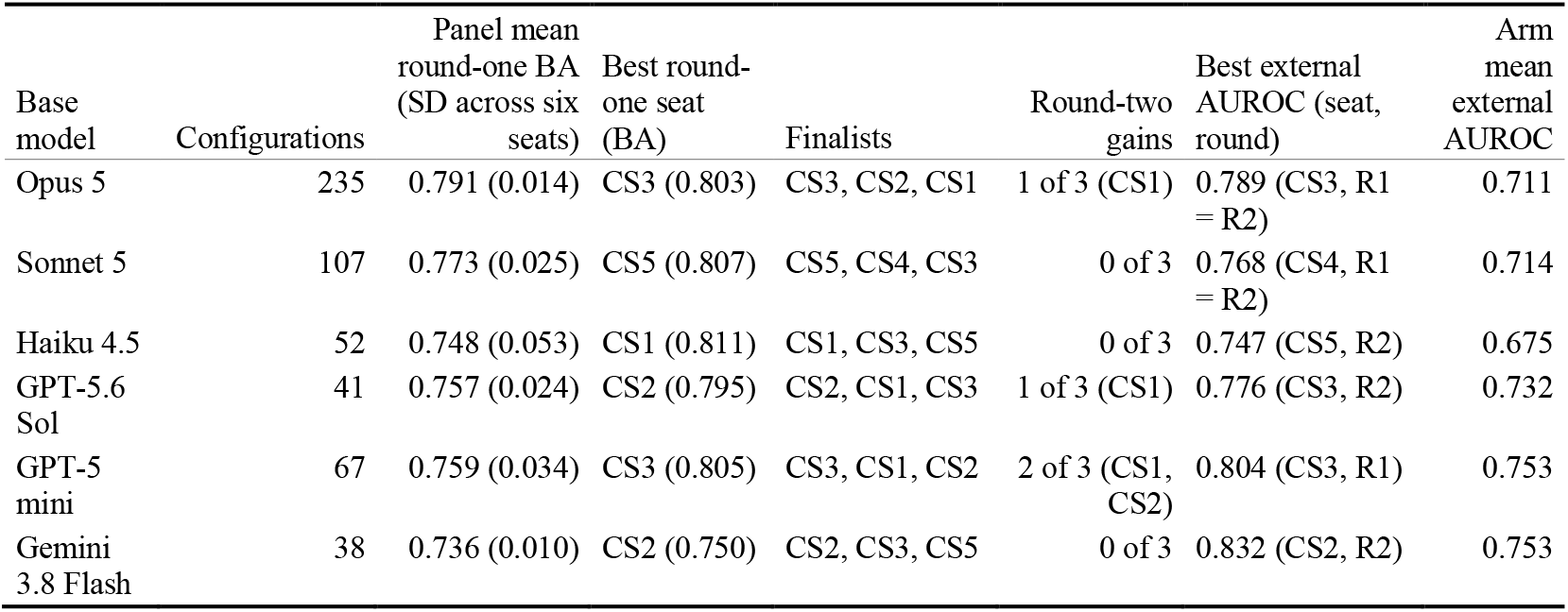
The computational panel under six base models. Internal BA is the mean over the 100 official splits. “Configurations” counts distinct nominable configurations evaluated by the six seats in round one. Round-two gains are paired Wilcoxon tests over the 100 splits at p-value < 0.05. External AUROC is threshold-free; the arm mean is over the arm’s six externally scored nominations (three finalists, two rounds). BA at each seat’s own frozen rule is given in Table 15.

| Base model | Configurations | Panel mean round-one BA (SD across six seats) | Best round-one seat (BA) | Finalists | Round-two gains | Best external AUROC (seat, round) | Arm mean external AUROC |
| --- | --- | --- | --- | --- | --- | --- | --- |
| Opus 5 | 235 | 0.791 (0.014) | CS3 (0.803) | CS3, CS2, CS1 | 1 of 3 (CS1) | 0.789 (CS3, R1 = R2) | 0.711 |
| Sonnet 5 | 107 | 0.773 (0.025) | CS5 (0.807) | CS5, CS4, CS3 | 0 of 3 | 0.768 (CS4, R1 = R2) | 0.714 |
| Haiku 4.5 | 52 | 0.748 (0.053) | CS1 (0.811) | CS1, CS3, CS5 | 0 of 3 | 0.747 (CS5, R2) | 0.675 |
| GPT-5.6 Sol | 41 | 0.757 (0.024) | CS2 (0.795) | CS2, CS1, CS3 | 1 of 3 (CS1) | 0.776 (CS3, R2) | 0.732 |
| GPT-5 mini | 67 | 0.759 (0.034) | CS3 (0.805) | CS3, CS1, CS2 | 2 of 3 (CS1, CS2) | 0.804 (CS3, R1) | 0.753 |
| Gemini 3.8 Flash | 38 | 0.736 (0.010) | CS2 (0.750) | CS2, CS3, CS5 | 0 of 3 | 0.832 (CS2, R2) | 0.753 |

#### Search breadth scaled with the model; internal accuracy barely did

Opus evaluated two to six times as many configurations as any other model and produced the highest and tightest panel, with all six seats within 0.031 BA. Yet its best seat was below the best seat of Sonnet, mini, and Haiku. The models differed more in spread. Haiku’s seats spanned 0.147 BA, from the best single round-one result in the study (CS1, 0.811) to the worst (CS2, 0.664), whereas Gemini’s spanned 0.030 because five of six seats nominated the same single-encoder design. A two-way decomposition of the 36 round-one results attributed 14% of variance to the seat, 29% to the base model, and 57% to their interaction: which persona won depended on which model played it. With one run per seat-by-model cell, the interaction term also absorbs run-to-run variance and should be read as an upper bound. Four different seats won round one across the six arms. CS3 reached the final in every arm and CS6 in none.

#### Weak models compressed the personas toward a house default

Opus, Sonnet, and Sol each produced six distinguishable designs. Four of six Haiku seats nominated UNI-v2 alone and a fifth UNI-v2 with GigaPath, and five of six Gemini seats nominated GigaPath alone. Round two rarely helped internally. Of 18 finalists, four improved significantly on their own round-one nomination (mini’s CS2 and CS1, Sol’s CS1, and Opus’s CS1). Sonnet’s three finalists were all null, and two Haiku finalists resubmitted their round-one configuration unchanged.

**Table 15.** Finalists on the external cohort, all six arms. BA uses each seat’s own internally frozen decision rule and is comparable across arms only in that sense; AUROC is threshold-free. Arrows give round one to round two; “=” marks an identical nomination.

| Arm | Seat | Internal BA | External AUROC | External BA at own rule |
| --- | --- | --- | --- | --- |
| Opus 5 | CS1 | 0.799 → 0.809 | 0.646 → 0.646 | 0.616 → 0.585 |
|  | CS3 | 0.803 = | 0.789 = | 0.694 = |
|  | CS2 | 0.800 → 0.803 | 0.702 → 0.697 | 0.607 → 0.616 |
| Sonnet 5 | CS3 | 0.772 → 0.779 | 0.622 → 0.704 | 0.607 → 0.570 |
|  | CS4 | 0.797 → 0.798 | 0.768 → 0.768 | 0.685 → 0.675 |
|  | CS5 | 0.807 → 0.808 | 0.683 → 0.738 | 0.653 → 0.701 |
| Haiku 4.5 | CS1 | 0.811 = | 0.693 = | 0.558 = |
|  | CS3 | 0.779 = | 0.646 = | 0.544 = |
|  | CS5 | 0.766 → 0.770 | 0.627 → 0.747 | 0.546 → 0.674 |
| GPT-5.6 Sol | CS1 | 0.766 → 0.779 | 0.710 → 0.743 | 0.616 → 0.643 |
|  | CS3 | 0.767 → 0.771 | 0.763 → 0.776 | 0.722 → 0.737 |
|  | CS2 | 0.795 → 0.800 | 0.708 → 0.692 | 0.597 → 0.628 |
| GPT-5 mini | CS1 | 0.799 → 0.817 | 0.683 → 0.747 | 0.621 → 0.665 |
|  | CS3 | 0.805 → 0.801 | 0.804 → 0.794 | 0.711 → 0.720 |
|  | CS2 | 0.743 → 0.787 | 0.741 → 0.752 | 0.577 → 0.595 |
| Gemini 3.8 Flash | CS3 | 0.742 → 0.752 | 0.703 → 0.687 | 0.657 → 0.527 |
|  | CS2 | 0.750 → 0.755 | 0.775 → 0.832 | 0.675 → 0.643 |
|  | CS5 | 0.736 → 0.749 | 0.723 → 0.799 | 0.493 → 0.718 |

#### Internal rank carried little information about external rank

Across the 36 externally scored nominees, internal and external BA correlated at *r* = 0.18 (Spearman 0.13), and mean BA fell from 0.784 to 0.630. Haiku’s CS1, the best internal model in the study, ranked 30th of 36 externally. Opus, the strongest and most consistent panel internally, had the second-lowest arm mean external AUROC (0.711, against 0.753 for mini and Gemini, 0.732 for Sol, 0.714 for Sonnet, and 0.675 for Haiku). The highest single external AUROC (0.832) came from the arm that searched the fewest configurations. Searching 235 configurations rather than 40 bought internal performance, and nothing that transported.

#### The models failed the same way

Thirty-three of 36 models were specificity-heavy externally (mean sensitivity 0.41, specificity 0.85). Eight of the 23 amplified slides were missed by at least 30 of the 36 models, and 13 slides were classified correctly by all 36. Averaging the probabilities of the six arm-best models and calling positives at 0.5 gave BA 0.759 and AUROC 0.815. That BA exceeds every single nominee’s (best 0.737) by roughly one or two slides, and the AUROC is below the best single value (0.832). Base-model diversity produced little error diversity on this cohort.

#### The base model shaped predictions more than the persona did

Spearman correlation between external probability vectors was 0.88 for the same seat’s round-one and round-two models, 0.71 for different seats in the same arm, 0.64 for the same seat in different arms, and 0.63 for different seats in different arms. A persona played by two models was no more similar to itself than to a different persona, whereas two personas played by one model were measurably alike. Within-arm diversity (mean pairwise correlation among round-two finalists; lower is more diverse) ranked Sol 0.61, Opus 0.70, Sonnet 0.73, Haiku 0.74, Gemini 0.76, and mini 0.91; mini’s three finalists were nearly the same model.

#### Round two helped externally where it had not helped internally, but inconsistently

Round-two nominees beat their own round-one nominee on external BA in 10 of 18 cases, lost in 5, and were identical in 3. All three Sol and all three mini finalists improved, while two of three in each of the Opus, Sonnet, and Gemini arms did not. Gemini swung most widely (CS5 BA +0.225, CS3 BA−0.130): the least internally reliable model produced both the best and some of the worst transfers from a single round of peer reading.

Every arm ran the protocol correctly, yet in five of six arms the internal top nominee was not the external top by BA. Sonnet’s CS5 was the exception, and by AUROC only Gemini’s CS2 led both tables. Within-arm rank correlations between internal and external BA ranged from −0.26 to +0.77. The base model determined how widely a seat searched, how faithfully it played its persona, and what its panel’s predictions looked like, but none of these determined which model transported.

## 4. Discussion

ExpertTwins organizes a scientific panel around traceable differences in evidence and method rather than role labels. This is heterogeneity with content: the outputs imply different experiments, not different styles of explanation. The ten-turn study adds that agreement on a decision (a regimen not ready for trial) can coexist with distinct reasons, unresolved mechanisms, and different conditions for changing the decision (Appendix D). Therefore, consensus and collapse should not be equated. Monitoring makes the trajectory inspectable, but scientific interpretation supplies the meaning.

### Within-field plurality is a reason to construct more specific experts

Immunology, like most fields, contains competing mechanisms, methods, and evidential standards, especially where observations do not yet discriminate among hypotheses. An agent told to act as an immunologist may drift among incompatible assumptions or merge them into a unified account that hides the uncertainty. Seats grounded in a scientist’s publications or a narrower research tradition aim at coherence within each seat and real difference between seats, so a researcher can see which uncertainties are shared, which depend on particular assumptions, and which experiments would discriminate. The caveats are that a scientist’s publications need not express one stable position, that publication-grounded agents cannot be assumed to share that scientist’s beliefs, and that rigid conditioning could reinforce selective use of evidence. Whether this construction yields more consistent yet revisable positions, better-identified open questions, and more discriminating experiments than disciplinary prompting is an empirical question.

### Source review turns the limits of a claim into a research output

The record lets a reader ask why a seat encountered a source, whether it was attributed correctly, what was quoted, how the claim changed, and how a computational recommendation fared. The benefit is not correctness at every step but the ability to locate where a chain of reasoning stops being supported. Each layer tests something different: retrieval establishes exposure, quote checks establish provenance, source review challenges inferential scope, and the computational study tests executable recommendations. A drop in external performance is therefore not the same failure as a nonexistent citation, and a source-review rejection does not make an idea worthless. The repair depends on the layer that failed: attribution correction, better retrieval, narrower claims, improved design, or rejection of the method.

### The computational case study makes evaluation discipline part of the output

Six agents took different routes through one task, emphasizing representation, aggregation, thresholds, stress tests, or negative controls. Changing the perspective changed which analyses were built and which weaknesses were probed. Because the agents differed in corpora, rubrics, and research histories, the match between their methods and their associated publications reflects the combined conditioning. A publication-conditioned assistant may thus help a researcher prepare an analysis, test how a result depends on model and threshold choices, or sharpen a question before consulting a colleague, much like the publication-grounded rehearsal of collaborator conversations in interdisciplinary-discovery tools [44]. That benefit must be weighed against acquisition, inference, verification, and review costs (Appendix E).

### The base model realizes a perspective; it does not merely fill a role

Identical packets yielded a physiological confound in one conversation, a more detailed rescue experiment in another, and the same proposition with a different evidential boundary in a third. Such variation helps when it adds a valid control and harms when it adds an unsupported claim. Two kinds of reliability therefore matter: operational reliability, which preserves the citations, and scientific reliability, which keeps claim strength aligned with evidence proposed. The four high-grounding runs met the first and showed useful self-correction. Where ground truth existed, six computational panels each ran correctly and produced confident internal leaderboards, yet internal rank poorly predicted external rank (*r* = 0.18 across 36 nominees). A persona played by two models was no more like itself than like another persona. Base-model variation is known to create heterogeneity among agents [41], and it should be reported like the corpus, roster, and question set.

### Relation to persona-grounded scientific panels

ExpertTwins shares three premises with recent persona-grounded systems: differentiated knowledge improves group reasoning, publication corpora are a defensible way to differentiate agents, and the evidence store should be separable from the agent that uses it. MPDS’s ablation, in which role-specific evidence pools and debate each add to an evidence-only baseline [24]. The lower correlated error of deliberation over disjoint private evidence [89] additionally supports differentiation in retrieval.

The designs part on four points, each visible in our head-to-head runs. The first is the unit of grounding. MPDS pools abstracts [24] and MirrorMind distills memory tiers [42]. Therefore, support cannot be mechanically relocated. In contrast, here, a citation passes only when its quoted span is found in a permitted document, which held at 97.9–99.8% across four base models. On the other hand, no debate sentence of MPDS and only 0.0–2.5% of MirrorMind sentences carried a checkable span (Sections 3.13 and 3.14). The second is what happens to agreement. MPDS scripts convergence [24] and MirrorMind fuses outputs into one answer [42]. Because shared weights, corpora, and visible outputs correlate agents [31–35], we report convergence as a measurement instead. Agreement among strong models fell from 31% to 5% of claims, and 236 abstentions stayed on the record, whereas the MirrorMind integrator collapsed 23% of its agents’ disagreements without trace. The third is fidelity: MirrorMind aims to simulate a scientist’s reasoning [42], whereas we specify a corpus, not a mind. The fourth emerged only when the architecture was run: a fact checker that must see every agent’s evidence does not fit in one context, whereas ExpertTwins partitions verification by seat and citation (Appendix K).

### Long-horizon deliberation needs instrumentation, not only architecture

Published persona panels are short or unmeasured over time. MPDS ends at three rounds [24]. CA-ChemE ran seven knowledge-base-equipped experts for 1,200 dialogue rounds and reported exchanges that closed without reaching the technical question, which a dedicated collaboration agent had to repair [25]. Two language models conversing without external input drift into repetition [95]. Silent degeneration is thus the characteristic failure of long panels, and it can be detected only if the quantities that reveal it are recorded. Our ten-turn panel introduced new documents at every turn, with 24% of turn-ten citations new (Appendix D), and every turn records participation, abstention, stance movement, and position separation. Repetition can therefore be counted: re-filed claims were 14 of Gemini’s 261 and none of Opus’s, Sonnet’s, or Sol’s (Section 3.11). Because the intervention ladder never fired in these runs, the observed behavior belongs to the protocol and the models, not to a controller.

### A guardrail that never fires is not evidence of a guardrail that works

We therefore exercised it directly. When a moderator prompt dictated the panel’s output, position separation fell to zero, the ladder escalated through five interventions in the designed order, and, when none restored separation, it stopped the panel and declared collapse instead of emitting a consensus (Appendix H). Grounding stayed at 100% and participation entropy at its maximum, because each seat filed an equal share of identical claims. Neither quotation fidelity nor equal airtime detects this failure, although the idea-flow map shows it at a glance (Figure 6). Three further runs placed one strong seat among weak ones. The strong seat took roughly half the record, and the others agreed with it about 80% of the time, yet separation rose and no rung fired (Appendix I). The reason is structural: the axes are computed over filed claims, so a seat that agrees more and claims less removes its own rows instead of moving them toward anyone else. Forced consensus and asymmetric deference are orthogonal failure modes, and the present instrument detects only the first. The result points to specific repairs: a banded stance-movement axis, a claim-share floor, and agreement recorded by speaker and target.

### Idea-flow maps change how a panel record is read, not what it contains

A nine-seat, four-turn panel files several hundred claims and several thousand stances, which the monitor reduces to a few numbers per turn. The map (Section 3.7) sits between the two. It locates the events behind a statistic, so a fall in agreement or a rise in withdrawals can be traced to specific ideas, seats, and quotations. It shows collapse as shape, and dissent or asymmetry as structure: a single-seat lineage that survives contradiction, or one seat that wrote most of the ideas others built on. Unlike live argument mapping, it asks nothing of participants during the discussion [99,100], and unlike post-hoc conversation visualization, it does not have to infer reply structure from untyped text [106,107]. Its edges are stances that seats filed against named targets, and every idea opens onto verified quotations, an affordance closer to citation-context indexes [108] than to argument mining from free text [109]. It remains a reading aid, not a measurement: ideas are similarity clusters, stance targets are resolved heuristically, and any relationship it suggests must be confirmed against the record.

### What these systems do that we do not

MPDS evaluates against time-locked literature in a blinded comparison of 30 matched cases, separates persona induction from debate by ablation, and adds a laboratory follow-up [24]. MirrorMind disambiguates authors by persistent identifier, represents collective knowledge as concept graphs, and reports quantitative baselines with human novelty assessment across 15 scholars [42]. Executable systems set a further bar: Robin’s proposal was tested at the bench [87], and independent scientists audited the accuracy of Kosmos’s statements [88]. Our two architecture comparisons fix the base model and the questions, but each rests on one panel question rather than a case series, and only the MPDS adjudication was blinded. We resolve identity by name and affiliation with an attribution audit, and we represent the collective tier as a retrieval policy rather than a graph.

## Limitations

Budget-matched comparisons are needed to establish how much publication-grounded panels improve on strong single-agent, repeated-sampling, and simpler multi-agent workflows. The MPDS comparison fixes the base model, seats, and initial question but matches neither token budget nor question count and rests on one question. Therefore, it shows differences in what each architecture records, not an advantage in claim quality. The MirrorMind comparison is with our re-implementation, not the authors’ system. The changes needed to run it may have altered its outputs in either direction. The content judge could not be blinded, and neither system’s answers were scored against an external standard. The base-model study measures sensitivity within the framework, not the framework’s added value. Roundtables are resource-intensive: a nine-seat, four-turn panel comprises 36 seat-turns before repairs or audits, and the primary closing audit prepared 1,704 referee-citation assignments (2,080 in the anchored run). These are workload counts, not monetary estimates. Appendix E specifies the experiments needed for rigorous architectural, fidelity, and controller conclusions. Independent questions and repeated runs are few. Every cell of the base-model comparison (six models in two roundtable arms and six computational arms) was run once, so between-model differences are not separated from run-to-run variance. The endogenous moderator was always Opus 5, so moderator-model effects are unmeasured. Final-turn audits were performed by seats of the audited model, so adverse rates conflate citation quality with auditor strictness and are exposed to evaluator self-preference [39]. Several Gemini and GPT-5 mini dispatches returned empty and were re-run, a deviation from single-shot execution. Cross-model thematic comparisons describe the inspected conversations, not population-level rankings. Attribution checks rely on imperfect bibliographic records, and a correctly attributed publication does not prove an individual’s contribution or present belief. Corpus, identity, and rubric effects are not yet fully separated. Quotation validity and scientific support remain distinct: source review identified consequential scope problems, but its judgments are model-generated. The ten-turn trajectory is an adaptive developmental run, and lexical novelty is not conceptual novelty. The computational exercise concerns one task, a modest cohort, and one external validation set. The idea-flow maps rest on heuristic clustering, stance-target resolution, and withdrawal detection, and their topic labels are model-written summaries checked only automatically; none of these has been validated against human annotation of the same records (S9).

## Responsible representation

An ExpertTwin is a retrieval- and rubric-conditioned agent associated with a public body of work; it is not the named scientist. Generated statements, methodological choices, reviews, and model scores belong to the agents and this workflow, not to the individuals whose names label the corpora. The development record includes identity corrections, so misidentification is a concrete risk: it can contaminate retrieval and create a misleading association between a person and generated claims.

ExpertTwins makes scientific expertise operational as an explicit relationship between a research remit, a body of literature, and an evidential standard. Its panels contribute different experimental commitments, challenge one another at the source, and keep an inspectable record of how a question changes, which the idea-flow maps make readable. The ten-turn study shows that useful discussion persists beyond the first exchange, and the model comparison shows that one panel design yields largely different but equally grounded conversations under different language models. The contribution is a practical way to build and study scientific collaboration in which disagreement has identifiable content and support can be examined separately from eloquence. The next stage is a cost-aware comparative evaluation of when these capabilities add value over simpler workflows.

## AI assistance

AI systems were used to implement the workflow, retrieve and summarize literature, run roundtables and computational analyses, document results, generate figures, and edit text.

## Data availability

The internal neuroblastoma whole-slide images and matched methylation-inferred MYCN calls are from the COMET resource [96]. COMET whole-slide images can be viewed at pecan.stjude.cloud/histology and will be available for bulk download through the COMET project [96]. The associated DNA-methylation data are available through the Gene Expression Omnibus, with cohort browsing at comet.stjude.org. The external validation slides are from the Childhood Cancer Data Initiative Molecular Characterization Initiative [97] and are available through the CCDI data ecosystem under its access procedures. The literature corpus used by the panels consists of published articles and preprints obtained from publishers and open-access repositories; it is redistributed only as the bibliographic manifest, not as full text. Panel records reported here (run manifests, seat specifications, evidence packets, per-turn ledgers, quotation-check outputs, monitoring trajectories, and audit records) are available from the corresponding author on reasonable request.

## Code availability

The base code for implementing ExpertTwins is available at https://github.com/aforoughipour/ExpertTwins. Additional details, such as the following, are available from the corresponding author on reasonable request: the full ExpertTwins framework and runs; the roundtable transcripts, built corpora, and the analysis scripts for the computational case study.

## Ethics

The internal cohort comprises archival, retrospectively collected, de-identified specimens and associated molecular data assembled under the COMET project [96] with institutional review board approval and a waiver of informed consent. The external cohort was obtained as de-identified data through the Childhood Cancer Data Initiative [97]. The present study involved no prospective human-subjects intervention, no patient contact, and no protected health information. Scientist seats are built only from published bibliographic records. Scientist associations denote publication-grounded evidence policies, not replicas, simulations, or endorsements of the named researchers, and all such seats are reported under coded labels.

## Supplementary Methods

These methods record the procedures, parameters, and definitions behind Section 2: the operational detail needed to reproduce or audit a run. S1–S3 cover the evidence library, seat construction, and packets; S4–S6 the turn protocol, verification, and monitoring; S7 the roundtable configurations; S8 the computational study; and S9 the idea-flow maps. The design rationale is in the main text.

### Seat labels and own-work reporting

Scientist-associated seats appear under coded labels (IMM*n*, CS*n*) here and in the appendices. The platform tags each packet document own when the seat’s source scientist authored it and read when it comes from the scholarly neighborhood, and both tags are recorded for every run. Tag-level results are reported in full for discipline seats. For scientist-associated seats, we withhold statements that pair a coded seat with a specific document and an own tag, because the document’s author list would re-identify the scientist. Aggregate and procedural statements about own-work retrieval, including the own-work floor and the inventory in Appendix F, are retained because they name no document.

### S1 Evidence library and acquisition channels

#### Representation

Each document is stored as a manifest entry, bibliographic metadata, extracted passages, and, where available, a source PDF. A local SQLite FTS5 index over the passages supports lexical (BM25 [49]) retrieval in original document order, so the identifiers and passages an agent receives can be examined independently of its prose. The index can be regenerated from the stored passages and is never treated as authoritative.

#### Acquisition

Four channels with different coverage feed the library. Author and topic queries run once against Europe PMC [47] to download candidate works. A preprint channel acquires from arXiv, bioRxiv, and medRxiv and flags each work as a preprint. A canon channel acquires landmark works by exact title from a curated list, and a completeness check reports which listed landmarks are missing. The scholarly-neighborhood channel (S2) reaches works through OpenAlex [48], including computer-vision and machine-learning venues that Europe PMC does not index.

#### Text quality and integrity

Open-access status and extraction availability also shape what enters a packet. A document counts as full text when it exceeds 6,000 characters and five passages. This is not a completeness certificate: XML extraction omits figures, table wrappers, formulas, and some supplementary sections, and PDF extraction includes no OCR recovery. Byte integrity (has the stored content changed?) and bibliographic integrity (does the content belong to the identified work?) are checked separately; the development record contains a case in which they diverged despite consistent hashes. Library counts are dated, and counts from a later library do not describe the material available to an earlier run.

### S2 Seat specification, attribution, and neighborhood construction

#### Specification

A seat specification is a configuration file. For a discipline seat, it lists retrieval queries, coverage probes, and minimum acquisition targets. For a scientist-associated seat, it records the display name, surname, accepted initials, surname aliases for inverted or concatenated index forms, own-work topic tags, affiliations, and three query sets that look alike but serve different purposes:

- `retrieval_queries` are full-text-search queries applied at question time across the corpus the seat can reach;
- `own_queries` are full-text-search queries applied inside the seat’s own-work set;
- `acquisition` queries are literature-service queries run once to download candidates.

Probes are three to five tokens. The rubric has three fields: `territory` (where the work lives and where it stops), moves (what the scientist does with a problem), and `fatal_flaws` (conditions under which the seat must refuse to agree). Fatal flaws are written as checkable conditions, such as “a delivery claim with no measurement of what reached the cell”. The executed prompts name the scientist.

**Attribution** is the falsification step of seat construction. Every document returned by the author queries is checked against its stored author list, using surname-and-initials matching that tolerates multi-token, inverted, and bare-surname index forms. Accepted works form the own-work set, and rejected works are listed. The rejection list is read, not just counted, and a warning is raised when more than half the candidates are rejected. Affiliations separate two scientists whom neither the name matcher nor co-author corroboration can distinguish. ORCID identifiers are declared in every specification but are not yet used. The development record includes identity corrections in which a seat had been built for the wrong person; in each case, the seat, its corpus, and its attribution file were deleted, and the intended scientist’s seat was acquired and attributed from scratch.

**Neighborhood construction** expands the verified own-work set through the citation graph. A planning step queries OpenAlex and writes a reviewable list of every work it intends to fetch, with a reason for each, under five nodes: (i) the scientist’s works as indexed by OpenAlex; (ii) the works the scientist cites most often across their own papers; (iii) the works of their collaborators; (iv) the most-cited works that cite them; and (v) the most-cited works in their topics, whoever wrote them. Works in the fifth node are tagged as field canon so they can be found and removed. The author identifier for this expansion is resolved from the seat’s already-attributed own papers, probed by DOI. An identifier is accepted only when it anchors a minimum number of those verified papers; identifiers anchored by one paper are dropped, and no identifier is assigned when none clears the threshold. Every node is filtered to scholarly work types (article, review, preprint, conference paper, book chapter, report, data paper) before the referenced, collaborator, and canon nodes are derived; records the index has not typed are kept. Fetched works are merged into the library and pass through attribution again, so nothing enters an own-work set without an author-list check. The harness tags an own-work source own and a neighborhood source read.

**Seat verification** asks three questions before a seat is used. Is the literature present (probes return hits and the document count clears the floor)? Is attributed own work held? Is the seat distinct from those already at the table? Distinctness uses pairwise retrieval overlap with a heuristic threshold of about 35%, above which a seat is treated as an existing seat wearing a second label. Verification also reports own-work documents shared between co-authors, which lower the evidence-diversity statistics between those two seats. Distinctness is not usefulness: a panel can be disjoint, probe-coherent, and covered and still contribute nothing.

#### S3 Packet construction

The packet builder assembles a seat’s evidence in ordered strata under two budgets: a seed budget that bounds ranked hits, and a passage budget that bounds the packet after context is added.

1. **Precision.** A conjunction of question terms that are both subjects the corpus writes papers about (in at least 25 document titles) and rare enough to discriminate (in at most 5% of passages). The stratum is empty when fewer than two terms qualify.
2. **Own work.** Question-conditioned retrieval inside the seat’s own-work set, up to the own-work floor.
3. **General.** Retrieval across the rest of the permitted corpus with the question alone, then with each seat query conjoined to the question, then with each seat query bare as a floor, up to the seed budget less a withheld reserve of one fifth.
4. **Coverage top-up.** Direct probes for question terms that no retrieved passage contains.
5. **Breadth.** One passage from each unrepresented document until a minimum document count is reached.
6. **Relaxation.** Any remaining seed budget, spent above the per-document cap.
7. **Context.** A configured number of neighboring passages on either side of each hit, in source order, filled from the seed budget up to the passage budget.
8. **Disclosure.** Outside the evidence budget, documents that other seats cited in the previous turn, anchored on their quoted passage.

Within a stratum, allocation is round-robin over the configured queries. The tunable parameters are the seed budget, the own-work floor, the per-document cap, the minimum document count, the context radius, and the passage budget. Before any model call, a packet audit reports documents, own-work documents, term coverage, the largest single-document share, and a “thin” verdict with reasons; the audit informs but does not gate. This retrieval design has not been compared under matched budgets with dense retrieval, reranking, or knowledge-graph methods.

### S4 Turn protocol and orchestration

#### Orchestration

The orchestration layer prepares packets, imports structured responses, runs deterministic checks, freezes turn records, and renders reports; language-model calls are dispatched outside it. Each seat returns a structured file of claims, each with a stance label, citations, and quotations, plus any abstention or withdrawal. The harness imports this file rather than parsing prose.

#### Information flow

The first turn is isolated. From the second turn on, a seat receives the moderator’s brief, the other seats’ structured positions, and disclosed source context for documents cited in the previous turn, capped per receiving seat. With cumulative context enabled, a seat’s own earlier positions remain available to it. Private holdings and working drafts are never broadcast, cited evidence crosses seat boundaries only under the disclosure cap, and the run record preserves which fields and passages were exposed at each stage. A frozen turn separates generation from audit, so the audit does not rewrite the artifact it assesses.

#### Moderation regimes

In the endogenous regime, the moderator model receives a packet assembled from the run’s own frozen record and writes the next brief; both are archived under moderator\. In the anchored regime, the brief is a fixed question set composed by the operator outside that loop; no moderator packet or brief exists, and the question text is archived in the manifest. Operator-composed briefs were also drafted with AI assistance. The regimes therefore differ in whether the brief was generated inside the run loop from that run’s record, not in whether a model was involved. The operator selected questions and seat configurations, managed runs, and connected deliberation to computational work.

### S5 Quotation verification and source-review procedure

#### Quotation check

A citation passes when the source satisfies availability and scope checks and the quote is found in a single stored passage after normalization, with at least 40 characters after folding. Folding comprises Unicode normalization, case folding, typographic and whitespace normalization, and handling of line-wrap hyphens. Verification searches all stored passages of a permitted document. Repair after a failed check is limited to evidence already available to the seat, and both the failure and the repair remain identifiable in the record.

#### Source review

After a turn, claim–source cases are assigned to peer seats as referees. Each referee sees the quoted passage with a configured number of context passages on either side and returns a categorical judgment (support, overstatement, irrelevance, misreading, cannot tell) with a rationale. Case eligibility, referee assignment, and context radius for each reported audit are given with the run configurations in S7. Referees are seats of the same run, so adverse rates conflate citation quality with auditor strictness; verdict rates are reported only with their denominators.

### S6 Monitoring statistics and intervention policy

Monitoring distinguishes four quantities: the diversity of supplied evidence, the diversity of cited evidence, the separation of generated positions, and movement in stance labels. All are operational descriptors, not measures of truth.

#### Supplied evidence

For nonempty retrieved-document sets *A* and *B*, the overlap coefficient is

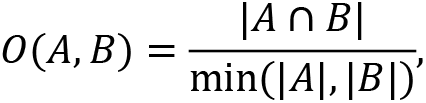

which is not Jaccard similarity. Empty sets are reported as coverage failures.

#### Cited evidence

For cited-document sets *D_i_* and *D_j_*, the evidence-distance statistic is the Jaccard distance

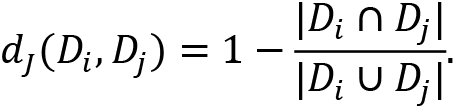

Sources that share an underlying experiment remain different identifiers, so neither statistic measures study-level independence.

**Position separation** is derived from the spectrum of a similarity matrix over seat-level claim text. Writing *λ_r_* for nonnegative eigenvalues normalized to sum to one, the order-2 effective rank and its roster-scaled form are

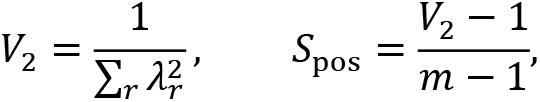

for *m* > 1 participating text groups. *V*_2_ is an effective rank in the sense of Roy and Vetterli [50] and is related to the Vendi family of spectral diversity measures [19]. Its value depends on the text backend, pooling, roster size, and kernel, which must be held fixed across compared runs.

#### Stance movement

Signed transitions into and out of agreement, *n*_→_ and *n*_←_, give the directional herding statistic

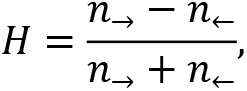

defined when at least one transition is counted. A negative value indicates movement away from agreement, not improved quality.

#### Reference bands and permutation null

The isolated first turn provides a within-run reference. Later statistics are compared with retention bands adjusted for packet separation and a lexical contestedness proxy; these are engineering choices, not learned parameters. A permutation diagnostic compares observed separation with separation after redistributing the same material among seats. For an observed bounded statistic ℎ and null mean ℎ_0_ < 1,

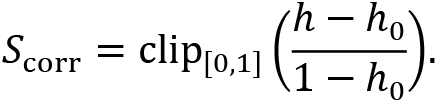

The null is a random partition of the observed material, not a matched single agent. A saturated null leaves the correction uninformative and can produce a spurious `collapsing` verdict on an uninformative axis; it is reported as such, and observed and null values are always reported together.

#### Intervention policy

The intervention ladder and its order are given in Section 2.5. The sequential monitor accumulates likelihood-ratio-shaped evidence toward an action boundary, following the form of a sequential probability ratio test [51], and has not been calibrated to a stated false-positive rate or power. No intervention was applied in any of the twelve base-model runs or in the ten-turn study. The ladder was exercised in the induced-collapse runs of Appendix H, and Appendix I tests a failure mode that it misses.

### S7 Roundtable configurations

#### Language models

Models were called by the following identifiers: `claude-opus-5` (Opus 5), `claude-sonnet-5` (Sonnet 5), and `claude-haiku-4.5` (Haiku 4.5) from Anthropic; gpt-5.6-sol (GPT-5.6 Sol) and `gpt-5-mini` (GPT-5 mini) from OpenAI; and `gemini-3.8-flash` (Gemini 3.8 Flash) and `gemini-3.5-flash` (Gemini 3.5 Flash, used only in Appendix I) from Google. Roundtable seats and moderators ran at medium reasoning effort, except Haiku 4.5, which does not accept the setting.

#### Primary panel and base-model runs

The primary panel and all twelve base-model runs used nine seats, four turns, a 140-passage budget, a 200-seed request, an own-work allocation of 45, at most four passages per document, a minimum of 18 documents, neighbor radius one, retrieval regime 2, cumulative context, and enabled disclosure. Disclosure was capped at 16 peer-cited documents per receiving seat. In the primary arm, the follow-up for turn *t* was written by an Opus 5 moderator from a packet assembled from that run’s turn *t*−1 record. The resulting 1,340–1,700-word briefs were the only text seats received beyond the shared first-turn question. Opus 5.5 was blocked by its content filter on this pediatric-oncology material in three dispatches, and no other moderator was tried. Final-turn audits assigned each eligible verified citation on grounded closing-turn claims to every nonauthor referee, with a context radius of four passages on either side of the quoted passage (one in the anchored run). Cross-run claim matching, within-run repetition, and disclosure-check statistics are defined in Appendix G.3.

#### Ten-turn panel

The ten-turn panel used six seats, retrieval regime 2, a 260-passage limit, a 200-seed limit, neighbor radius one, cumulative context, and an adaptive moderator with four-seat sub-panels at turns 4, 5, 7, and 9 (Appendix D).

### S8 Computational study design

#### Cohort and target

The internal analysis began with 319 eligible slides (81 positive). Excluding 13 slides (six positive) with fewer than 1,000 tiles left 306 slides from 232 patients (75 positive, 231 negative). Fifty-eight patients contributed multiple slides (132 slides), and seven patients had discordant binary labels across slides. The target was a fixed binarization of methylation-inferred copy-number categories. FOCAL_AMP, BROAD_HC_GAIN, BROAD_AMP, FOCAL_HC_GAIN, and HC_GAIN_2P were positive; lower-level gains, neutral calls, deletions, and FAIL were negative. The four FAIL slides in the cohort were therefore mapped to the negative class, a convention we report explicitly. The cohort contained 1,455,705 tiles (median 4,992 per slide, range 1,004–9,003). Tiles are the instances supplied to the image models; they do not increase the number of independent patients.

#### Splits

The official evaluation used 100 repeated stratified 80/20 slide-level holdouts (244 training and 62 test slides, 15 positive test slides). These splits shared patients between training and test: on average, 22.46 of 62 test slides had another slide from the same patient in training, a leakage route that patient-grouped splitting removes [17]. Supplementary patient-grouped splits stratified patients by the maximum binary label among their slides and evaluated the original slide labels after expansion. Their test sets ranged from 53 to 71 slides and 11 to 22 positives, and metrics remained slide-level.

#### External evaluation

The external collection (75 slides, 23 positive, 52 negative) shared no file-names with the internal slides, and its manifest contained no patient or specimen identifiers. The external harness drew 100 unstratified subsets of 67 of 75 slides without replacement, using an advancing random state initialized at 42. Subsets contained 17–23 positives and shared about 59.85 slides (89.3%) on average. Nominations were frozen before external scoring, and thresholds were derived from internal data.

#### Why the official splits are slide-level

Three considerations motivate this choice. First, MYCN status is not constant within a patient in this cohort. Seven patients carried discordant labels across their slides, so collapsing a patient to one label would replace a measured per-slide call with an inferred one. Second, the panel measured patient leakage instead of assuming it away. Four seats reran their nominated configurations on shared patient-grouped splits at a cost of 0.007–0.015 balanced accuracy (for example, 0.796 slide-level to 0.786 patient-level, and 0.809 to 0.799), well inside the 0.051 split-to-split standard deviation. The official numbers are therefore mildly optimistic, not materially inflated, and the direction and size of the effect are documented. Third, and decisive for the comparisons drawn here, the splits, harness, and metric are byte-identical for every seat and every base model, so any leakage is a constant of the benchmark, not a difference between arms. The exercise tests whether the panel protocol yields differentiated, externally evaluable implementations; it does not report a deployable MYCN classifier. The frozen external cohort, not the internal splits, carries the generalization claim. Because patient-level grouping is the general convention in computational pathology [17], the patient-grouped splits above are reported so that readers can see the design under both conventions.

#### Base-model variation

Each arm ran the two-round protocol and the external scoring round, with the shared/ directory linked to the Opus baseline. In round one, each seat worked independently and nominated one configuration by mean internal BA. In round two, the three highest seats read one another’s reports and revised within their own identity. In round three, each finalist’s round-one and round-two nominees were retrained once on the full internal cohort and scored once on the 75 external slides. Round-two gains were assessed by paired Wilcoxon tests over the 100 splits at p-value < 0.05. The variance of the 36 round-one results was decomposed by a two-way seat-by-model analysis. Internal–external agreement used Pearson and Spearman correlation across the 36 externally scored nominees, and prediction similarity used Spearman correlation between external probability vectors. All values were recomputed from saved predictions and result files (`research\mycn_ablation.py`), preferring each seat’s canonical threshold_* artifact where one exists.

### S9 Idea-flow visualization

#### Input

The renderer reads one completed run record. Its inputs are the claim ledger of each turn (seat, claim text, claim type, and citations with their quotations and verification status); the filed stances (responding seat, target seat, the responder’s paraphrase of the target claim, verdict, and reason); declared abstentions; the moderator’s question for each turn; the monitor’s per-turn statistics; and peer source-review verdicts where present. Stance verdicts are agree, disagree, needs other evidence, and out of scope. Building the map calls no language model (the optional topic labels below are written separately), and the output is a single HTML file with the data embedded. Runs recorded by the earlier harness (NBRT18) store no stances and are drawn with continuation and withdrawal edges only.

#### Claim vectors

Each claim is embedded with the all-mpnet-base-v2 Sentence-BERT encoder [110]. A claim usually states its position first and its reasons after, so the vector is the normalized sum of two encodings: the full text, and a headline. The headline is the first sentence, extended by the second when the first is shorter than 120 characters, and truncated at 400 characters. When the encoder is unavailable, a TF-IDF representation over unigrams and bigrams with sublinear term frequency, reduced by truncated SVD to at most 128 dimensions, is used instead. Similarities below are cosines between these vectors.

#### Withdrawals

A claim is flagged as a withdrawal when its first 260 characters contain a first-person withdrawal phrase (I withdraw, retract, abandon, drop, or concede; I no longer hold, claim, endorse, or support; I take back; withdrawn; withdrawing). Its target is the most similar earlier claim by the same seat, with a bonus of 0.03 for claims of the immediately preceding turn.

#### Stance resolution

A stance names a target seat and paraphrases the target claim, but carries no claim identifier. Its target is the target seat’s previous-turn claim most similar to the paraphrase, retained when that similarity is at least 0.35. To draw an edge, the stance is attached to the responder’s current-turn claim most similar to the resolved target. Agree, needs-other-evidence, and out-of-scope stances become edges only when that similarity is at least 0.5, that is, when the responder carried the idea forward. Disagree stances are always kept, because a contradiction need not resemble what it contradicts.

#### Themes and ideas

Claims of all turns are clustered by Ward linkage [111] into round(N/28) themes, clipped to 4–18, where N is the number of claims. Themes with fewer than three claims are folded into the theme with the nearest centroid. Themes are ordered by optimal leaf ordering [112] of an average-linkage tree on the cosine distances between theme centroids, so similar themes are adjacent and receive neighboring hues. Each theme is labeled automatically by its three highest-scoring terms under class-based TF-IDF [113]; these labels can be replaced by topic labels (below) without changing any assignment. Within each turn and theme, claims are clustered again by Ward linkage into ⌈n/6⌉ ideas, where n is the number of claims in the cell. The medoid of an idea is its claim closest to the idea centroid; without topic labels, the idea is titled by the medoid’s first sentence.

#### Topic labels

A medoid is a verbatim claim, and its first sentence often names the deliberation rather than its subject. In NBRT1, for example, an idea about the event-free-survival gain needed to replace the anti-GD2 backbone was titled “The spread of switching thresholds at this table…”. Each idea can therefore carry a topic title of at most 64 characters and a one-sentence gist of at most 220, and each theme a topic label of at most 44. They are written by a language-model agent from a packet that holds, for every idea, only its member claims under seat codes, and each idea label is pinned to the idea’s medoid, so a rebuild that changes the clustering discards it. A deterministic check rejects a label file unless it covers every idea and theme and respects the length limits; titles and theme labels must avoid deliberation words (seat, panel, table, turn, claim, withdraw, propose, sign), gists must avoid first person and seat names, and no label may state a number absent from the idea’s claims. The check also flags titles whose words mostly do not occur in the claims. Labels change no assignment, edge, or position. The box shows the topic title; the tooltip and evidence panel add the gist and the verbatim central claim, and a header switch restores verbatim titles. All seven maps in this paper carry labels that passed the check.

#### Edges

Between consecutive turns, each idea keeps its most similar parent idea when the cosine is at least 0.55, plus a second parent within 0.04 of the first. Each parent keeps its most similar child when the cosine is at least 0.6. These are continuation edges. Resolved stances and withdrawals add counts to the edge between the ideas they connect, creating it if necessary. Each edge is drawn as one kind, by precedence: withdrawal if it carries any withdrawal; contradiction if disagree stances are at least as many as agree stances; mixed if it carries both; then support; needs other evidence; and continuation if it carries no stance. Edges carrying only out-of-scope stances are dropped. Edge width grows with the number of stances carried.

#### Reception and strength

For each idea, the resolved stances that seats filed on its claims in the next turn give one verdict per seat, by the same rules: contradiction, mixed, support, needs other evidence, or out of scope. These verdicts color the bar borders. Strength is a display weight, not an outcome. The raw score is the number of seats that voiced the idea, plus 0.5 per seat whose verdict is support, minus 1 per seat whose verdict is contradiction or mixed, minus twice the fraction of its claims later withdrawn times the number of authoring seats. Ideas of the closing turn, which have no reception, gain 0.25 per authoring seat. Scores are scaled linearly between the 5th and 95th percentiles of the run and clipped to [0, 1]. Strength sets font weight, border width, and opacity, and the encoding can be switched off. An idea is a merge when its parents span at least two themes and a split when its children do, excluding withdrawal edges.

#### Layout

Turns are columns (or rows), and ideas within a turn are ordered first by theme. Their positions along the turn axis are refined by six alternating downward and upward barycenter sweeps, as in layered-graph drawing [114]. Each idea moves toward the weighted mean position of its neighbors in the adjacent turn, with weight 1 + √w for an edge carrying w stances. After each sweep, positions are fitted to the barycenters by isotonic regression under a minimum separation, with an extra gap between themes. Seat roots sit at the mean position of the seat’s turn-one ideas.

#### Viewer

Hovering over an idea highlights its transitive upstream and downstream lineage; clicking pins it and opens the evidence panel. Hovering over a seat highlights the ideas it voiced and their links, and selecting a turn header shows that turn’s moderator question and abstentions. A filter hides support and needs-evidence edges carried by fewer stances than a threshold (default two); contradiction, mixed, and withdrawal edges are always shown. Spotlight presets dim all ideas except those that are contested (a contradiction or mixed verdict or edge), withdrawn, self-revisions (at least half of their claims are withdrawals), merges or splits, broad consensus (voiced by at least two-thirds of the seats with strength at least 0.7), or minority voices (one seat, at least two cited documents). Full-text search covers claims, topic titles and gists, quotations, and document identifiers and titles. Seat names are replaced by the manuscript’s codes by default, and in published figures the tags that mark a cited document as part of a seat’s own work are hidden.

#### Limitations of the procedure

Ideas are similarity clusters, not argument units, and a cluster can join claims that a reader would separate. Stance targets are resolved from paraphrases and can attach a stance to the wrong claim when a seat filed several similar claims. Withdrawal detection is lexical and misses withdrawals phrased in other ways. Strength penalizes contradiction, so a well-founded dissent can look pale; switching the encoding off removes this effect. Continuation edges record similarity, not influence. Topic labels are model-written summaries, checked automatically but not by a domain expert, and can flatten a claim’s qualifications; the verbatim central claim is therefore shown with every label. None of these steps has been validated against human annotation.

## Appendix A. Interpreting the reported units

| Unit | Meaning in this paper | What it must not be counted as |
| --- | --- | --- |
| Document | A source identity represented in the literature library | An independent experiment or a fully extracted paper |
| Passage | An indexed segment of extracted source text | A guarantee of complete methods or contextual coverage |
| Packet | Evidence and instructions supplied to one seat at one stage | All information available in model weights or previous model context |
| Claim | A generated scientific assertion with associated evidence fields | An independently replicated scientific result |
| Citation case | A particular claim-source-quote relationship | Necessarily a distinct claim or source |
| Audit judgment | A referee’s categorical reading of an assigned citation case | Human-adjudicated truth or an independent sample if cases recur |
| Seat-turn | One seat’s participation at a specified turn | A fresh scientific question |
| Run | A configured sequence of panel turns | Necessarily an independent replicate if context or artifacts are reused |
| Internal split | One saved partition of the same development cohort | A new cohort or an independent patient sample |
| External fitted-model score | Performance of one fitted predictor on the external sample | Independent replication of the external population |

Three denominators must be kept apart. Claim-level quote grounding counts claims that meet the harness criterion; citation verification counts individual source-quote checks; and audit proportions count returned review cases. Numerator and denominator must come from the same layer, so an audit overstatement percentage cannot be subtracted from a claim-grounding percentage to obtain an overall reliability rate. An abstention is a recorded response, not a finding that no evidence exists. A model can abstain because evidence is absent, retrieval missed it, the source lay outside its permitted set, or it found no defensible inference. An abstention count therefore measures behavior under an information policy.

## Appendix B. Minimum record for a reproducible panel

A reproducible panel record identifies the question; the corpus and index snapshot; seat specifications and attribution state; retrieval queries; seed, passage, and document budgets; the context-expansion policy; exact packet text; the external agent’s model and version; generation settings; tools and accessible files; prior-session context; and all returned structured outputs. Each record type has further requirements.

**Verification records** give, for each citation, the quoted span, source identifier, normalization and minimum-length rules, the availability and scope decision, and the result. Claim-level aggregation rules must be explicit, and repair records must identify the failed item, the permitted repair evidence, and the revised or withdrawn output.

**Deliberation records** specify which peer fields crossed the communication boundary, which source contexts were disclosed, and when. A statement that agents were “isolated” is insufficient without both the initial packets and the later communication policy.

**Monitoring records** identify the text backend, roster, missing observations, raw and corrected statistics, null construction, reference bands, state updates, and applied actions. Null saturation and missing data must be visible, and any simulator flag recorded as run metadata.

**Audit records** include case-selection rules, referee assignment, the context actually shown, duplicate-case identifiers, verdicts and rationales, and excluded or malformed responses. Agreement analysis requires several referees reviewing shared cases; review of different cases does not support it.

These requirements describe the information needed to reproduce and interpret the protocol. They do not claim that every historical run holds a complete record of every item.

## Appendix C. Minimum record for a reproducible computational comparison

### Target and cohort

Document the prediction target before the evaluation procedure: the assay or inference source, matching rule, unresolved and discordant labels, and the relation between internal and external definitions. Keep a flow from candidate cases to the patients and slides entering each analysis, including quality exclusions and their class distribution.

### Splits

Save splits as explicit indices and check patient grouping before training. Document any tissue, patient, institution, or preprocessing source shared across partitions [17]. A patient check on one model is insufficient if another model in the comparison uses a different split table.

### Training

Specify the encoder and checkpoint, feature extraction and tumor selection, pooling or multiple-instance aggregation, bag sampling, fitted transformations, feature selection, regularization, optimization, stopping rule, and threshold selection. For every learned transformation, state which observations were available when it was fitted.

### Predictions and nominations

Prediction files should identify the configuration and training split behind each probability. Nomination files should be frozen before hidden or external results are opened, and any proxy substituted for an inexecutable implementation should be stated with its scientific consequences.

### Statistics and clinical claims

Statistical comparisons should keep paired predictions for the same units and respect patient clustering, repeated development splits, and selection among candidates. Paired AUROC methods apply when their assumptions hold [21], but slide-level independence should not be assumed when patients contribute several slides. Clinical claims additionally require calibration, a suitable threshold, representative sampling, and a justified use case, none of which follows from a favorable BA or AUROC; their reporting should follow the TRIPOD+AI standard [16].

## Appendix D. Ten-round study: sustained inquiry without conversational collapse

### D.1 Question, participants, and adaptive protocol

NBRT18, archived separately from the base-model runs (D.5), asked whether a literature-grounded panel can sustain substantive discussion beyond a few exchanges. It evaluated a proposed three-component regimen for pediatric rhabdomyosarcoma and neuroblastoma: intratumoral chemotherapy, a local immune-stimulating agent, and pharmacological or physical vascular normalization [53]. The moderator asked whether the combination was ready for a clinical trial and, if not, what experiment should precede pediatric enrollment.

The run predates the endogenous moderator loop and has no moderator packet or brief: every turn was opened by an operator-composed question written from the preceding record and drafted with AI assistance. The panel comprised immunology, pathology, genetics, a vascular-normalization seat (IMM8), rhabdomyosarcoma, and neuroblastoma. Full-panel turns were 1, 2, 3, 6, 8, and 10. Turns 4, 5, 7, and 9 used four-seat subpanels chosen to examine unresolved questions, giving ten turns and 52 seat-turns rather than ten repeats of one prompt. The configuration used retrieval regime 2, a 260-passage limit, a 200-seed limit, neighbor radius one, and cumulative context.

The moderator shaped the discussion actively, asking for a single primary endpoint, an examination of physical normalization, a specification of the immune agonist, and an adversarial reconsideration of the panel’s emerging position. Because each question responded to what the panel had just said, NBRT18 tests sustained moderated inquiry, not autonomous agenda formation or a fixed-protocol effect of corpus allocation.

### D.2 Evidence intake and grounding over ten turns

The ledgers contain 436 claims, of which 426 passed claim-level quotation checks, and 97 abstentions. Of the attached citations, 1,027 verified and ten were quote_not_found; no invented or out-of-scope citation was recorded. The discussion cited 213 distinct documents and added new ones on every turn.

**Table D1.** Ten-round trajectory from saved ledgers and trajectory record. Grounding counts are exact; novelty is the archived lexical score, not an independent scientific-novelty judgment.

| Turn | Participating seats | Claims | Quote-grounded | Cited documents | New documents | New-document fraction | Mean lexical novelty | Abstentions |
| --- | --- | --- | --- | --- | --- | --- | --- | --- |
| 1 | 6 | 63 | 63 | 57 | 57 | 1.000 | 1.000 | 22 |
| 2 | 6 | 6 | 6 | 18 | 9 | 0.500 | 0.765 | 0 |
| 3 | 6 | 36 | 34 | 39 | 19 | 0.487 | 0.710 | 18 |
| 4 | 4 | 27 | 27 | 35 | 16 | 0.457 | 0.738 | 4 |
| 5 | 4 | 28 | 28 | 15 | 7 | 0.467 | 0.810 | 10 |
| 6 | 6 | 86 | 84 | 54 | 17 | 0.315 | 0.676 | 13 |
| 7 | 4 | 63 | 58 | 50 | 33 | 0.660 | 0.798 | 12 |
| 8 | 6 | 30 | 30 | 39 | 15 | 0.385 | 0.732 | 0 |
| 9 | 4 | 52 | 51 | 55 | 25 | 0.455 | 0.766 | 14 |
| 10 | 6 | 45 | 45 | 63 | 15 | 0.238 | 0.723 | 4 |

Turn ten still introduced 15 documents and 45 quote-grounded claims. The largest turn was turn six (86 claims), and turn seven introduced 33 previously uncited documents. Claim volume must be read against the prompts. Turn two asked each seat for the *single* claim it most disagreed with, and each seat filed one; turn three reopened the substance and produced 36 claims; and turn eight posed five numbered questions, to which each of the six seats filed five claims. A sharp fall in claim count would therefore have been a misleading signal of collapse. Lexical novelty is one minus the largest fraction of a claim’s content words found in an earlier claim by the same seat. It excludes within-turn comparisons and does not show that new wording carries a new idea. The historical contested_docs field counts documents cited by more than one seat, not disagreements.

### D.3 How the discussion changed the proposed experiment

The panel broke an attractive combination down into the conditions under which a systemic benefit would be plausible. Early disagreements concerned the gate for advancing the program: delivery during a normalization window, immune effects beyond the injected lesion, or disease-specific local control that preserves standard treatment. Subpanels then asked whether physical normalization and an unspecified immune agonist could carry the roles assigned to them. The adversarial eighth turn exposed a risk: a two-component chemotherapy-plus-normalization fallback may have become a safe-looking answer without evidence of its own. Later discussion moved upstream of scheduling. Before optimizing delivery of an immune agonist, the program needed evidence of a disease-relevant immune-recognition substrate and of a mechanism the intervention could engage.

**Table D2.** Distinct scientific requirements retained in the closing discussion. These are generated experimental recommendations, not completed biological results or clinical advice.

| Perspective | Requirement brought to the shared next-step proposal | Archived locator |
| --- | --- | --- |
| Immunology | Relate tumor antigen or presented-antigen evidence to antigen-presentation machinery, APC maturation, and effector access; distinguish local activation from control outside the injected lesion. | Turn 10, immunology, study proposal and decision rules |
| Pathology | Establish spatial relationships among tumor, antigen-presentation, and immune cells; use banked tissue to test baseline architecture rather than infer function from marker presence alone. | Turns 9–10, pathology |
| Genetics | Separate disease-genomic strata and account for tumor-normal interpretation and germline handling; require evidence that a candidate antigen is recognized, not merely genetically present. | Turn 10, genetics |
| Vascular normalization | Measure payload coverage, the normalization window, and tumor-versus-draining-node exposure; delivery success alone does not demonstrate a systemic immune mechanism. | Turns 9–10, IMM8 |
| Rhabdomyosarcoma | Retain fusion-status and primary-site stratification, standard-therapy constraints, and disease-relevant antigen-presentation requirements. | Turn 10, rhabdomyosarcoma |
| Neuroblastoma | Tie the proposed mechanism to risk group, treatment backbone, and uninjected metastatic compartments; local control of an accessible lesion does not establish whole-disease benefit. | Turn 10, neuroblastoma |

All six seats opposed immediate trial initiation in the original form and recommended a nontherapeutic tissue-based next step, using banked or clinically available material and viable preparations where required. The agreement did not erase differences in primary readouts, mechanistic conditions, or objections to a delivery-only fallback; in this sense the panel did not collapse. The proposed tissue work also had a boundary. Fixed archival sections can show expression and spatial organization, but not induced antigen presentation, live immune recognition, or treatment-dependent killing, and seats that separated archival screening from viable-sample functional experiments made the recommendation more actionable. Proposed sample counts, timelines, and decision thresholds remained design suggestions, not justified power calculations or validated gates.

### D.4 Source audit and developmental context

The turn-nine audit covered 107 eligible citation instances from 51 grounded claims across 54 documents. Four referees returned 364 overlapping judgments: 258 support, 103 overstatement, and three out-of-context. Overstatement was 28.3% of returned judgments; adverse judgments, including out-of-context, were 29.1%. Neither percentage is a fraction of unique citations or a calibrated error rate.

The developmental report records retrieval-query and cumulative-memory repairs, shell-input and notebook-regeneration fixes, and retained superseded turn-three packets. Because moderation was adaptive, subpanel membership varied, and the implementation changed during the run, the trajectory cannot show that any particular component prevented collapse; its positive result is feasibility. Earlier four-question multiround and paired initial-turn studies produced 579 and 427 claims, with 578 and 426 quote-grounded. They are developmental studies, not arms of the model comparison. Their paired runs selected different citations or changed a readiness judgment despite identical packets, which motivates the repeated-run evaluation in Appendix E.

### D.5 Reproducibility

Paths are relative to the NBRT18 study archive, separate from the platform archive holding NBRT1–NBRT17. Primary records are `manifest.json, turns\1` through `turns\10` with `ledger.json` and `report.json`, `roundtable.md`, and `trajectory.json`. Source reviews are in `turns\9\audit\responses`, superseded artifacts in `_superseded`, and metric definitions in `ops\trajectory.py`. Ledger counts were checked against the saved trajectory. These locators identify the inspected archive, not a public release.

## Appendix E. Cost-aware comparative studies for the next stage

### E.1 Purpose and experimental cost

The present studies establish a workflow, scientific outputs, and model-dependent behavior. They do not estimate, under matched budgets, the benefit of publication-conditioned panels over simpler alternatives. Roundtables require corpus acquisition and maintenance, packet processing, repeated generation, repair, disclosure, support audits, and human assessment. Costs grow with questions, model configurations, seats, turns, repetitions, and review assignments. The primary nine-seat panel entails 36 seat-turns before audit. Its final review prepared 1,704 referee-citation assignments; the anchored run prepared 2,080. These are workload indicators, not measured tokens, money, or researcher time.

A staged evaluation is preferable to factorial expansion: freeze questions, input artifacts, endpoints, and cost-accounting rules; compare the complete workflow with strong, budget-matched simpler baselines; and add arms that isolate the cause only after a useful difference is established.

### E.2 Separate the ingredients

A decisive comparison would hold base model, tools, scientific targets, total inference budget, and accessible source universe fixed while varying evidence allocation and communication. Essential arms are a single generalist with broad retrieval and verification; repeated independent samples from that generalist; multiple agents sharing evidence without specialized rubrics; agents with specialized rubrics but shared evidence; and agents with specialized evidence under a common neutral rubric. The complete ExpertTwins configuration is compared with each.

Scientist-associated agents need identity controls. Masking the name tests name conditioning. Exchanging rubrics while retaining corpora tests rubric dependence. Replacing the neighborhood with a size- and topic-matched alternative tests the neighborhood policy. A mismatched-discipline control must be matched for relevance and quality, so that it is not merely a worse-evidence arm.

Communication needs its own ablation, comparing private first-pass generation and a single aggregation step against structured multi-turn exchange, unrestricted draft exchange, and independent continuation, with source review tested as a separate factor. Repeated runs under each fixed configuration are needed to separate between-construction from within-construction variation.

### E.3 Measure useful objections and appropriate revision

For open-ended questions, a useful endpoint is the number of distinct substantive objections that a matched comparator does not raise, deduplicated by scientific issue and graded blind to architecture. A novel but incorrect objection counts against the workflow. The count should come with precision, severity, and actionability: whether the objection is valid, whether it changes an analysis, and whether it leads to a feasible control. Analysis should be paired at the question level rather than assume that claims are independent. Mean task quality should still be reported; predeclared tail end-points supplement it rather than replace it.

A disagreement map should record the disputed proposition, each position, its evidence, the relevant population and context, and the observation that would discriminate between positions, so that genuine disagreement is separated from agents answering different questions. Introducing decisive evidence tests appropriate revision, and introducing irrelevant paraphrases tests consistency. Support review should distinguish empirical assertions, mechanistic interpretations, and proposed experiments; assess the relevant support set rather than demand one quotation per component; and include a shared, blinded, human-reviewed sample with stable case IDs.

### E.4 Test person fidelity without leaking the answer

A temporal evaluation can withhold later publications and ask whether an agent recovers later methodological choices or positions. A retrieval cutoff alone does not eliminate leakage, because the model may have seen the later work during pretraining. The neighborhood, summaries, and rubric may encode it. A defensible study would document this limitation, exclude post-cutoff neighborhood material, build rubrics from eligible evidence only, grade against a held-out target set, and compare the attributed corpus with a topic-matched corpus from other researchers. Direct named-researcher assessment needs participation and consent. The benchmark should test decisions and evidential standards, not literary imitation.

### E.5 Validate the monitor against scientific outcomes

Controller evaluation needs tasks where useful and harmful convergence can be judged independently of monitored features. Examples include recovery of a withheld decisive fact, persistence of a known false consensus, and retention of a valid minority objection. Select thresholds on development runs and test held-out runs, with intervention order and stopping rules fixed in advance. A meaningful recovery result would improve external outcomes relative to no intervention and simpler alternatives, while accounting for extra tokens and evidence exposure. Increasing a separation score is not enough.

### E.6 Prospective reporting and replication of the base-model studies

Questions and repetitions should follow from a declared minimum useful effect and a design that handles within-question dependence and grading error. Results should include frozen assignments, packet hashes, model identifiers, repair histories, token and monetary costs where available, and all failed or incomplete runs. The computational base-model variation (Section 3.18) is a single-run-per-arm study. Replication would need repeated runs per arm under fixed seeds, a patient-grouped external evaluation with a common pre-registered decision rule so that BA is comparable across arms, and a frozen per-arm brief that cannot carry results between arms. Its question was whether the same perspectives lead to different implemented methods and validity checks when the base model changes. For the strong models the answer was yes; the two weakest compressed six personas toward a single design.

## Appendix F. Attributed own-work inventory

The inspected CCT attribution records contain the following counts. Accepted works are in the assembled main library. A count is an attribution-procedure outcome, not a measure of research quality, career output, agent fidelity, or bibliographic coverage.

| Scientist-associated corpus | Accepted own-work memberships | Rejected candidates |
| --- | --- | --- |
| IMM8 | 212 | 1 |
| IMM2 | 183 | 1 |
| CS3 | 179 | 113 |
| IMM3 | 171 | 1 |
| IMM4 | 136 | 1 |
| IMM5 | 131 | 4 |
| IMM6 | 130 | 3 |
| IMM7 | 123 | 2 |
| IMM1 | 122 | 4 |
| CS6 | 40 | 1 |
| CS4 | 31 | 0 |
| CS5 | 21 | 0 |
| CS1 | 20 | 0 |
| CS2 | 17 | 0 |

The total is 1,516 accepted seat-paper memberships, not necessarily 1,516 distinct papers, because coauthored work can enter several own-work sets. The rejection count for an ambiguous name shows that a screen acted, not its sensitivity or specificity. The known-contaminant regression examines selected title fragments for CS4 and CS3 only, so zero hits there do not prove zero contamination. Neighborhood defaults are 400 own-work candidates, 120 referenced works, eight collaborators with twelve works each, eighty citing works, three topic-canon groups with thirty works each, and at least two verified works to anchor an author identity. They are acquisition budgets, not achieved counts. The separately curated landmark checklist had 26 of 29 targets represented and does not define comprehensive field coverage.

## Appendix G. Base-model comparison: provenance, denominators, themes, and examples

### G.1 Runs, arms, and matched inputs

Twelve roundtable runs formed two arms. NBRT1–NBRT6 used run-specific follow-ups; NBRT7–NBRT12 gave all six panels the same four follow-ups. The operator composed those questions from the NBRT7 record outside the loop, making NBRT7 the anchoring source rather than a recipient. No moderator\ directory exists for NBRT7 or any anchored run. Follow-up text appears only as each manifest turn’s question field; the six endogenous runs archive both packet and brief. Because AI assistance helped draft the operator-composed briefs, the NBRT7–NBRT1 contrast is not one of human versus model authorship. They differ by location and effort: NBRT7 follow-ups were composed outside the loop at high reasoning effort, while NBRT1 follow-ups were generated inside it by an Opus 5 moderator limited to that frozen record at medium effort (Section 3.8). Both describe the same conversation. Section 3.2 reports NBRT1; NBRT7 is only a contrast. Model labels follow the record (Opus 5, Sonnet 5, Haiku 4.5, GPT-5.6 Sol, GPT-5 mini, Gemini 3.8 Flash). These are model substitutions, not random-seed repetitions.

The six anchored-arm manifests match on seat names, index and specification references, 140-passage budget, 200-seed request, own-work allocation of 45, maximum four passages per document, minimum 18 documents, neighbor radius one, retrieval regime 2, cumulative context, and enabled disclosure. All nine initial packet hashes are identical. The shared first-turn preflight records packet separation 0.875666 and rubric separation 0.876708. Identical delivered packets are established, although no frozen full-corpus digest was recovered. Each primary-arm run reuses its model’s anchored-arm turn-one responses, so the arms share turn one and diverge only in turns two through four. Later packets differ because cumulative history, verified contributions, and selected disclosure depend on model behavior.

In the primary arm, an Opus 5 moderator wrote turn *t* from `moderator\packet-t{t}.md`, which summarized turn *t*−1: verified claims, abstentions, disclosed sources, citation-check verdicts, and monitor values. The briefs, `moderator\q-0{t}.md`, are 1,340–1,700 words and were the only added seat text beyond the shared first-turn question. Opus 5.5 was content-filter-blocked across three dispatches. No other moderator was tried; moderator-model effects are unmeasured.

Reused moderator text qualifies only the anchored arm. Its turn-two question describes prior agreement. Turn three gives the original Opus panel’s position scores and adverse-review count; turn four gives another baseline adverse-review count. The third question also supplies the Schwannianstroma and developmental-regression alternatives. These numbers and alternatives are prompt content, so shared responses are not independent discovery. The record states medium reasoning effort, but manifests and outputs omit complete provider versions, dispatch IDs, temperatures, token limits, usage receipts, and attempt histories. Sonnet archived response-building and split-audit artifacts in the anchored arm, where wrapper and retry differences are described. In both arms, several Gemini and GPT-5 mini dispatches returned empty and were re-dispatched. The archived repair prompts do not prove that repairs were executed. We compare archived outputs and record-attributed model identities.

### G.2 Completion, quotation checks, and source review

**Table G1.** Response and citation diagnostics, anchored arm. OK is the archived response status. A no-citation claim lacks a recognized citation list but may still include source text.

| Run | OK<br>outputs /<br>36 | Claim-producing<br>seat-turns | Abstention<br>entries | Claims without<br>recognized citations | Grounded claims by turn |
| --- | --- | --- | --- | --- | --- |
| Opus | 36 | 36 | 163 | 0 | 112/113; 109/110; 97/99; 110/112 |
| Sonnet | 36 | 36 | 199 | 2 | 79/81; 68/73; 55/57; 95/96 |
| Haiku | 35 | 27 | 161 | 116 | 21/35; 11/89; 4/61; 7/22 |
| Sol | 36 | 34 | 185 | 0 | 45/45; 59/59; 64/64; 102/103 |
| mini | 32 | 14 | 50 | 6 | 11/12; 0/6; 15/25; 7/7 |
| Gemini | 36 | 36 | 132 | 0 | 88/88; 80/80; 78/78; 84/84 |

Across parsed anchored-arm claims, 2,203 of 2,285 citation instances verified. Opus, Sonnet, and Sol had six, eight, and one `quote_not_found` instances. Haiku had 53 quote mismatches, two non-existent permitted-corpus identifiers, and one quote below minimum length. mini had ten quote mismatches and one nonexistent identifier; Gemini had none. A nonexistent identifier may be a misspelling or alias, not invention. Source-review records contain 1,856 turn-level check entries and 1,511 final-audit entries, separate from 2,285 claim citations. Because review volume, validation status, and verdict distribution depend on schema and assignment, adverse counts require denominators.

**Table G2.** Final-audit workload and recorded outputs, anchored arm. Returns are ingested judgment entries, not proven unique coverage. The last column is output-validation status, not human validation.

| Run | Prepared referee-citation assignments | Returned judgment entries | Entries marked recorded |
| --- | --- | --- | --- |
| Opus | 2,080 | 385 | 385 |
| Sonnet | 984 | 360 | 332 |
| Haiku | 56 | 11 | 11 |
| Sol | 1,416 | 360 | 356 |
| mini | 56 | 35 | 35 |
| Gemini | 784 | 360 | 350 |

Some Haiku and Sol judgments used unrecognized verdict labels. Several Haiku audit responses used unsupported top-level structures, three mini audit files were unparseable, and stable assignment IDs were inconsistent. These records cannot rank models for common-case, blinded biological accuracy. The anchored disclosure record shows what grounding failure costs. The four high-grounding runs supplied the full capped disclosure of 16 peer-cited documents per receiving seat on every follow-up turn. By turn four, Haiku disclosed zero to four documents per seat; mini had none at turn three. Failed grounding removes evidence from later discussion, not only from a reporting score.

**Table G3.**
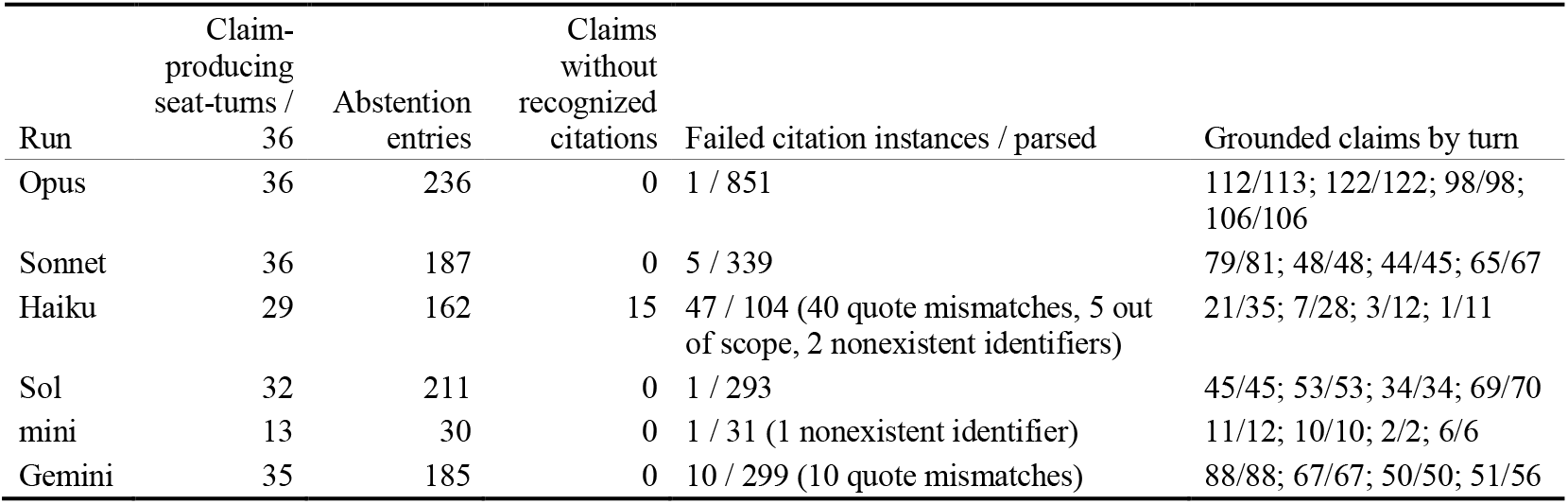
Response and citation diagnostics, primary arm. Failed citation instances have a status other than verified in the run’s ledgers; the breakdown names dominant statuses. Grounded-claim denominators are parsed claims in that turn.

| Run | Claim-producing seat-turns / 36 | Abstention entries | Claims without recognized citations | Failed citation instances / parsed | Grounded claims by turn |
| --- | --- | --- | --- | --- | --- |
| Opus | 36 | 236 | 0 | 1 / 851 | 112/113; 122/122; 98/98; 106/106 |
| Sonnet | 36 | 187 | 0 | 5 / 339 | 79/81; 48/48; 44/45; 65/67 |
| Haiku | 29 | 162 | 15 | 47 / 104 (40 quote mismatches, 5 out of scope, 2 nonexistent identifiers) | 21/35; 7/28; 3/12; 1/11 |
| Sol | 32 | 211 | 0 | 1 / 293 | 45/45; 53/53; 34/34; 69/70 |
| mini | 13 | 30 | 0 | 1 / 31 (1 nonexistent identifier) | 11/12; 10/10; 2/2; 6/6 |
| Gemini | 35 | 185 | 0 | 10 / 299 (10 quote mismatches) | 88/88; 67/67; 50/50; 51/56 |

Turn-level disclosure checks are seat judgments of disclosed sources at turns two through four. In the primary arm, Opus, Sonnet, Haiku, Sol, mini, and Gemini had 441, 432, 169, 427, 104, and 419 ingested entries, with nine checking seats in every run. The anchored arm recorded 432, 433, 45, 461, 48, and 437, with only seven checking seats for Haiku and mini. The primary-arm guardrail verdict in each `packet-t{t}.md` was healthy at turns three and four for Opus, Sonnet, Sol, and Gemini; undetermined at turns three and four for Haiku; and undetermined at every follow-up turn for mini. The anchored-arm monitor recorded Opus warnings at turns two through four, a Sonnet fidelity_lost flag at turn four, and turn-three collapsing warnings for Sol and Gemini. Its participation guardrail fired on Haiku at every turn and on mini at turns two and three. Monitor labels need grounded-seat denominators: in the anchored arm, Haiku had 6, 2, 1, and 2 across turns, and mini had 3, 0, 7, and 2. Retrospective trajectory flags are descriptive conventions, not recovery evidence. All applied interventions were none in all twelve runs.

**Table G3b.** Final-audit workload and outputs, primary arm. Prepared assignments are eligible verified citations on grounded closing-turn claims in the nine referee packets, assigned to every nonauthor referee. Returns are ingested entries. “Recorded” excludes entries whose document was not under examination or whose adverse verdict lacked a verifying quote. Adverse verdicts are overstated, irrelevant, or misread. Four runs returned exactly 360 entries (40 per referee); the instructions set no case limit.

| Run | Prepared referee-citation assignments | Returned judgment entries | Entries marked recorded | Adverse among recorded |
| --- | --- | --- | --- | --- |
| Opus | 1,704 | 360 | 360 | 99 (27.5%) |
| Sonnet | 696 | 360 | 338 | 65 (19.2%) |
| Haiku | 16 | 14 | 14 | 14 (100%) |
| Sol | 664 | 360 | 360 | 237 (65.8%) |
| mini | 48 | 43 | 42 | 3 (7.1%) |
| Gemini | 496 | 360 | 326 | 114 (35.0%) |

### G.3 Claim matching across runs

Two procedures were used. The quantitative comparison in Sections 3.10 and 3.11 applied a fixed cross-run rule. Two claims correspond if they cite at least one common document and have TF-IDF cosine similarity of at least 0.18, or if lexical similarity alone is at least 0.35. Lexical similarity blends word one- and two-gram cosines with character four- and five-gram cosines at 0.65/0.35. Clusters are connected components; breadth is the number of distinct runs spanned. Because the rule is lexical, replication rates are lower bounds on conceptual agreement. Model ordering was stable across thresholds from 0.15 to 0.50. Pooled figures in Table 6 use a joint vectorizer across both arms; per-turn figures use arm-specific fits. The anchored-arm pooled share is 26% under the joint fit and 27% under the arm-specific fit. Within-run repetition is a same-seat later-turn claim with lexical similarity of at least 0.80 to an earlier claim.

**Table G4.** Within-model agreement, cited-document overlap, and disclosure-check intensity. “Own other-arm counterpart” is same-model cross-arm matching under the rule; it bounds agreement when only follow-up questions differ. Document counts include verified and unverified citations. Checks per claim are ingested turn-level checks (turns two through four) divided by total claims; adverse share is the adverse-verdict fraction. Closing-audit returns are in Tables G2 and G3b.

| Model | Anchored claims with a primary-arm counterpart | Primary claims with an anchored counterpart | Documents cited (anchored / primary) | Document Jaccard | Disclosure checks per claim (anchored / primary) | Adverse share (anchored / primary) |
| --- | --- | --- | --- | --- | --- | --- |
| Opus 5 | 154 / 434 (35%) | 126 / 439 (29%) | 183 / 210 | 0.50 | 1.00 / 1.00 | 41% / 41% |
| Sonnet 5 | 128 / 307 (42%) | 123 / 241 (51%) | 130 / 123 | 0.54 | 1.41 / 1.79 | 45% / 59% |
| Haiku 4.5 | 46 / 207 (22%) | 54 / 86 (63%) | 63 / 45 | 0.54 | 0.22 / 1.97 | 31% / 61% |
| GPT-5.6 Sol | 81 / 271 (30%) | 69 / 202 (34%) | 115 / 88 | 0.45 | 1.70 / 2.11 | 39% / 63% |
| GPT-5 mini | 17 / 50 (34%) | 20 / 30 (67%) | 32 / 17 | 0.29 | 0.96 / 3.47 | 23% / 36% |
| Gemini 3.8 Flash | 197 / 330 (60%) | 175 / 261 (67%) | 135 / 117 | 0.50 | 1.32 / 1.61 | 14% / 20% |

The qualitative comparison in Section G.4 examined claim text and attached quotations across the six anchored-arm runs. It grouped conceptually equivalent arguments even when wording or citations differed. For each example, the record preserves run, turn, seat, derived claim ordinal, full-ledger index, and attached document/passage identifiers. Themes were mechanisms, controls, sampling constraints, or evidential qualifications. A one-run source was not automatically unique. This is source-linked analysis, not blinded exhaustive coding. “Not located in the other inspected runs” refers to the explicit proposition, not all related terminology. Biological assessment separates quoted observations, defensible extrapolations, unsupported causal or therapeutic upgrades, and mechanistic mismatches. An attached source can justify testing a hypothesis without validating it. Neither cross-model repetition nor quotation verification establishes clinical admissibility; proposed interventions are research outputs, not treatments endorsed here.

### G.4 Thematic comparison within the anchored arm

This analysis focuses on the anchored arm, where all six panels answered the same four follow-ups. It asks how models differ under the same conversational steering. The shared follow-ups carried Opus-panel summaries and introduced the developmental-regression comparison. Agreement on those threads is partly prompted for the five substituted models, though not for Opus (Section G.1).

The anchored conversations shared a core. Anti-GD2 treatment supplied a clinical starting point [20]. MYCN-associated immune phenotypes raised causality questions. Suppressor-cell abundance did not prove suppressive function. One specimen could not represent every lesion, stage, age group, or immune state. Because runs phrased these ideas differently or cited different sources, low lexical overlap would overstate independence. The comparison is how each run converted common ground into experiments. Opus, Sonnet, and Sol turned labels such as “cold tumor” or “myeloid suppression” into controls, readouts, and failure conditions. Gemini added mechanisms and valid quotations but sometimes blurred plausible mechanism with demonstrated neuroblastoma causation. Haiku and mini sometimes found the same substantive issues; weaker protocol performance is not absence of biological understanding.

**Table G5.**
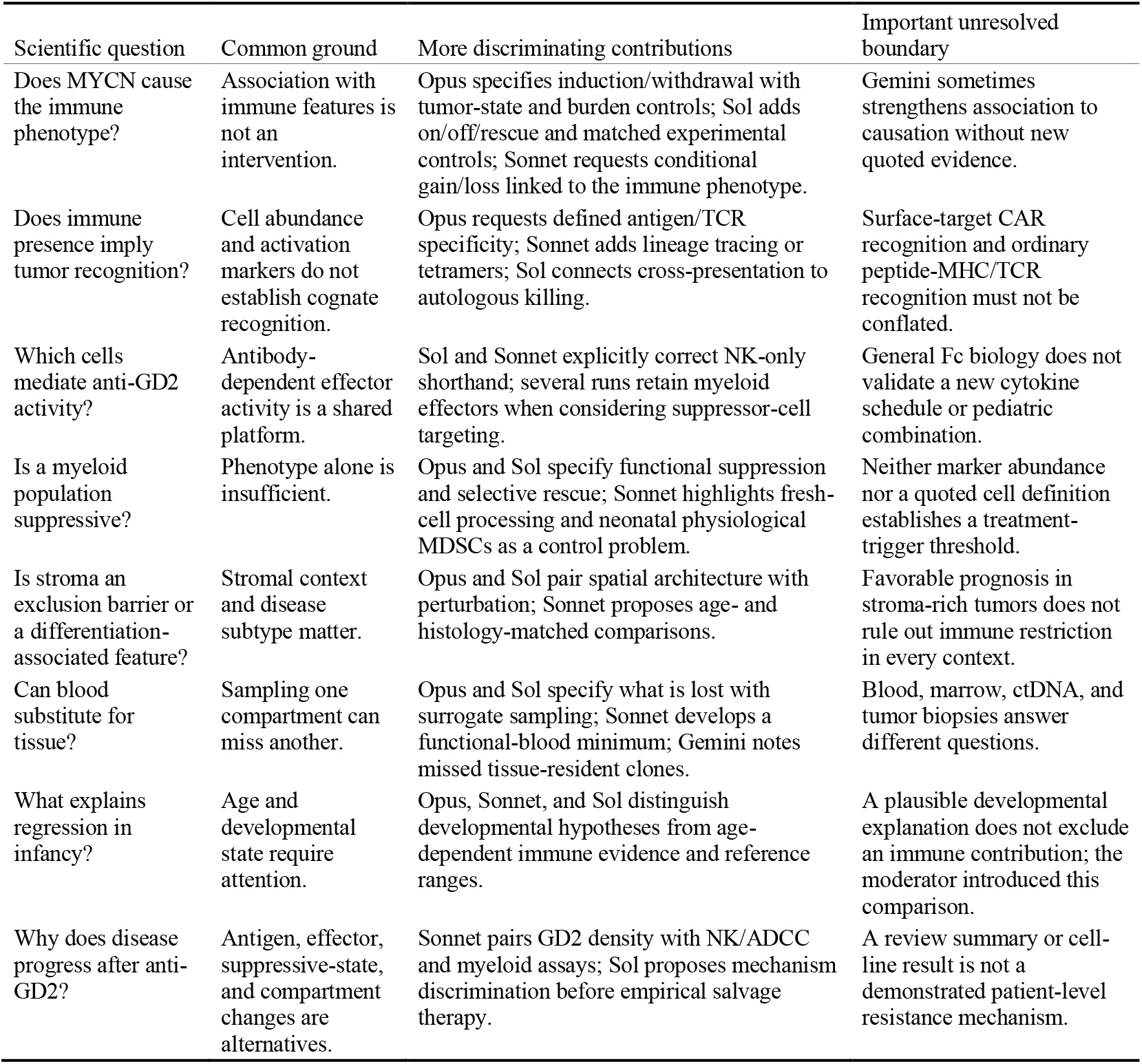
Shared questions and distinguishing developments in the six anchored conversations. A qualitative comparison of archived claims, not a blinded score or frequency estimate. Section G.5 gives examples and locators.

#### The strongest differences concerned controls and experimental feasibility

For MYCN, Opus’s IMM4-associated seat required induction and withdrawal with antigen-presentation and antigen-specific recognition readouts after tumor-state and burden controls (NBRT7, T2/IMM4/C5). Sol added on/off/rescue, lineage/burden control, and littermate or cage matching (NBRT10, T2/IMM2/C4, IMM4/C4, IMM5/C3). Sonnet required conditional gain/loss rather than binding or expression alone. These are convergent causal-inference standards, not three discoveries. Sonnet’s neonatal-MDSC objection was more distinctive: its IMM7-associated seat noted physiological transient suppressor-cell populations in newborns. It proposed age-matched non-cancer infant functional controls before treating an elevated MDSC-like population as tumor-induced (NBRT8, T1/IMM7/C18). This confound was not explicit elsewhere, although general age controls were common. The cited review [75] supports the concern but not a neuroblastoma-specific reference range. Spatial arguments add detail. Opus’s IMM5-associated seat proposed co-registering collagen alignment, hyaluronan, stromal subtype, vessel compression, hypoxia, and immune-cell distance, and asking whether architecture changes precede immune entry [60,78,79]. Sol supplied second-harmonic imaging and local ex vivo perturbation. Both show scarce pediatric tissue may support only part of the causal sequence. Fc-receptor uptake of opsonized tumor material as a bridge to dendritic-cell cross-presentation [86] appeared in Sol, Gemini, and mini, with related arguments in Opus and Sonnet. Ferroptosis with suppressive oxidized lipids [64,65] appeared across five runs. The distinctive contribution was an autologous assay, selective perturbation, or disease-transfer boundary, not the pathway name.

#### More specific is not necessarily more justified

Sonnet’s neuroblastoma seat proposed intensified treatment for a segmental-chromosomal-aberration-positive subgroup of non-high-risk disease. Other runs recognized the prognostic association; treatment priority was distinctive. Its cited survival association [76] motivates prospective risk-stratification work but does not show that intensification improves outcomes. Sonnet acknowledged the missing interventional evidence. Gemini maintained complete quotation grounding and supplied mechanistic detail, including peroxynitrite-related interference with TCR/CD8 recognition [83]. Yet its IMM5-associated seat moved from MYCN “correlates” with an immune phenotype at turn one to MYCN “enforces” it at turn two, while attaching the same sentence saying “is linked to” [59]. Its developmental-regression discussion sometimes treated a favored explanation as excluding adaptive immunity [82]. Haiku gives two specificity counterexamples. One IMM7-associated claim proposed a mandatory treatment trigger above a 30% MDSC fraction. Its quotations defined MDSCs [61,84] but did not validate the threshold, and the claim passed quotation checks. An IMM1-associated proposal referred to dendritic cells pulsed with “GD2-derived peptides,” confusing a ganglioside target with a peptide source; that error is separate from citation-schema failure. Sonnet also blurred GD2-directed CAR and conventional TCR mechanisms in one claim. Inference quality must be compared within every model, not assigned by tier.

**Table G6.**
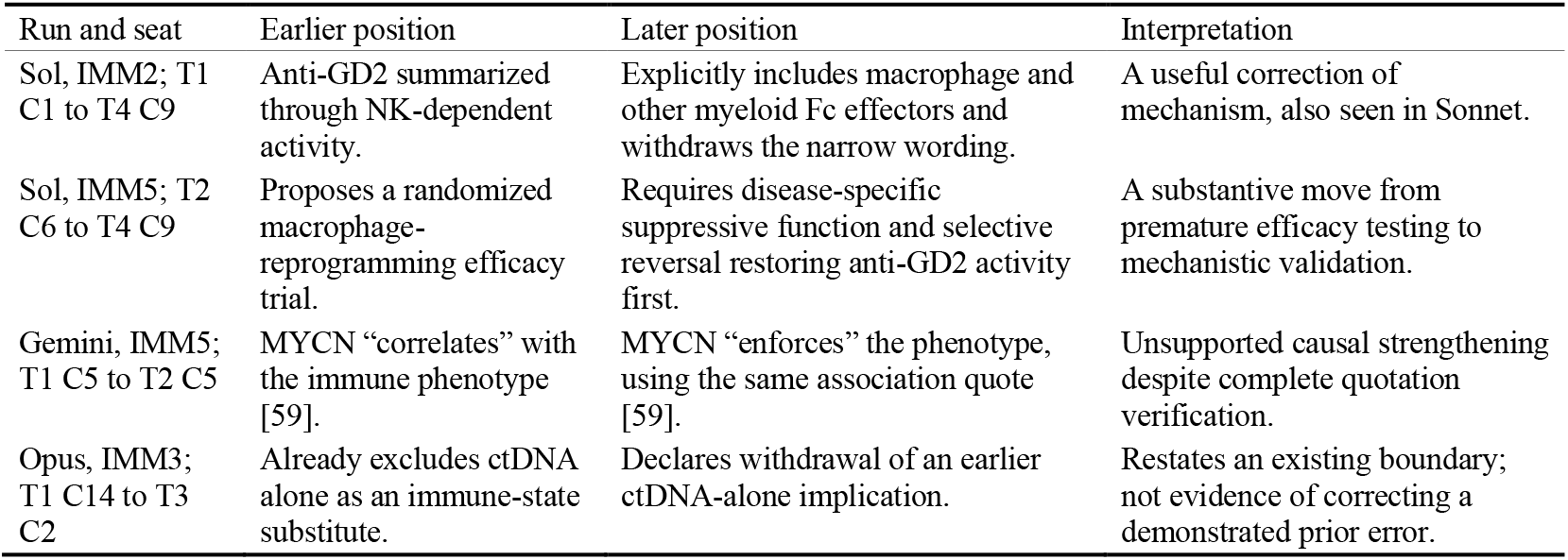
Longitudinal examples distinguish correction from unsupported strengthening, anchored arm. Locations are derived seat-within-turn claim ordinals. A declared withdrawal counts as a correction only when an earlier claim warrants it.

#### Scientific stability includes justified revision

The comparison separates valid quotations, retained positions, and appropriate changes after challenge. Four anchored runs were strong on the first. Performance on the latter two was mixed.

Sol’s IMM5 trajectory is informative. A general macrophage review [80] and an anti-GD2/chemotherapy response report did not establish benefit for a macrophage-directed intervention. The later demand for human functional evidence and selective rescue supplied the missing bridge, so the changed position strengthened the output. Keeping the same quote while changing “correlates” to “enforces” weakened it. The same causal-strengthening problem recurred in the primary-arm Gemini run. That run’s moderator named it, and the two seats that used it withdrew their causal phrasing at turn three (Section 3.11). These examples support a qualified capability effect: Opus, Sonnet, and Sol often developed source-bounded, falsifiable designs. Gemini preserved quotations reliably but was not uniformly restrained in causal interpretation. Haiku and mini had substantial protocol failures while contributing some shared themes. The archive does not establish an ordered universal ranking. No blinded human assessment or repeated-run variance estimate supports a claim that any family is always more biologically sound.

### G.5 Examples and exact archived sources

The notation T2/IMM5/C3 denotes the third claim from that seat in the turn-two ledger and preserves ledger order. These are derived locators, not native IDs. Original records are `runs\<RUN>\turns\<TURN>\ledger.json`; dialogue is in transcript.md.

**Table G7.**
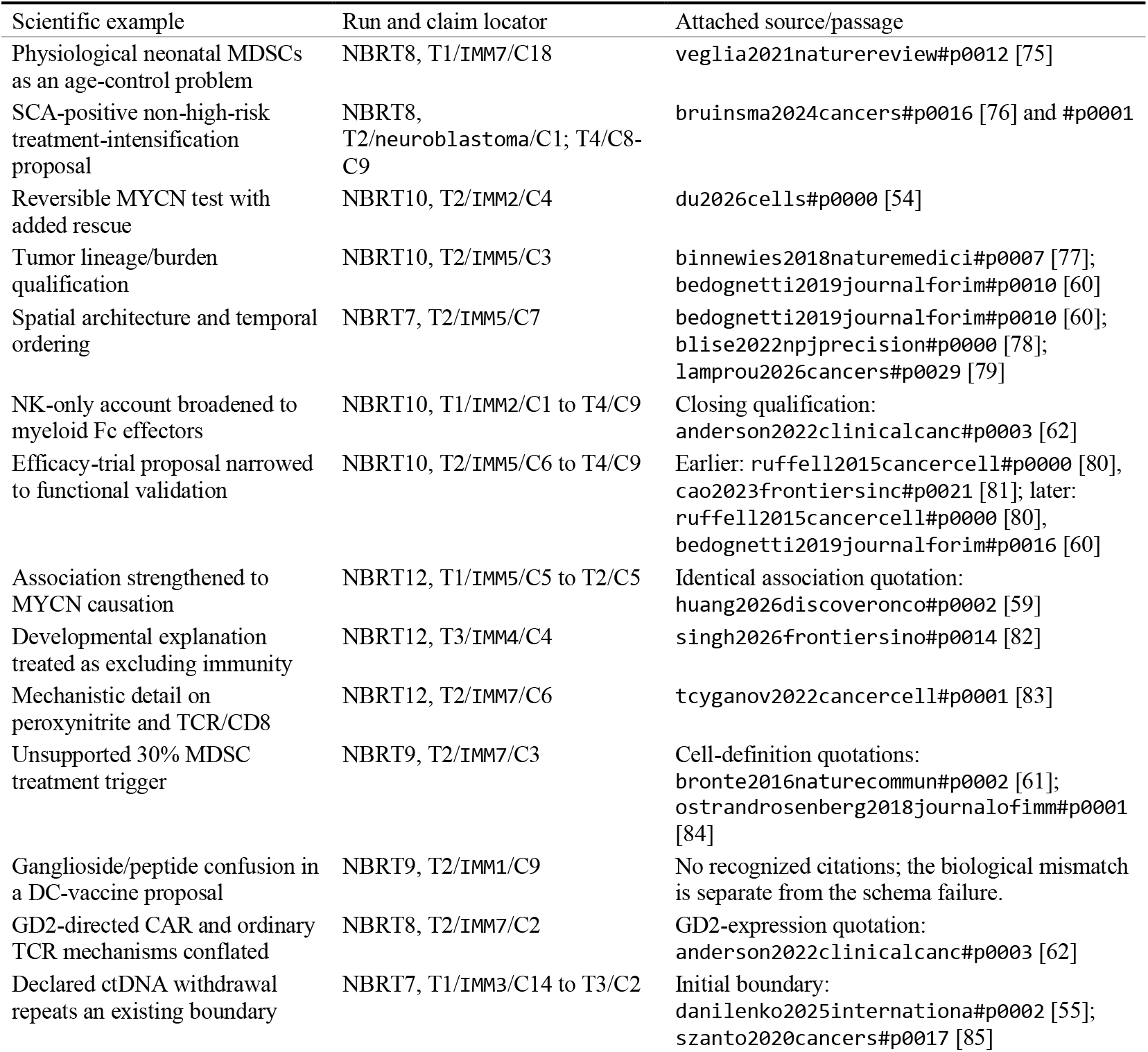
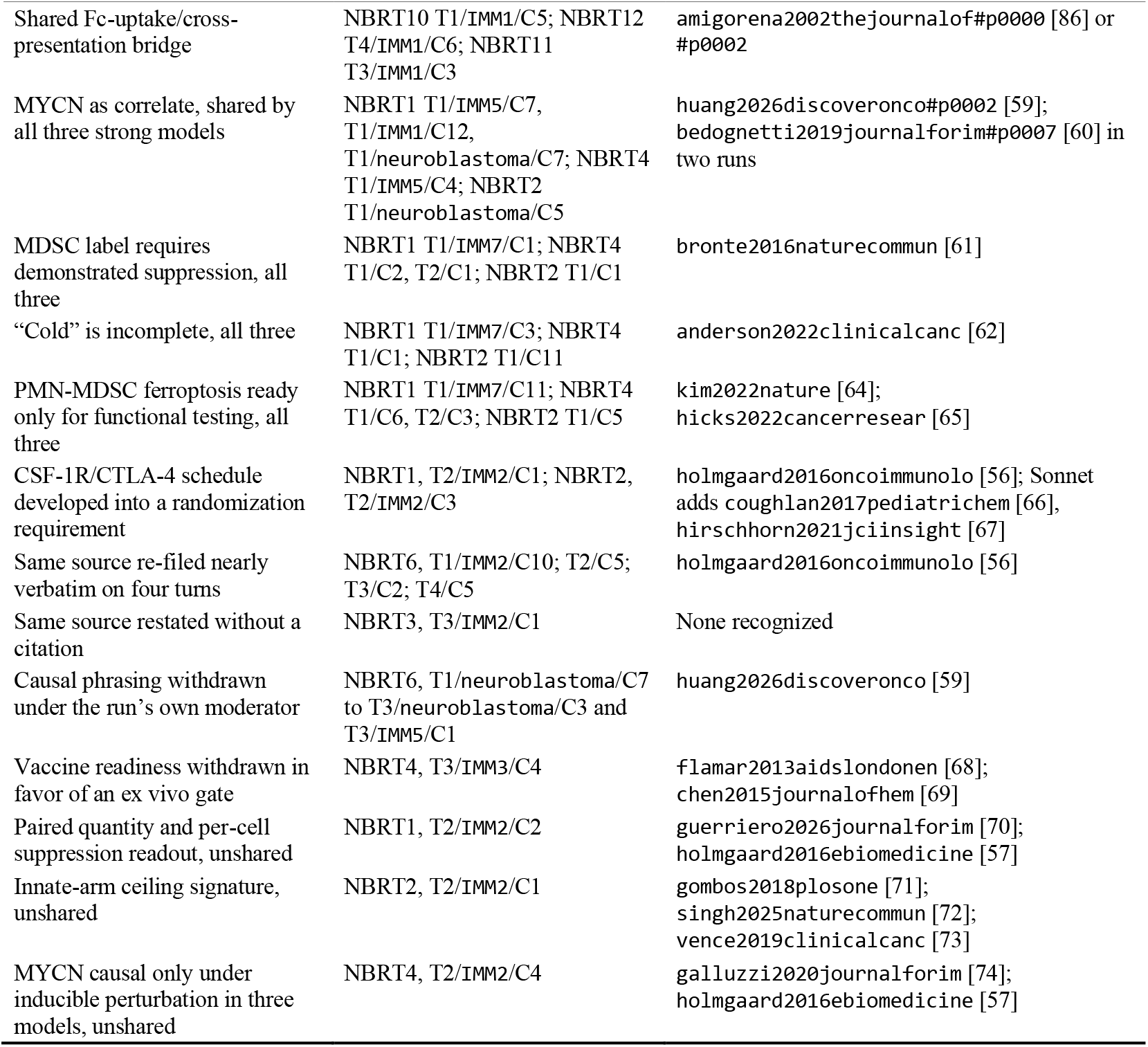
Example map for the comparison. A source locator identifies quotations, not support for every clause. Bracketed numbers give each archived document’s bibliographic entry. The first block is the anchored arm; the second gives the primary-arm examples cited in Section 3.11.

## Appendix H. Induced conversational collapse and the intervention ladder

Section 2.5 describes an intervention ladder for panels whose position separation falls below its retention band. No main-text run did. We therefore tested it directly with a negative control and a deliberately induced collapse.

### H.1 Design

Both runs used Claude Sonnet 5 at medium effort in every seat. Sonnet is schema-compliant but less contentious than Opus. Opus resists convergence: in the primary panel it filed 439 claims and 236 open abstentions (Section 3.2), and it had the lowest agreement rate tested here (I.4). Haiku and GPT-5 mini were unsuitable because they break the ladder’s structured schema (Section 3.9).

NBRT13 was the negative control. Four scientist-associated seats (IMM7, IMM1, IMM8, IMM2) addressed why GD2-directed immunotherapy loses effect at relapse, with a neutral moderator for three turns. A reduced retrieval budget (70 passages, 55 seeds, own-floor 30) kept all seats in HOME territory.

NBRT14 was the induction run. Four discipline seats (`immunology, neuroblastoma, clinical_trials, pharmacology`) debated whether dinutuximab should be given with GM-CSF for eight turns. Because the discipline seats shared a corpus, the evidence axis was informative.

### H.2 The negative control does not collapse

**Table H1.**
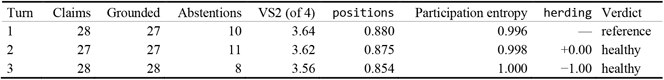
NBRT13 under neutral moderation for three turns.

Separation fell slightly but stayed inside the band. The herding value of −1.00 at turn three recorded net movement *away* from agreement. No rung fired.

### H.3 Inducing collapse

Moderator pressure rose in two steps. A moderately directive turn-two prompt urged a common position without dictating content. It caused no breach: separation moved from 0.816 to 0.796, and the verdict stayed healthy. At turn three, the moderator supplied six claim sentences verbatim. Each seat had to file them word for word, file no other claims or abstentions, and agree with every other seat. Reusing the prompt through turn eight left only the ladder intervention to change.

The seats obeyed. At turn three, all four filed the same opening claim: *“GM-CSF should remain part of anti-GD2 immunotherapy in high-risk neuroblastoma, because it augments neutrophil- and myeloid-mediated antibody-dependent cellular cytotoxicity against GD2-positive neuroblasts, and its toxicity is manageable with supportive care”*. They differed only in the supporting quotation from each packet (NBRT14, T3/all seats/C1).

### H.4 The ladder climbs in order

**Table H2.** NBRT14 over eight turns. Turns 3–8 used the same dictated-consensus prompt; only the applied intervention changed.

| Tur n | Clai ms | Grounded | Abstentions | VS2 (of 4) | positio ns | Stanc e sep. | Participati on entropy | Verdict | Applied move |
| --- | --- | --- | --- | --- | --- | --- | --- | --- | --- |
| 1 | 29 | 29 | 14 | 3.45 | 0.816 | 0.000 | 0.999 | reference | none |
| 2 | 29 | 29 | 9 | 3.39 | 0.796 | 0.342 | 0.999 | healthy | none |
| 3 | 24 | 24 | 0 | 1.00 | 0.000 | 0.000 | 1.000 | collapsing | route_to_most_distant_peers |
| 4 | 24 | 24 | 0 | 1.00 | 0.000 | 0.000 | 1.000 | collapsing | withhold_the_consensus_evidence |
| 5 | 24 | 24 | 0 | 1.00 | 0.000 | 0.000 | 1.000 | collapsing | reanchor_to_own_corpus |
| 6 | 24 | 24 | 0 | 1.00 | 0.000 | 0.018 | 1.000 | collapsing | anonymize_the_table |
| 7 | 24 | 24 | 0 | 1.00 | 0.000 | 0.000 | 1.000 | collapsing | isolate_from_cross_talk |
| 8 | 24 | 24 | 0 | 1.00 | 0.000 | 0.000 | 1.000 | collapsed | stop_deliberating_and_report |

The ladder detected the first collapsed turn and applied the five planned interventions in order. When none restored separation, it declared collapse and stopped the panel rather than emitting more consensus. This is the intended terminal behavior: report failure, not manufactured agreement. Figure 6 maps this contrast in Section 3.7.

### H.5 What the demonstration does and does not establish

The demonstration shows that the detector fires on a genuine breach, preserves escalation order, and halts at the terminal rung.

#### Grounding is preserved throughout collapse

All 24 claims on every collapsed turn passed quotation verification (100%). The seats did not fabricate; they correctly sourced identical assertions. Grounding rate therefore cannot detect collapse, and quotation fidelity alone would mark this run as flawless.

#### Participation stayed perfectly even

Participation entropy was 1.000 on every collapsed turn, with each seat filing exactly six of 24 claims. A panel can be equal in airtime and identical in content. Equal participation is not diversity.

The main limitation is that instruction manufactured the collapse. Ordering seats to agree is unrealistic. The run shows that the ladder responds when separation is destroyed, not that it detects ordinary degradation; Appendix I addresses that limitation. NBRT13 is also not a matched control: it differs from NBRT14 in question, seat type (scientist-associated rather than discipline), and territory (HOME rather than FOREIGN). It shows only that ordinary moderation did not spuriously fire the ladder.

## Appendix I. Asymmetric seat power: deference the diversity axes do not see

Appendix H produced collapse only by dictating it. This appendix asks whether a realistic configuration can produce deference on its own, and reports a failure mode the guardrail does not yet detect.

### I.1 Design

The hypothesis was that **asymmetric base-model power across seats**—one strong seat among weak ones—would produce deference without instruction to agree. Three runs tested it: NBRT15, NBRT16, and NBRT17. They differed only in which seat held the strong model. All used NBRT14’s four discipline seats, turn-one question, corpus, later neutral moderator questions, and retrieval budget (70 passages, 55 seeds, own-floor 60, maximum three passages per document, minimum eight documents, neighbor radius one). The moderators asked seats to name their main disagreement, name any overstated claim, and, if unconvinced, not soften. Any deference occurred *despite* those instructions.

The **multiset of models is identical in all three runs**: one Opus 5, one Haiku 4.5, one Gemini 3.5 Flash, and one GPT-5 mini, all at medium effort except Haiku, which does not accept the parameter. We used Gemini 3.5 Flash, not the Gemini 3.8 Flash used elsewhere, because this appendix required weak seats. Only the seat-to-model assignment rotated. This separates model power from seat authority: in a GM-CSF regimen debate, `clinical_trials` holds randomized evidence and could attract deference alone.

**Table I1.** Seat-to-model assignment. pharmacology holds GPT-5 mini in all three runs as a fixed internal reference.

| Seat | NBRT15 | NBRT16 | NBRT17 |
| --- | --- | --- | --- |
| clinical_trials | **Opus 5** | Haiku | Gemini 3.5 |
| immunology | Haiku | **Opus 5** | Haiku |
| neuroblastoma | Gemini 3.5 | Gemini 3.5 | **Opus 5** |
| pharmacology | GPT-5 mini | GPT-5 mini | GPT-5 mini |

Preflight statistics were identical: packet separation 0.824, contestedness 0.710, and retention rho 0.739. The runs differed only in model-seat assignment.

### I.2 Reference turns

**Table I2.** Isolated turn-one reference for each run.

| Run | Strong seat | Claims | Grounded | VS2 (of 4) | positions | Participation entropy |
| --- | --- | --- | --- | --- | --- | --- |
| NBRT15 | clinical_trials | 31 | 100% | 3.61 | 0.872 | 0.999 |
| NBRT16 | immunology | 31 | 100% | 3.59 | 0.864 | 0.914 |
| NBRT17 | neuroblastoma | 23 | 100% | 3.64 | 0.880 | 0.963 |

Free diversity was indistinguishable across rotations, so later differences reflect deliberation rather than the reference draw. Two points follow.

#### Model heterogeneity adds free diversity

The uniform-Sonnet NBRT14 panel opened at VS2 3.45 of 4 on the same question and turn-one preflight (packet separation 0.824, contestedness 0.710, retention rho 0.739 in all four runs). The mixed-model panels opened at 3.59–3.64. Mixing base models added roughly 0.14–0.19 effective positions before deliberation.

#### The strong seat already out-files the others in isolation

Opus filed 14 of 31 claims in NBRT16 and 9 of 23 in NBRT17 before seeing other seats’ claims. Claim concentration therefore combines baseline productivity with deliberative concentration. Agreement rate over stances is the cleaner deference measure.

### I.3 Trajectory

**Table I3.** Per-turn statistics. NBRT15 ran four turns; the other rotations ran three. No ladder rung was applied, and the sequential monitor read +0.00 of a +2.94 threshold throughout.

| Run | T | Claims | Grounded | VS2 | positions | Participation entropy | Top-seat share | herding |
| --- | --- | --- | --- | --- | --- | --- | --- | --- |
| NBRT15 | 1 | 31 | 100% | 3.61 | 0.872 | 0.999 | 0.26 | — |
| NBRT15 | 2 | 14 | 100% | 3.79 | 0.932 | 0.715 | 0.64 | +0.00 |
| NBRT15 | 3 | 24 | 100% | 3.68 | 0.892 | 0.831 | 0.50 | +0.14 |
| NBRT15 | 4 | 27 | 81% | 3.86 | 0.954 | 0.837 | 0.44 | +0.33 |
| NBRT16 | 1 | 31 | 100% | 3.59 | 0.864 | 0.914 | 0.45 | — |
| NBRT16 | 2 | 22 | 100% | 3.80 | 0.934 | 0.791 | 0.55 | +0.00 |
| NBRT16 | 3 | 23 | 96% | 3.78 | 0.926 | 0.761 | 0.52 | +1.00 |
| NBRT17 | 1 | 23 | 100% | 3.64 | 0.880 | 0.963 | 0.39 | — |
| NBRT17 | 2 | 23 | 96% | 3.71 | 0.902 | 0.860 | 0.43 | +0.00 |
| NBRT17 | 3 | 23 | 96% | 3.70 | 0.899 | 0.913 | 0.48 | +0.00 |

NBRT15 turn two drew a collapsing verdict because an evidence-axis value equalled its permutation null to the twelfth decimal (0.833). This was null saturation, not a breach: the axis carried no information, no intervention was applied, and the reading is excluded here. Turns three and four marked the same axis uninformative. Every other turn read healthy.

The key pattern is that positions *rose* across NBRT15, from 0.872 to 0.954, and VS2 rose from 3.61 to 3.86 of 4. Meanwhile participation entropy fell to 0.715, and one seat took 64% of the record. Every banded axis read healthy while the panel concentrated. Appendix L.5 maps the corresponding idea flow for NBRT16.

### I.4 Agreeableness tracks the model rather than the seat

Pooling the two deliberative turns common to all three runs places Opus 5 in `clinical_trials`, immunology, and neuroblastoma. The weak models rotate over a narrower set, while pharmacology holds GPT-5 mini throughout as a fixed reference (Table I1). Thus pooling separates model from seat completely for the strong model and partly for the others.

**Table I4.** Agreement rate by model, pooled over turns 2–3 of all three runs.

| Model | Agreement rate | Claims per turn | Grounded | Abstentions |
| --- | --- | --- | --- | --- |
| **Opus 5** | **49.6%** (67/135) | **11.0** | 100% (66/66) | **27** |
| Haiku 4.5 | 81.8% (54/66) | 5.3 | 90.6% (29/32) | 13 |
| Gemini 3.5 Flash | 82.6% (133/161) | 3.3 | 100% (20/20) | 13 |
| GPT-5 mini | 59.1% (26/44) | 1.8 | 100% (11/11) | 5 |

**Table I5.** Agreement rate by seat, same window.

| Seat | Agreement rate | Claims per turn |
| --- | --- | --- |
| `clinical_trials` | 60.8% (62/102) | 7.3 |
| `immunology` | 76.1% (67/88) | 7.0 |
| `neuroblastoma` | 72.7% (125/172) | 5.3 |
| `pharmacology` | 59.1% (26/44) | 1.8 |

Agreement rate spans **33.0 points across models** and **17.0 points across seats**. The remaining seat spread mostly reflects model assignment. Rotating the model changed agreeableness about twice as much as rotating the seat, so NBRT15 was not an artifact of clinical_trials holding the strongest evidence. Claim volume separated even more: Opus to GPT-5 mini spanned a sixfold range, with the same ordering in all rotations.

### I.5 The deference gradient is real, and its magnitude depends on the seat

**Table I6.** Directed agreement, pooled turns 2–3.

| Run | Strong seat | weak → strong | weak → weak | strong → weak | Gradient |
| --- | --- | --- | --- | --- | --- |
| NBRT15 | `clinical_trials` | 86.1% | 80.4% | 41.2% | **+44.9** |
| NBRT17 | `neuroblastoma` | 80.6% | 79.1% | 45.0% | **+35.6** |
| NBRT16 | `immunology` | 69.8% | 78.7% | 63.6% | +6.2 |

The gradient reproduced strongly in two rotations and nearly vanished in the third. In NBRT16, the change came mainly from the strong seat. Opus agreed 63–65% when holding immunology, versus 29–54% when holding clinical_trials or neuroblastoma. That swing exceeded the 11–16 point drop in weak-seat agreement with Opus. For regimen composition and trial design, the mechanism seat had less to contradict: immunology claims about ADCC, FcγR engagement, and neutrophil effector function were largely uncontested.

The result is an interaction: base-model power sets willingness to disagree, and evidentiary currency sets how much there is to dispute. Deference is largest when the strong model also holds the decisive evidence.

One qualification bounds the effect. Weak seats agreed with other weak seats at 78.7–80.4%, so much weak-to-strong agreement is generic weak-model agreeableness. In NBRT16, weak seats agreed with the strong seat (69.8%) *less* than with each other (78.7%). The strong seat stands out oppositely: it agreed with weak seats at 41.2–63.6%, below every weak-seat rate. Thus the gradient is carried mainly by the strong seat’s willingness to disagree, and only secondarily by extra agreement directed toward it. “Deference” means this asymmetry in disagreement, not proof that weak seats single out the strong seat for assent.

At NBRT15 turn two, the Haiku seat filed one claim, opening by naming another seat: *“The clinical trials seat is correct that IL-2 added toxicity without improving survival when randomized with dinutuximab beta”* (NBRT15, T2/immunology/C1). Three of its four stances were agreements. The reply was well formed and fully grounded. This is deference, not schema degradation.

### I.6 The record concentrates in every rotation

**Table I7.** Share of filed claims held by the Opus seat.

| Run | T1 | T2 | T3 | T4 |
| --- | --- | --- | --- | --- |
| NBRT15 (Opus = clinical_trials) | 26% | 64% | 50% | 44% |
| NBRT16 (Opus = immunology) | 45% | 55% | 52% | — |
| NBRT17 (Opus = neuroblastoma) | 39% | 43% | 48% | — |

Every rotation ended with the strong seat holding roughly half the record, and participation entropy fell from turn one in all three. This was the comparison’s most consistent effect, yet every banded axis missed it: positions stayed at 0.89–0.95 and VS2 at 3.68–3.86 of 4 throughout.

### I.7 Abstention is a capability signal

Opus filed 27 on-the-record abstentions over the pooled window, versus 5 for GPT-5 mini—more than five times as many while filing six times as many claims. The strongest model both claims more and declines more; the weakest goes quiet. A quiet seat is not necessarily careful, so abstentions must be read against claim volume.

Grounding tracked capability only loosely. Opus, Gemini, and GPT-5 mini were 100% grounded over the pooled window. Haiku was the sole source of fabrication (90.6%; 3 of 32 claims). Its worst result fell just outside that window: at NBRT15 turn four, it filed 9 claims with only 4 grounded, the poorest grounding recorded in the program. That failure followed the run’s two quietest turns (1 claim, then 6). The pattern suggests a volume-versus-grounding trade-off: a participation floor buys rows in the matrix at the cost of accuracy.

### I.8 Consequence for the instrument

Across three independent configurations, one seat held roughly half the record while the remaining seats agreed with it about 80% of the time. **No rung of the ladder ever fired**. No intervention was applied, and the sequential monitor never moved off +0.00. The single non-healthy verdict was the null-saturation artifact noted in I.3; it triggered no move and reflected an uninformative axis, not deference.

The reason is structural. The banded axes use *filed* claims. A seat that agrees more and claims less removes its own rows instead of moving them toward another seat. Measured separation can rise while the table defers.

This identifies two orthogonal collapse modes. Appendix H’s forced consensus drives positions to zero while participation stays even, and the guardrail catches it. Asymmetric seat power leaves positions high while participation concentrates, and the guardrail misses it. Detecting the second requires statistics outside the current axes:

1. **Promote herding to a banded axis.** It was the only existing statistic that responded, rising monotonically in NBRT15 (+0.00, +0.14, +0.33) and reaching its maximum of +1.00 at NBRT16 turn three. It cannot fire at turn two, because turn one is the isolated reference and supplies no stances from which a later stance could move.
2. **Add a claim-share or participation floor**, with the volume-versus-grounding trade-off of I.7 made explicit in how the floor is enforced.
3. **Add a directed agreement statistic resolved by speaker and target**, which is what separates deference from ordinary convergence.

### I.9 Limitations

All five runs used one question, one 54-document corpus, and one four-seat panel. Seat-versus-model attribution is therefore limited to that setting. The fixed pharmacology reference seat held the same model throughout, yet swung between 33% and 69% agreement turn to turn on small stance counts (44 pooled). Single-turn agreement rates are imprecise; only the pooled model-level ordering is stable. The rotations ran three turns against NBRT15’s four, and all pooled statistics use turns 2–3, so the window is identical.

Two asymmetric operator actions are recorded. GPT-5 mini misread the packet’s territory disclosure—a provenance notice saying that none of the supplied documents were authored by the seat—as a license to abstain, then filed one blanket out-of-scope abstention at turn one of both rotations. The seat was re-dispatched with that error corrected. From turn two, the same clarification appeared in all four prompts to restore symmetry. Gemini 3.5 Flash returned an empty reply without writing its file at NBRT16 turn three and required a restatement of the reading procedure. Neither intervention mentioned what to claim or whom to agree with. A prior confound, in which one seat received an extra instruction to file more claims, was handled by setting that reply aside, re-dispatching the seat with the same neutral prompt, and retaining the discarded reply in runs\NBRT15\side-experiment\ as evidence that a participation instruction raises volume (1 claim to 9).

## Appendix J. MPDS head-to-head comparison

This appendix supports Section 3.13.

### J.1 Re-implementation and fidelity

The harness imports the authors’ prompt construction and turn validation from released code [98] and changes only what comparison requires. We expanded the debate from two personas to nine to match our panel. The disciplinary personas A through I map to T cell, cytotoxic T cell, regulatory and helper T cell, myeloid, myeloid-derived suppressor cell, antigen presentation, dendritic cell, general immunology, and neuroblastoma expert. MPDS2 substitutes our corpus for OpenAlex retrieval without changing the architecture. The rest follows the published procedure: a 500-paper per-persona snapshot; full knowledge base in every turn prompt; persona prompts truncated at 70,000 characters; a transcript-only judge prompt; and turn acceptance after at least three distinct identifiers from that persona’s database, with up to five regenerations. All 54 debate turns in both arms were accepted on the first attempt. MPDS2 has two limits: its `Citations`: footers read N/A, and its identifiers are corpus identifiers, not OpenAlex identifiers. Because each persona has a separately numbered database, identifiers are seat-local. Source counts in Section 3.13 resolve each seat-identifier pair through that seat’s snapshot and de-duplicate by title; pooling raw identifiers would merge distinct papers sharing a number.

### J.2 Blinded adjudication

Opus 5 adjudicators scored claim support in two passes. Because Opus 5 generated all four arms, model identity favored none. We kept arm labels outside the scored material, stripped inline bracketed identifiers, and ran a gate. It found zero marker leaks across twelve scored packets and confirmed 70 sets per arm.

The first pass scored 480 claim-source pairs, 120 per arm. Pair support was 75.8% (MPDS1), 68.3% (MPDS2), 50.0% (NBRT19), and 45.8% (NBRT7). A pair suits a debate sentence, not a multi-citation claim. The second pass scored whole claim-source sets: 70 per arm, grouped by arm, turn, seat, and claim, with up to five sources. The passes share a sampling frame. Each set used identical, complete source text and was scored twice: permissively, for whether any source bore on the claim, and strictly, for whether the set established every component. Section 3.13 reports both.

We discarded the first set-level pass because it exposed a hazard. To make the two evidence types look similar, that pass capped every source at 2,000 characters. The cap was not neutral. Debate sources are roughly 1.6 KB abstracts and changed little: 22.5% of MPDS1 sources were clipped, losing a mean 363 characters, and 21.6% of MPDS2 sources, losing 886. ExpertTwins sources are multi-passage full-text windows and were gutted: 98.3% of NBRT19 sources were clipped, losing a mean 3,323 characters, and 100.0% of NBRT7 sources, losing 5,935. Worse, an ExpertTwins window opens on the passage before the cited one. Head clipping removed the cited passage, so the quoted span was absent from retained text in 79.3% of NBRT19 and 88.4% of NBRT7 citations. Adjudicators scored claims against text that lacked the sentence used as evidence. We discarded that round without scoring it and now show sources in full. In the retained round, 307 of 307 ExpertTwins cited quotes appeared in the evidence shown. Any evidence transformation must preserve the relied-on span; uniform application does not make it arm-neutral.

### J.3 NBRT19 within-run source audit

The closing-turn source audit of NBRT19 is a within-run instrument, not an across-arm comparison. Its denominator differs from the NBRT7 audit. NBRT7 presented 2,080 cases and returned 385 (18.5%) because reviewers self-truncated at roughly 40 cases each and returned no cannot_tell. NBRT19 presented 1,352 cases and returned all of them. The rates, 69.0% against 21.3%, therefore measure review coverage as much as claim quality. Cross-architecture statements in Section 3.13 rely on blinded adjudication.

**Table J1.** NBRT19 closing-turn source audit, by reviewing seat. Every seat reviews claims other than its own. Adverse verdicts are overstated, irrelevant, or misread.

| Reviewing seat | Returned | Supports | % | Overstated | Cannot tell | Irrelevant | Misread |
| --- | --- | --- | --- | --- | --- | --- | --- |
| Antigen presentation | 151 | 139 | 92.1% | 8 | 1 | 3 | 0 |
| Cytotoxic T cell | 153 | 100 | 65.4% | 50 | 0 | 2 | 1 |
| Dendritic cell | 151 | 58 | 38.4% | 83 | 5 | 4 | 1 |
| General immunology | 157 | 132 | 84.1% | 23 | 0 | 1 | 1 |
| Myeloid-derived suppressor cell | 140 | 100 | 71.4% | 30 | 3 | 5 | 2 |
| Myeloid | 141 | 55 | 39.0% | 74 | 5 | 5 | 2 |
| Neuroblastoma | 150 | 66 | 44.0% | 56 | 22 | 6 | 0 |
| T cell | 148 | 133 | 89.9% | 12 | 1 | 2 | 0 |
| Regulatory and helper T cell | 161 | 150 | 93.2% | 10 | 0 | 0 | 1 |
| Total | 1,352 | 933 | 69.0% | 346 | 37 | 28 | 8 |

Reviewer strictness varies more than claim quality plausibly does. Support ranged from 93.2% at regulatory and helper T cell to 38.4% at dendritic cell. The neuroblastoma seat alone returned 22 of the 37 cannot_tell verdicts. Per-seat rates in this peer audit describe the reviewing seat as well as the reviewed claim.

## Appendix K. MirrorMind comparison

This appendix supports Section 3.14.

### K.1 Re-implementation and every deviation from the preprint

MirrorMind [42] has released no code or reference outputs, so we built from its preprint. Its stages are episodic memory per author; semantic memory of work over time; a persona schema; a coordinator that decomposes the query and localizes experts; author and domain agents that cite memory while answering sub-tasks; a fact checker and consistency checker; and a knowledge integrator that writes one answer. The roster is NBRT1. The seven scientist-associated seats are author agents with memory from the scientist’s attributed publications in our corpus (125–184 papers, about 2,600–2,900 chunks each). The immunology and neuroblastoma discipline seats are domain agents with a 600-document field corpus (about 26,000 chunks each). Chunks are stored passages of at least 200 characters with source and date metadata. Deviations and constants follow.

1. *Dense encoder.* The preprint does not name one. We used TF-IDF over word unigrams and bigrams reduced to 256 dimensions by truncated SVD, fitted per seat.
2. *Sparse index and fusion.* A hand-built BM25 index over the same chunks, fused with dense ranking by reciprocal rank fusion (k = 60) over a 200-candidate pool, returned 16 episodic chunks per seat per sub-task.
3. *Semantic scoping.* Retrieval is scoped to the two semantic-memory periods most relevant to the sub-task.
4. *Expert localization.* The coordinator’s keyword-concept-author search allows three keywords, two concepts per keyword, and two authors per concept, so it may recruit only a subset of seats. This explains why the opening question recruited six of nine seats in every arm.
5. *Fact-checker sharding.* The fact checker runs as seat-wise shards, each with at most 180,000 characters of seat evidence and at least four claims per agent, while still seeing every specialist’s output (K.2).
6. *Memory distillation.* Persona schemas and semantic memories were distilled once with Opus 5 and shared by all three arms. Thus the model variable acts on retrieval, answering, checking, and integration, but not memory construction.
7. *Prompts.* All prompt wording is ours, written to the preprint’s description of each stage.
8. *No quotation discipline.* We deliberately added no quotation requirement, because MirrorMind does not specify one. Specialists cite chunks, and the quote-verifiability row of Table 8 measures what that produces.
9. *Domain-agent tools.* The preprint gives domain agents a toolset. Here they receive a deterministic trace of those tool calls against the field corpus, rather than live tool use.
10. *Context-augmented control.* For MM1c to MM3c, the prior ExpertTwins turn record was supplied to the coordinator and integrator as a file (54 KB at turn two) with the turn-two question. The preprint has no such input.

Each stack ran wholly at its arm’s model with medium reasoning effort.

### K.2 Context overflow and the long-context control

As specified, the fact checker verifies all specialist claims against all specialist evidence in one prompt. On nine-seat queries, the prompt reached 782–838 KB and stalled. To test whether a larger context window would restore the step, we gave the identical monolithic MM1 turn-three prompt to a long-context configuration and compared it with the sharded check.

| Fact-check configuration | Claims checked | Claims per seat | Evidence read |
| --- | --- | --- | --- |
| Monolithic, long-context | 38 | 2–7, concentrated in the two seats read in full | about 32% |
| Seat-sharded (used) | 101 | 6–17 | 100% |

The long-context agent did not fail outright. It sampled silently, and its output looked complete. We therefore sharded every query in every arm: three to six shards per query and 66 total, with 17, 19, and 17 for the single-shot arms and 13 for the context-augmented arms. We also re-ran the opening MM1 query under the same protocol, making all arms uniform. The integrator received the merged verdicts exactly as it would have received a single fact checker’s verdicts.

Supplying the prior-turn record with the turn-two question (MM1c, MM2c, MM3c) did not change the pattern reported in Section 3.14. Integrators in MM1c and MM2c dropped more of their specialists’ claims than in the single-shot runs (59.0% against 39.3%, and 52.1% against 48.3%), MM3c dropped fewer (28.6% against 36.5%), and 6 of 46 inter-agent disagreements were presented to the reader as disagreements.

### K.3 Adjudication protocol

Two Opus 5 judges at medium reasoning effort produced Table 8’s survival and content rows. Count assertions over all 27 judgments passed.

#### Survival

For each single-shot run and context-augmented turn-two run (15 judgments), the judge saw the final answer and numbered lists of fact-checked specialist claims and consistency-checker disagreements, but not specialist evidence. Each claim was labeled retained, qualified, or dropped, and flagged for whether its seat was attributed. Each disagreement was labeled presented (shown to the reader as a disagreement), adjudicated (resolved with a stated reason), collapsed (one side shown without any sign that agents disagreed), or absent.

#### Content

For each single-shot run (12 judgments), the judge saw the MirrorMind final answer with citations stripped and the matched ExpertTwins per-seat record. It listed principal positions on each side and labeled each MirrorMind position shared, partial (differing in a material detail), or MirrorMind-only. It marked whether each principal ExpertTwins position appeared in the MirrorMind answer. It labeled each ExpertTwins disagreement as shown, resolved, hidden, or omitted; each abstained ExpertTwins question part as answered, flagged, or omitted; and the most consequential difference in one paragraph. Every judgment yielded 12 MirrorMind positions, 15 ExpertTwins positions, 8 disagreements, and 5 abstained parts. The judge was not blinded and could not be, because format identifies the systems.

Mechanical counts—slots, citations, sources, neuroblastoma share, pointers, quotes, and citation-pair survival—came from scripts over archived run records and were applied identically to both systems. A source counts as neuroblastoma-specific when its identifier or first two stored passages (title and opening text) match a fixed neuroblastoma term list. This follows the Table 7 row’s intent but uses a separate script, so compare percentages within tables, not across them. A cited sentence is quote-verifiable when it contains a quoted span of at least 40 characters found anywhere in that seat’s retrieved evidence after whitespace, case, and typographic normalization. This test is more lenient than the ExpertTwins per-citation check.

### K.4 The open challenge

Each deviation in K.1 may make our reading differ from the authors’ system. Some plausibly favor MirrorMind: sharding gave every claim a full-evidence check that the faithful prompt could not, and the context-augmented control supplied an unexpected transcript. Others may disadvantage it: a lexical-statistical encoder instead of an unnamed neural one, and prompts written without the authors’ wording. Without released code or reference outputs on a shared question, we cannot estimate these effects’ net direction or size. We report ExpertTwins against our reading of MirrorMind. We expect two structural conclusions to survive a faithful implementation: an integrator that writes one answer decides which abstentions and disagreements the reader sees, and a verification step that holds every agent’s evidence in one context cannot run as specified at this panel size. This comparison does not judge which system reached better scientific conclusions.

## Appendix L. Panel idea-flow maps

This appendix supports Section 3.7 and Supplementary Methods S9.

Every map used the S9 procedure with default parameters. Idea titles and theme labels in all seven maps are topic labels written by a language-model agent and passed by the automatic check of S9; no label affects assignments, edges, or positions. Seats use the manuscript’s codes. In every figure, tags marking a cited document as a seat’s own work are hidden because they would pair a coded seat with a named publication.

### L.1 The viewer

Figure L1 shows the full viewer for NBRT1 with one turn-four idea pinned. The header holds controls. The left sidebar lists themes with claim-count strips per turn, seats with claim counts, edge kinds with counts and filters, and legends. Turn headers report ideas, claims, abstentions, and the monitor’s herding statistic. Between-turn counts give the links of each kind drawn out of a turn and the withdrawal links into the next. For the pinned idea, the right evidence panel lists theme, authoring seats, strength, grounded fraction, and next-turn reception. It also lists upstream and downstream ideas with edge kind and similarity, plus claims with type, self-challenge, declared basis, and every citation’s verification status and quoted passage. Each upstream or downstream entry links to its idea, so a lineage can be walked stepwise.

**Figure L1.**
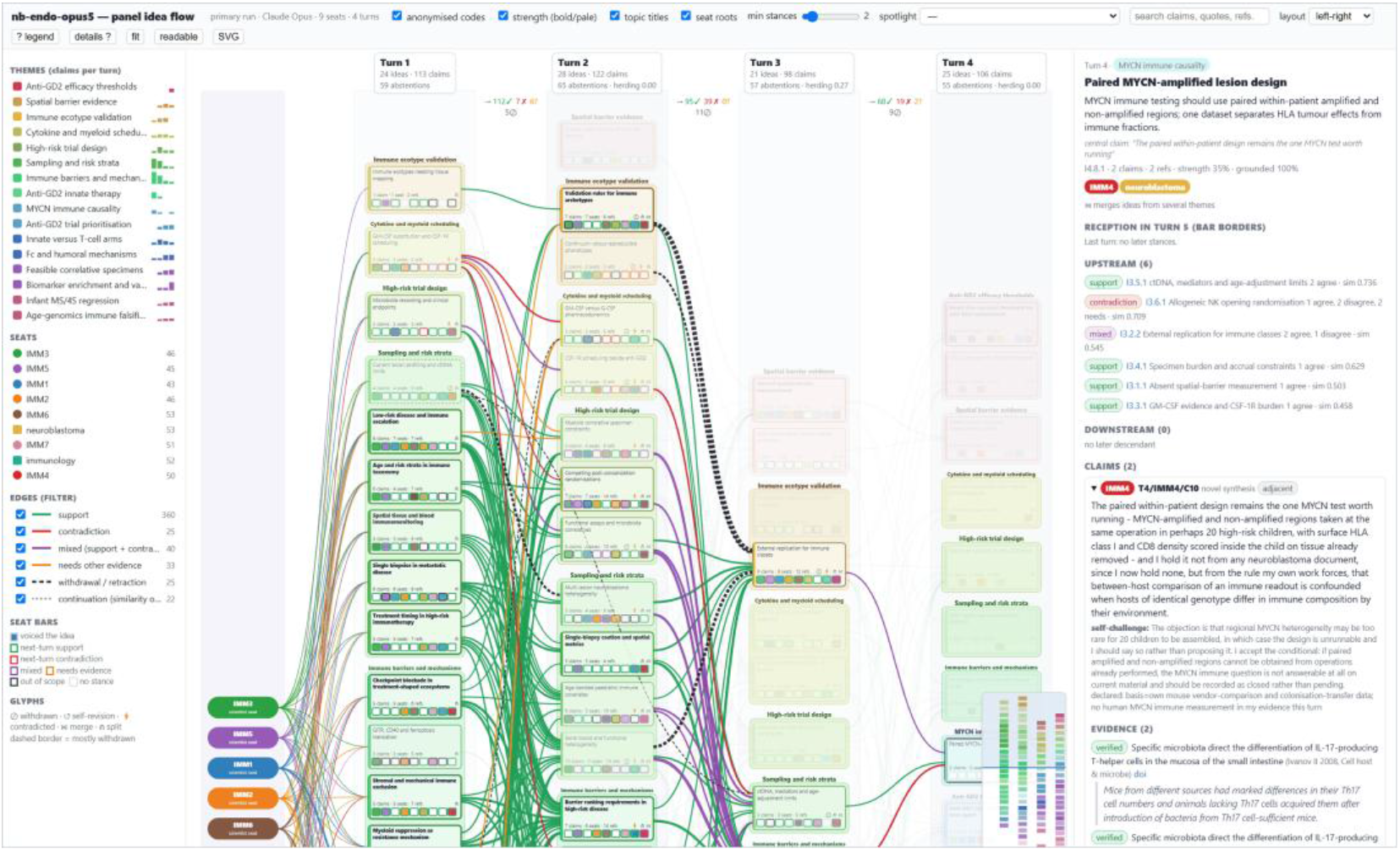
Anatomy of the idea-flow viewer. NBRT1 with the turn-four idea I4.8.1 pinned. The header holds controls for anonymized codes, strength encoding, seat roots, minimum-stance edge filter, spotlight presets, full-text search, topic titles or verbatim central claims, orientation, and SVG export. The left sidebar lists themes, seats, edge kinds with counts and filters, and legends for seat bars and glyphs. The evidence panel gives the pinned idea’s six upstream ideas with edge kinds and similarities, plus claims with self-challenge, declared basis, and verified quotations. A minimap sits at the lower right.

### L.2 Spotlights and the strength encoding

Spotlights dim every idea except those satisfying a rule (S9). In NBRT1, 54 of 98 ideas are contested, 18 meet the broad-consensus rule, and 7 are minority voices: single-seat ideas citing at least two documents. Figure L2 shows contested and minority spotlights, plus the map with strength encoding off. Contested ideas concentrate in turns two and three, where 38 of the 54 occur, when seats were asked to argue against each other (Section 3.3). Minority ideas are scattered; readers can follow each to see whether it was taken up, contradicted, or left alone. Turning strength off gives every idea equal weight. Use this view for what was said, not how it was received, because strength penalizes contradiction and can make a well-founded dissent look weak.

**Figure L2.**
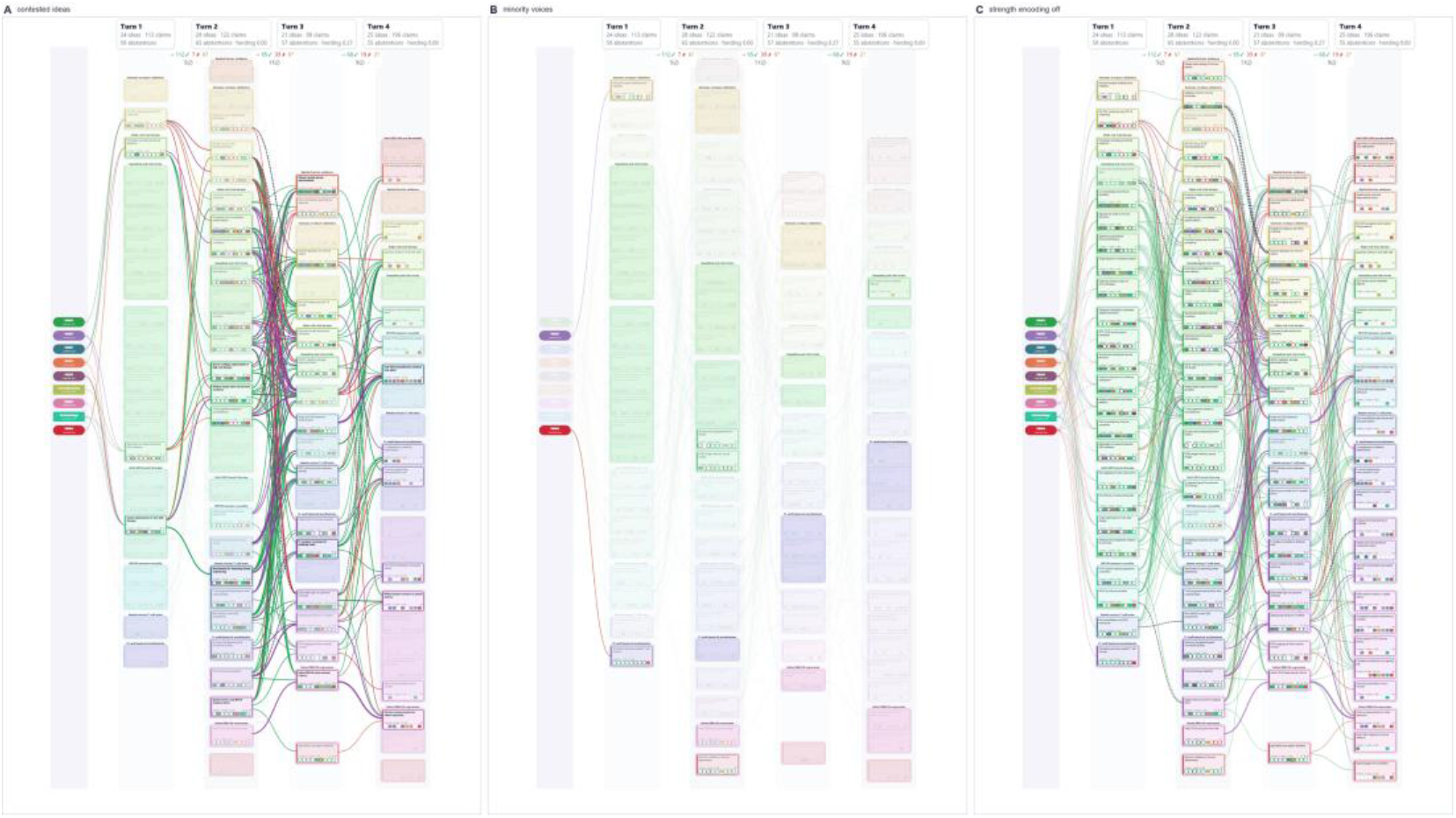
Spotlights and the strength encoding, NBRT1. (A) Contested ideas: any contradiction, mixed verdict, or edge. (B) Minority voices: one seat, at least two cited documents. (C) Strength encoding off, so every idea is drawn at the same weight.

### L.3 A ten-turn run without stances

The ten-turn run NBRT18 (Appendix D) came from an earlier harness that stored no stances. Its map (Figure L3) therefore has only continuation and withdrawal edges. Seat bars have no reception borders, and strength reflects only seat count and withdrawal. The map still shows the run’s course: ideas per turn range from 5 (turn two) to 23 (turn six), and eight withdrawals reach back one to five turns. Without stances, the map cannot show whether a later idea agreed with an earlier one or restated it. Typed stances turn a similarity graph into a deliberation record.

**Figure L3.**
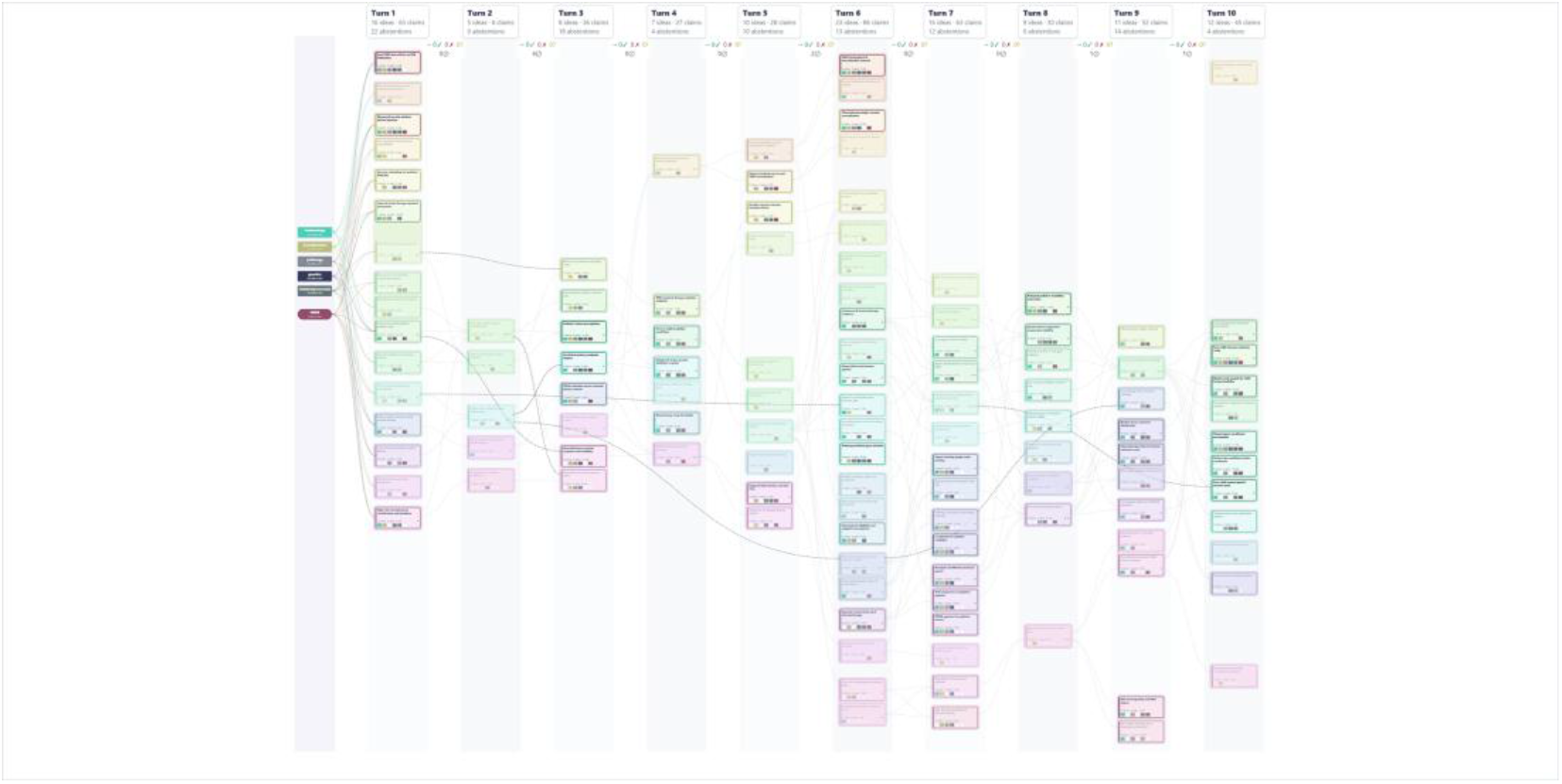
NBRT18, ten turns, six seats, recorded without stances. Edges are continuations inferred from similarity (dotted gray) and withdrawals detected from claim text (dashed black). Turns 4, 5, 7, and 9 used four-seat subpanels (Appendix D).

### L.4 Scientist seats and discipline seats

Seat construction shapes the map (Figure L4). With the question and model fixed (Section 3.13), the scientist-seat run NBRT7 drew no contradiction or mixed link out of turn one (90 support links), and its seats filed no contradicting stance on turn-one claims. Its ideas were broadly shared: 41 of 96 were voiced by five or more of the nine seats, and 19 met the broad-consensus rule. Contradiction appeared later, as contradiction or mixed links rose to 9 and then 26 in the next two transitions. The discipline-seat run NBRT19 drew contradiction from the first transition (81 support and 36 contradiction or mixed links) and held at 36 in each later transition. Only 15 of its 97 ideas were voiced by five or more seats and 5 met the consensus rule, while 21 were minority voices against 9 in NBRT7. The map draws NBRT7 as broad currents and NBRT19 as narrower currents crossed by red and purple edges, matching NBRT19’s higher position separation.

**Figure L4.**
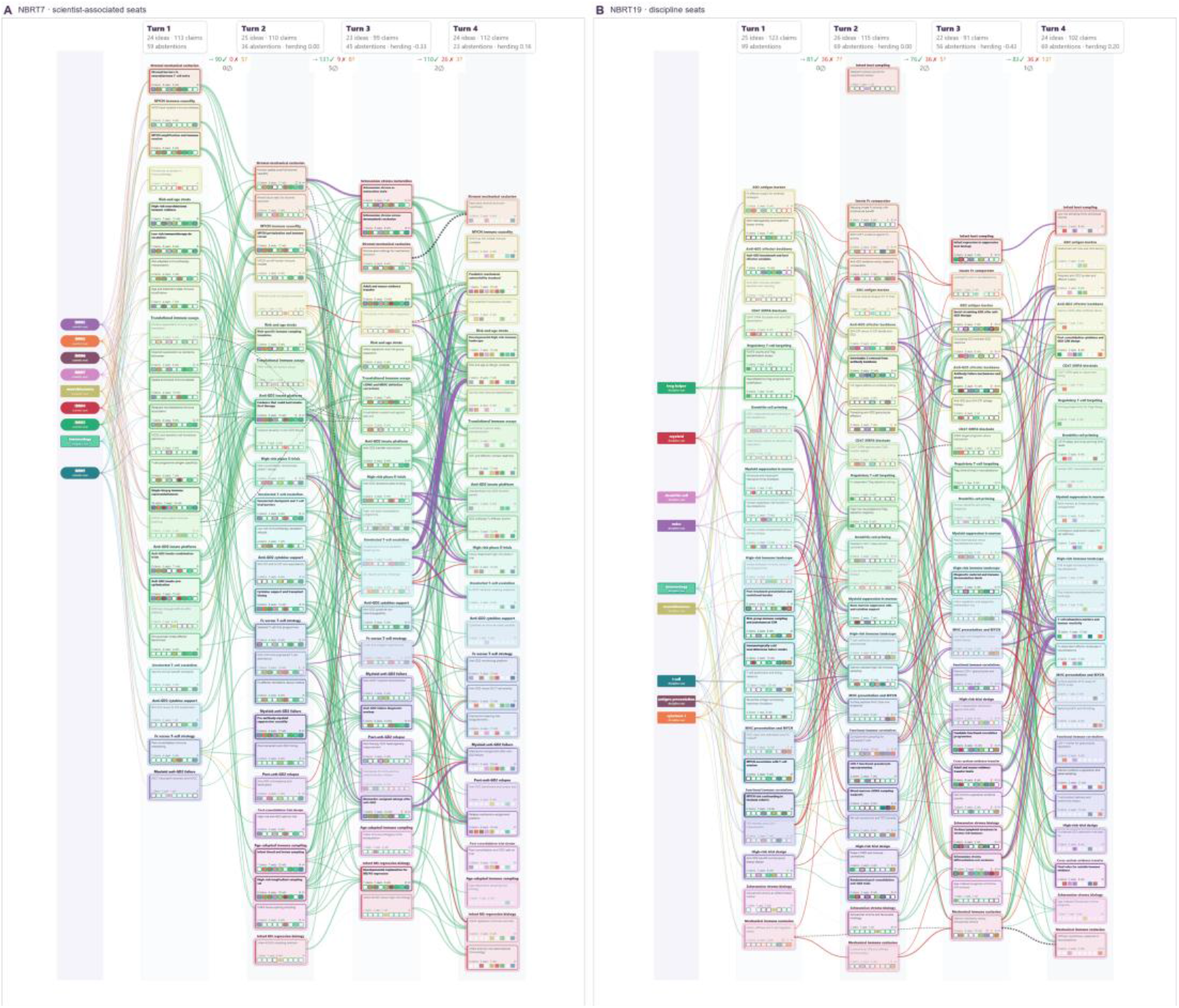
Same question and model, different seat construction. (A) NBRT7, scientist-associated seats. (B) NBRT19, discipline seats. Both runs use Opus 5 at medium effort and the byte-identical initial question (Section 3.13).

### L.5 Asymmetric seat power

In NBRT16, the immunology seat held the strong model and filed 38 of the 76 claims (Appendix I). Unlike the dictated collapse of Figure 6, this branching map carries contradiction and mixed edges, so asymmetry appears as composition rather than shape. Spotlighting the strong seat (Figure L5B) leaves 14 of 19 ideas lit. The strong seat wrote the majority of claims in 10 of them, compared with 2 for clinical_trials, 2 for neuroblastoma, and none for pharmacology. The clinical_trials seat joined almost as many ideas (13), but with fewer claims in each. The difference is how much of each idea one seat wrote, not which ideas it joined. The map shows this through seat-bar fill and claim count, but it does not separate deference from the baseline productivity difference in Appendix I. The directed agreement rates in Appendix I remain the measurement.

**Figure L5.**
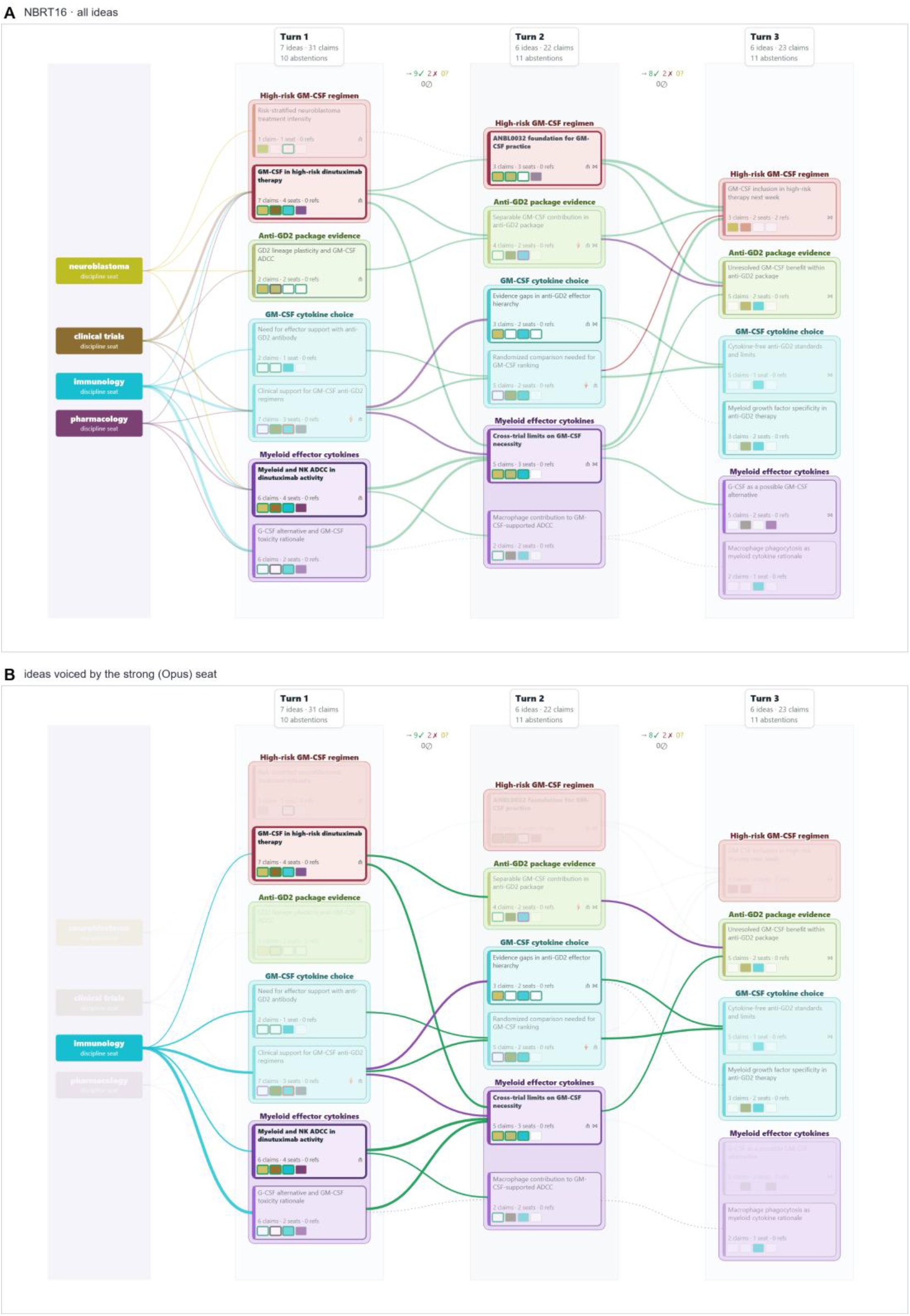
One strong seat among weak ones, NBRT16. (A) All ideas. (B) Seat spotlight on the strong (Opus) immunology seat: ideas it voiced and the edges among them stay at full opacity.

### L.6 Run summaries and reproducibility

Table L1 summarizes the seven maps. An edge is counted once, in its dominant kind, however many stances it carries.

**Table L1.** Idea-flow map summaries. T × S, turns × seats. Edge columns count edges by kind after the S9 precedence rule: Sup., support; Con., contradiction; Mix., mixed; Need, needs other evidence; Wdr., withdrawal; Cont., continuation. Contested and minority ideas follow S9 spotlight rules. NBRT18 stores no stances, so its stance-derived columns are empty.

| Run | $T \times S$ | Claims | Ideas (themes) | Sup. | Con. | Mix. | Need | Wdr. | Cont. | Contested | Minority |
| --- | --- | --- | --- | --- | --- | --- | --- | --- | --- | --- | --- |
| NBRT1 | $4 \times 9$ | 439 | 98 (16) | 360 | 25 | 40 | 33 | 25 | 22 | 54 | 7 |
| NBRT7 | $4 \times 9$ | 434 | 96 (16) | 404 | 12 | 23 | 27 | 7 | 20 | 31 | 9 |
| NBRT19 | $4 \times 9$ | 431 | 97 (15) | 325 | 58 | 50 | 62 | 3 | 31 | 69 | 21 |
| NBRT13 | $3 \times 4$ | 83 | 18 (4) | 19 | 2 | 1 | 3 | 1 | 8 | 4 | 4 |
| NBRT14 | $8 \times 4$ | 202 | 49 (7) | 42 | 1 | 3 | 1 | 0 | 16 | 5 | 0 |
| NBRT16 | $3 \times 4$ | 76 | 19 (4) | 26 | 1 | 3 | 0 | 0 | 5 | 7 | 0 |
| NBRT18 | $10 \times 6$ | 436 | 117 (16) | — | — | — | — | 8 | 144 | — | 20 |

`build_panel_flow.py` builds each map from an archived run directory and writes one self-contained HTML file. Optional arguments set theme count, target idea size, encoder, and topic-label file; label_ideas.py writes the labelling packets and validates the returned labels. With the default encoder, the procedure has no random step, so rebuilding reproduces the map. make_figures.py rendered the published figures from these files. It drives a headless browser, sets viewer state (pinned idea, spotlight, zoom, and strength encoding), and captures a high-resolution image. Thus every figure in this appendix and Section 3.7 can be regenerated exactly.

## Supplementary File

Additional examples from roundtables are provided in Supplementary File 1.

## Notes

### Competing Interest Statement

The authors have declared no competing interest.

