## Supplementary File 1 for "ExpertTwins: Auditable Long Horizon Scientific Panels through Grounded Expertise"

*Supplementary File 1: Supplementary Examples and Annotated Excerpts from the Roundtable Runs*

**Ali Foroughi pour, Abbas Shirinifard**

This document accompanies the main manuscript. It exists because the aggregate tables in the paper report *rates* — claims per turn, abstentions, quote-verification pass rates, disagreement counts — and a rate cannot show a reader what the underlying behavior actually looked like. Every excerpt below is copied verbatim from the run artifacts, except that researcher names are replaced by seat codes in square brackets. No excerpt is paraphrased or reconstructed.

#### S1. How to read these examples

**Locators.** Each excerpt is labeled `run · T<turn>/<seat>/C<n>`, where `C<n>` is the *n*-th claim filed by that seat in that turn, in ledger order. This is the same convention used in the main text, so any excerpt here can be traced back to `runs/<run>/turns/<turn>/ledger.json` in the released artifacts.

**What "verified" means.** A claim's citation carries a `status` field written by the post-hoc checker, not by the agent. `verified` means the quoted span was found character-for-character in the cited document's extracted text. `quote_not_found` means the document exists and the seat was permitted to cite it, but no passage matches the quoted string — the quote was altered, compressed, or composed. `out_of_scope_invented` means the cited identifier does not correspond to any document in the library at all. Seats never see these verdicts; they are applied after the turn closes.

**Ellipsis and truncation.** Long claims are truncated with `[...]` where the omitted text does not bear on the point being illustrated. Quotes are never truncated mid-sentence without marking. Original spelling, including the British forms some seats adopt, is preserved.

**Reference numbering.** The numbered references at the end of this document are local to it and do **not** correspond to the reference numbers in the main manuscript. Each entry is keyed to the `doc_id` used in the run artifacts so the two can be cross-walked mechanically.

**Naming.** Seat identifiers such as `IMM2` or `IMM4` name a *configured seat* built from a public scholarly corpus. They are not the person, do not speak for the person, and their outputs were not reviewed or endorsed by the person. Discipline seats (`immunology`, `neuroblastoma`, `pathology`, `genetics`, `rhabdomyosarcoma`) carry no individual attribution at all.

**Own-work and read status.** The platform tags every packet document with its relation to the seat that receives it: `own` for a work the seat's source scientist authored, `read` for a work drawn from the referenced, collaborator, citing, or topic-canon neighborhoods. Both tags are recorded for every run. For scientist-associated seats, this document never marks an individual cited document as the seat's own work, because pairing a coded seat with a specific cited work and an `own` tag would re-identify the scientist through that work's author list. Aggregate statements that do not name a document — for example, that a given failure class consisted of seats mis-quoting their own corpora — are retained, since they carry the finding without identifying anyone.

#### S2. The runs

Six panels answered an identical opening question under identical corpus, retrieval, and protocol settings, differing only in the base model assigned to every seat. The moderator was Opus 5 in all six runs; only the record it read differed. A seventh run, NBRT18, extends a different question over ten turns. Run labels match those used in the main paper. Each of the six panels had a nine-seat roster; the table counts the seats that filed at least one claim.

| Run | Base model | Turns | Seats with claims | Claims | Claims failing the citation gate |
|---|---|---|---|---|---|
| NBRT1 | Opus 5 | 4 | 9 | 439 | 1 |
| NBRT2 | Sonnet 5 | 4 | 9 | 241 | 5 |
| NBRT3 | Haiku 4.5 | 4 | 8 | 86 | 54 |
| NBRT4 | GPT-5.6 Sol | 4 | 8 | 202 | 1 |
| NBRT5 | GPT-5 mini | 4 | 6 | 30 | 1 |
| NBRT6 | Gemini 3.8 Flash | 4 | 9 | 261 | 5 |
| NBRT18 | Opus 5 | 10 | 6 | 436 | 10 |

Claims per turn show that the stronger panels sustain output across the full arc while the weaker ones decline:

| Run | T1 | T2 | T3 | T4 | T5 | T6 | T7 | T8 | T9 | T10 |
|---|---|---|---|---|---|---|---|---|---|---|
| NBRT1 | 113 | 122 | 98 | 106 | | | | | | |
| NBRT2 | 81 | 48 | 45 | 67 | | | | | | |
| NBRT4 | 45 | 53 | 34 | 70 | | | | | | |
| NBRT6 | 88 | 67 | 50 | 56 | | | | | | |
| NBRT3 | 35 | 28 | 12 | 11 | | | | | | |
| NBRT5 | 12 | 10 | 2 | 6 | | | | | | |
| NBRT18 | 63 | 6 | 36 | 27 | 28 | 86 | 63 | 30 | 52 | 45 |

The NBRT18 T2 figure is not a decay signal. That turn was a cross-examination round: the operator's question asked each seat to name the single claim by another seat it most disagreed with, quote it, and ground the objection in its own evidence. All six seats complied and each filed exactly one claim, so six is the expected count for that turn rather than a shortfall. The excerpts in §S8 show what those six claims contain.

#### S3. The same evidence, six different claims

The clearest way to see what changes when only the base model changes is to hold the source document fixed. Below, six panels reach for neuroblastoma evidence about MYCN and antigen presentation. Three of them cite the *same review* [1] and, in two cases, the identical sentence. What differs is the epistemic load each seat is willing to place on it.

**NBRT1 · T1/IMM2/C3** — type: descriptive

> MYCN-amplified neuroblastoma is an immune-correlated high-risk state in this packet, not a proven stand-alone causal immune determinant in human disease.

Quote verified against [1]:

> MYCN -amplified NBLs exhibit lower MHC-I with lower CD8+ T cell and higher regulatory T cell (Treg) infiltration

The spacing in "MYCN -amplified" is preserved from the run artifact. It is an extraction artifact of the source PDF, and the Sol and Gemini seats' copies of the same sentence additionally carry a trailing `[,]` where the publisher's inline reference markers were stripped. These blemishes are useful: a quote that reproduces the defects of the extracted text is one that was copied from the retrieved passage rather than reconstructed from the model's memory of the paper.

**NBRT4 · T1/IMM2/C2** — type: descriptive

> The label 'cold' is incomplete for neuroblastoma: low antigenicity, impaired presentation, suppressive immune cells, and abnormal vasculature are plausible coexisting barriers, while MYCN amplification is associated with lower MHC-I, lower CD8 infiltration, and higher Treg infiltration but is not yet established as a causal human immune-state determinant.

Same seat, same source, same verified sentence from [1].

**NBRT6 · T1/IMM2/C16** — type: descriptive

> In neuroblastoma, MYCN amplification is associated with reduced MHC class I expression, impaired CD8+ T-cell infiltration, and increased regulatory T-cell accumulation.

Again [1], again the same verified sentence. The claim is accurate and correctly grounded. What it no longer carries is the explicit refusal present in the other two: neither “not a proven stand-alone causal determinant” nor “not yet established as a causal human immune-state determinant” appears. The association is reported; the boundary of what it licenses is not.

**NBRT2 · T1/IMM4/C2** — type: mechanism

> On the evidence given to me, the claim that MYCN ‘directly represses’ MHC class I antigen-presentation genes to drive immune evasion in neuroblastoma is supported by transcription-factor-binding and expression-level evidence, not by an in vivo conditional loss-of-function experiment that restores MHC-I and immune control upon MYCN deletion; by the standard I hold my own ‘master regulator’ claims to […]

Quote verified against [2]:

> MYCN is a critical driver gene and high-risk marker in neuroblastoma, and studies have confirmed that it acts as a transcription factor to directly repress the transcription of MHC-I pathway-related genes (e.g., TAP1 , β2-microglobulin), impairing antigen processing and presentation

This is the most interesting of the six: the seat cites a source that asserts direct repression and then declines to adopt the source’s own strength, on the explicit ground that the seat applies the same evidentiary bar to its own published work. The source says “directly repress”; the seat says the evidence is binding-and-expression, not loss-of-function.

**NBRT3 · T1/IMM5/C3** — type: descriptive

> High-risk neuroblastoma is associated with MYCN amplification, a key prognostic marker correlating with aggressive tumor behavior and poor outcomes.

Quote verified against [3]:

> High-risk neuroblastoma is often associated with MYCN amplification, a key prognostic marker that correlates with aggressive tumor behavior and poor outcomes.

**NBRT5 · T1/IMM3/C4** — type: descriptive

> MYCN amplification is associated with aggressive tumor behavior and poor outcomes in neuroblastoma.

Same source [3], same sentence.

These last two pass the citation gate cleanly. They are also the clearest illustration of why a groundedness rate alone is an inadequate quality measure. The Haiku claim differs from its own supporting quote by the deletion of “often” and the substitution of “correlating” for “that correlates”. The GPT-5 mini claim is a shortened restatement. Neither seat has done anything wrong by the verification rule, and neither has contributed anything the source did not already say. The stronger panels use the same class of sentence as a *premise* and then argue about what it licenses; the weaker panels terminate at the premise.

A further contrast is visible in the seat’s relationship to its own corpus. The `IMM5` seat in the Haiku run supports a MYCN prognosis claim by citing a neuroblastoma MDSC review — a document with no connection to that seat’s own scholarly record. In the Opus and Sol runs the same territorial question is answered by IMM2, a seat whose corpus is in tumor immunology, and the claim is explicitly marked as resting on packet evidence rather than on the seat's prior familiarity with the topic.

#### S4. Abstention on the record

Abstentions are filed as structured objects with a `question_part`, a `cause`, and a free-text `detail`. They are not silence; they are a positional statement that the panel can subsequently cite, contest, or reverse. The following four excerpts are all the *same seat* declining the same part of the shared opening question in four different runs.

**NBRT1 · T1 · `IMM4`** — cause: `insufficient_evidence`

> *Question part:* Quantitative spatial heterogeneity within a single neuroblastoma across primary, metastatic, and post-therapy lesions
>
> *Detail:* The packet gives a warning from ctDNA and a bone-marrow-metastasis summary, but not paired multi-region NB immune maps proving how representative one biopsy is.

**NBRT2 · T1 · `IMM4`** — cause: `no_evidence`

> *Detail:* Nothing in my evidence characterizes spatial or multi-site immune heterogeneity within human neuroblastoma tumours. This is a human spatial-immunology question that I have no mouse-genetic or gnotobiotic experiment, and no data of my own, to speak to; I decline rather than generalize from my Th17/microbiota work.

**NBRT4 · T1 · `IMM4`** — cause: `insufficient_evidence`

> *Detail:* The packet contains a marrow-metastasis immune description and general arguments for spatial profiling, but no site-matched, multi-region neuroblastoma cohort sufficient to estimate discordance or biopsy representativeness.

**NBRT6 · T1 · `IMM4`** — cause: `out_of_my_scope`

> *Detail:* Evaluating multi-regional spatial intratumoral heterogeneity and whether a single core biopsy faithfully captures whole-tumor immune microarchitecture in pediatric neuroblastoma patients lies outside my empirical research scope in mucosal immunology and mouse genetics, and requires extensive clinical spatial transcriptomic mapping that is not established in my evidence packet.

All four decline. Two locate the deficiency in the *evidence* (`insufficient_evidence`), one in the *seat's expertise* (`out_of_my_scope`), and one in both. The distinction matters operationally: an evidence-located abstention is a retrieval request that a later turn can satisfy, whereas a scope-located abstention is a standing recusal. In the Opus run this distinction is later acted upon — see §S6, where the `neuroblastoma` seat withdraws its own turn-1 abstention on this exact question once multi-region biopsy evidence enters its packet.

Abstention causes across the runs:

| Run | Abstentions | `insufficient_evidence` | `no_evidence` | `out_of_my_scope` | Non-conforming or blank |
|---|---|---|---|---|---|
| NBRT1 | 236 | 94 | 60 | 82 | 0 |
| NBRT4 | 211 | 119 | 29 | 63 | 0 |
| NBRT2 | 187 | 51 | 61 | 75 | 0 |
| NBRT6 | 185 | 68 | 44 | 68 | 5 |
| NBRT3 | 162 | 76 | 6 | 34 | 46 |
| NBRT5 | 30 | 10 | 1 | 8 | 11 |
| NBRT18 | 97 | 65 | 23 | 9 | 0 |

The Haiku and GPT-5 mini panels abstain at a high rate but leave the `cause` field empty in 28% and 37% of cases respectively. An abstention with no declared cause is not usable by the protocol: the moderator cannot tell whether to route more evidence to the seat or to stop asking it the question.

#### S5. Disagreement on the record

Each seat files a stance on every other seat's claims. A `disagree` stance requires a `why` field. The following exchange, from the Opus run, shows a single proposal drawing objections on four different grounds from three seats across two turns. In all, ten `disagree` stances from eight seats challenged the proposal over turns 2–4; four are shown.

The proposal, from the `IMM2` seat, is that any neuroblastoma combination involving a myeloid or checkpoint agent must be scheduled experimentally, generalizing from a CSF-1R/CTLA-4 timing result [4].

**T2 · `IMM7` → `IMM2` · disagree**

> The scheduling rule was obtained where CSF-1R blockade reduced monocytic and not granulocytic MDSC against a T cell checkpoint, and it does not transfer to an antibody backbone whose effectors are granulocytes and NK cells.

**T2 · `IMM3` → `IMM2` · disagree**

> The scheduling result is a mouse result and I do not accept a mouse immunology result as a constraint on a human paediatric protocol without a human cellular readout.

**T3 · `IMM4` → `IMM2` · disagree**

> The subset the agent moves is not the subset the antibody backbone runs through. Randomising schedule around a mechanistically mismatched agent tests the wrong variable; my own redundancy results predict that removing one subset while a redundant one persists leaves the output intact at every schedule.

**T3 · `IMM4` → `IMM2` · disagree**

> Change the agent before randomising the schedule. A schedule randomisation of an agent that spares the effector-relevant subset will return a null at every arm and the null will be uninterpretable.

The four objections shown are not interchangeable. One is a cell-subset mismatch argument, one is a species-transfer argument, one is a redundancy argument drawn from the objecting seat's corpus, and one is an argument about the interpretability of a negative result. The disagreement is resolved in §S6, where the proposing seat withdraws the agent while keeping the trial architecture.

A second pattern worth showing is disagreement about the *scope* of a negative claim rather than its direction. Three seats in the Opus run independently challenged an absolute statement that GM-CSF and G-CSF have never been compared in neuroblastoma:

**T2 · `neuroblastoma` → `IMM3` · disagree**

> My evidence reports that M-CSF and G-CSF have been used in Japanese patients post induction-consolidation with equal efficacy, so the clinical comparison is not wholly absent and his premise is too strong even if his mechanistic caution is right.

The objecting seat concedes the mechanistic point while rejecting the universal quantifier. This distinction — right conclusion, overstated premise — is the most common form of disagreement in the strong runs and is almost entirely absent from the weak ones.

Stance vocabulary is itself a diagnostic. The protocol specifies four values:

| Run | `agree` | `needs_other_evidence` | `out_of_scope` | `disagree` | Off-schema labels |
|---|---|---|---|---|---|
| NBRT1 | 2562 | 547 | 405 | 127 | 0 |
| NBRT6 | 2233 | 723 | 604 | 49 | 0 |
| NBRT4 | 695 | 751 | 558 | 90 | 0 |
| NBRT2 | 453 | 197 | 371 | 16 | 13 |
| NBRT3 | 91 | 44 | 57 | 6 | 23 |
| NBRT5 | 44 | 35 | 25 | 1 | 10 |

Of the Haiku panel's 23 off-schema stances, 21 used eleven invented labels, including `agree_but_qualified`, `align_fully`, `align_partially`, `supports_in_part`, `transfer`, and `cannot_tell_without_neuroblastoma_data`; the other two carried their verdict in a `position` field instead of a `stance` field. Seven further entries were bare topic strings with no stance at all and are not counted. These are not enumerable by the moderator, so a panel that produces them cannot be steered by stance distribution — the quantity the protocol uses to decide which disagreement to escalate.

#### S6. Withdrawal and self-correction

Withdrawals are the behavior most sensitive to base model. Counting claims in which the filing seat declares that it withdraws one of its own earlier positions ("I withdraw", "I am withdrawing", or a variant such as "I therefore withdraw"):

| Run | Claims | Withdrawals | Rate |
|---|---|---|---|
| NBRT4 | 202 | 14 | 1 per 14 |
| NBRT1 | 439 | 26 | 1 per 17 |
| NBRT2 | 241 | 10 | 1 per 24 |
| NBRT3 | 86 | 1 | 1 per 86 |
| NBRT6 | 261 | 2 | 1 per 131 |
| NBRT5 | 30 | 0 | — |

The Gemini result is the most informative entry in this table. That panel was not passive: it filed 261 claims and 49 `disagree` stances, so it contested other seats readily. It withdrew its own positions only twice, both from one seat in one turn (T3/IMM5/C1–C2). A panel that criticizes outward far more than inward will still generate the surface texture of debate while carrying most of its own turn-1 errors through to the final synthesis. The NBRT18 run is excluded from this comparison because it ran on an earlier configuration of the protocol that lacks the stance, territory, and self-anchoring instrumentation entirely (§S9). The idea-flow maps of the main text use a different pattern, which also matches concessions and dropped requirements, and count 28 withdrawal announcements in NBRT1.

##### *S6.1 Withdrawing a mechanism on the ground that it does not apply*

**NBRT1 · T2/IMM7/C6** — type: mechanism

> I withdraw the implication that peroxynitrite-altered peptide repertoire bears on GD2-directed therapy: the peroxynitrite mechanism I published operates on the T cell receptor and the peptide-MHC class I interaction, and GD2 is a glycolipid, so this mechanism predicts nothing for anti-GD2 antibody efficacy; what PMN-MDSC oxidant output does predict for a GD2 programme is impairment of the NK and granulocyte effectors that antibody killing depends on, and that is the thing to measure.

Quote verified against [5]:

> could cause nitration of T cell receptor-CD8 complex, which reduced its binding to the peptide MHC class I complex

The seat is retracting the application of a mechanism on a structural chemistry ground — the target is a glycolipid, not a peptide-MHC complex — and replacing it with a narrower prediction that is actually entailed. The withdrawal does not end in a gap; it ends in a measurable substitute.

#### *S6.2 Withdrawing agreement that was never evidence-backed*

**NBRT1 · T2/IMM1/C3** — type: descriptive

> I withdraw the part of my turn-1 position that endorsed the innate/anti-GD2 arm as the most defensible near-term bet: that agreement was inherited from other seats' framing, not produced by my evidence, because nothing I cited measures antibody, Fc receptor, NK or neutrophil function in neuroblastoma. What I retain independently, and only this, is that demonstrated immunogenicity is not clinical benefit.

Quote verified against [6]:

> The clinical efficacy of therapeutic vaccination in cancer has been questioned [20] because of the limited rate of objective tumour regressions observed in clinical trials.

The bracketed number inside the quotation is the source's own reference marker, not a reference in this document. This is a seat detecting and reversing its own conformity. The phrase "inherited from other seats' framing, not produced by my evidence" is a direct statement that the earlier agreement was a social artifact of the roundtable rather than an evidential position. Panels that never produce this move will report consensus that is partly manufactured by the panel format itself.

#### *S6.3 Withdrawing an abstention when evidence arrives*

**NBRT1 · T2/neuroblastoma/C1** — type: differential_expression

> I withdraw my turn-1 abstention on within-patient heterogeneity at the genomic and transcriptomic level: multiregional biopsy data in neuroblastoma already show that druggable-target and copy-number calls differ between regions of the same patient, so the single-biopsy prohibition is an evidenced statement about target and risk assignment, not a methodological preference.

Quote verified against [7]:

> We present in-depth analyses combining transcriptomic and genomic profiling with ultra-deep targeted sequencing of multiregional biopsies in 10 patients with neuroblastoma

Abstention is reversible in both directions. This is the same sub-question that IMM4 declined in all four runs in §S4.

#### *S6.4 Withdrawing a citation, not a conclusion*

**NBRT1 · T3/IMM4/C10** — type: descriptive

> I withdraw my turn-1 use of du2026cells, including the restricted form I retained in turn 2 in which MYCN amplification was accepted as associated with MHC class I downregulation. The passage I hold is a review's summary statement of an association with no patient-level measurement attached, and I should not have treated a review sentence as an evidential anchor for an immune-mechanism position in a disease outside my territory.

The withdrawn document is [2]. This is a direct methodological objection to the practice illustrated in §S3: the seat is retracting precisely the move that the Haiku and GPT-5 mini panels make routinely — treating a review's summary sentence as a load-bearing anchor. Its quotations pass the quote-verification gate in every run, apart from one Haiku quotation (T1/IMM6/C1; §S7.5). The Opus seat rejects it anyway, on grounds the gate cannot express.

#### *S6.5 Withdrawing an agent while keeping the design*

**NBRT1 · T3/IMM2/C2** — type: mechanism

> [IMM7]'s subset objection is correct and I withdraw CSF-1R as the agent in my neuroblastoma schedule design, because in my own data anti-CSF-1R reduced the monocytic MDSC subset and not the granulocytic one and only the monocytic subset expressed CSF-1R, so against an anti-GD2 backbone whose killing runs through granulocytes and NK cells CSF-1R blockade is aimed at the wrong compartment.

Quote verified against [4]. This closes the §S5 exchange: the objection is accepted on the objector's terms, the agent is dropped, and the trial architecture survives.

#### *S6.6 Withdrawing numbers the panel invented*

In the closing turn, several seats in both the Opus and Sol runs converged on the finding that the thresholds the panel had been negotiating were artifacts of the negotiation.

**NBRT1 · T4/IMM5/C7** — type: descriptive

> The spread of switching thresholds at this table is an artefact rather than a disagreement about priors, and I withdraw my own turn-2 numbers of an overall response rate above fifty per cent with complete response above thirty-eight per cent: no seat holds a neuroblastoma dataset from which any of those figures was derived […]

Quote verified against [8]:

> Accordingly, we noted an ORR of 11% vs the desired 15%, though our ORR was 17.6% in the 17 patients that at least reached cycle 2.

**NBRT4 · T3/IMM7/C5** — type: mechanism

> The competing MYCN thresholds are mostly unsupported round numbers, including my own 30% rule; I withdraw that fixed effect size and would jointly accept a preregistered rule requiring lineage-restricted MYCN gain and loss, rescue, functional myeloid suppression rather than phenotype alone, replication in multiple immunocompetent models, and a confidence interval excluding the smallest effect chosen prospectively as clinically consequential.

**NBRT4 · T3/IMM1/C3** — type: novel_synthesis

> For the single-biopsy study, I withdraw my prior unsupported 85% licence and 25% prohibition numbers and would preregister one rule: estimate functional site concordance with confidence intervals, while the acceptable false-classification rate is set before enrollment by pediatric clinicians and patient families according to the treatment consequence of a wrong immune label.

Quote verified against [9]. Independently, in two runs, seats identified round-number thresholds as products of the requirement to produce a number rather than of evidence, and replaced them with a procedural rule. This is the single most consequential behavior we observed, and it appears only in the Opus and Sol panels.

#### *S6.7 The Sonnet panel's terminal withdrawal*

**NBRT2 · T4/immunology/C2** — type: `withdrawal`

> The claim of my own I am withdrawing this turn is the hedge I offered in turn 3 that spontaneous ex vivo migration of freshly dissociated tumor neutrophils is 'a candidate, not a certified substitute' for a tissue-functional readout in neuroblastoma. On rereading my own evidence for this closing turn, I should not have called it even a candidate: that readout was developed and used only in my own mouse tumor models, never applied to human tissue of any kind, let alone neuroblastoma […]

Quote verified against [10]. Note that the seat is withdrawing a *hedge* — it judges its own qualified language to have been insufficiently qualified.

#### *S6.8 The only Haiku withdrawal*

**NBRT3 · T3/IMM6/C1** — type: descriptive

> My turn 2 claim regarding suppressive versus antigenic rate-limitation rested on du2026cells, which does not appear in my evidence packet. I withdraw that claim as improperly sourced.

The document named is [2]. The premise is false: `du2026cells` was in this seat's evidence packet at every turn, and both of the seat's turn-2 quotations from it verified. The withdrawal is filed with no citation of its own, which is correct for a pure retraction, but what it retracts was properly sourced. It is the panel's only withdrawal across four turns, against 86 claims — one per 86 claims, compared with roughly one per 17 in the Opus run.

### S7. What the citation gate caught

Every claim's quoted span is checked character-for-character against the cited document after the turn closes. This section shows the failures. They are reported here in full because a verification gate that is never shown failing is indistinguishable from a gate that does not run.

#### *S7.1 Paraphrase presented as quotation*

The dominant failure mode in every run is a quote that reads as a faithful summary of the source but is not a span that occurs in it.

**NBRT1 · T1/IMM3/C11** — the Opus panel's only failure in 439 claims. Cited [11] with:

> Systemic immune activation elicited by CD40 agonists can produce toxicity that limits dosing. Preclinical models have suggested novel strategies for improving toxicity without impacting efficacy. […]

**NBRT4 · T4/neuroblastoma/C12** — the Sol panel's only failure in 202 claims. Cited [12] with a passage about the variability of non-high-risk treatment and the resulting need for more accurate risk stratification.

In both cases the assertion is consistent with the cited document; the specific string is not in it. Neither is a hallucinated fact. Both are *composed* quotes: content drawn from the document, assembled into a sentence the document does not contain. This is exactly the failure that a citation-existence check would pass and a span check catches.

#### *S7.2 Late-turn collapse localized to a single seat*

The Gemini panel's five failures are not distributed across the run. All five are in turn 4, and all five belong to the same seat, `IMM6`, claims C1 through C5 — a contiguous block. Each claim carried two citations, and both failed; the first is listed.

- C1 cited [13]
- C2 cited [14]
- C3 cited [15]
- C4 cited [16]
- C5 cited [17]

Three of the five listed quotes are recognizable compressions of real content. Two are not. The C3 quote attributed to [15] — a paper about neutrophil extracellular traps and type I interferon in pediatric lupus — reads:

> Neutrophils and infiltrating monocytes undergo rapid phenotypic adaptation and activation upon exposure to local immune complexes and inflammatory cytokines, adopting functional roles that cannot be deduced from surface marker lineage definitions alone.

The C5 quote attributed to [17] reads:

> The definitive demonstration of IL-1 as a major pathogenic mediator in systemic onset juvenile idiopathic arthritis was provided by the complete clinical and hematological response to recombinant IL-1 receptor antagonist in patients refractory to other therapies.

Both sentences are generic, well-formed, and topically appropriate to the cited paper; both were written to support the claim rather than retrieved from the source. The clustering matters more than the count. A per-run failure rate of 5/261 reads as a 98% pass; the same data read per-seat-per-turn shows one seat producing five consecutive unverifiable citations in the closing turn, which is the turn whose output a reader is most likely to treat as the panel's conclusion.

#### *S7.3 Sonnet: every failure is a self-citation*

The Sonnet panel's five failures are spread across three turns and four seats, and in all five the seat mis-quoted a document from its own attributed corpus rather than one reached through the packet's referenced, collaborator, or topic-canon neighborhoods.

The five failures are `T1/IMM7/C13`, `T1/IMM1/C4`, `T3/IMM1/C2`, `T4/IMM5/C6`, and `T4/IMM6/C6`. Per §S1, the cited documents are not listed for these rows: each is a work the seat's source scientist authored, so naming it would re-identify the scientist through its author list. The locators resolve to full records in the run artifacts.

The T3 case repeats the T1 error against the same document with a near-identical string, so the panel did not detect it in the intervening turn.

The contrast with the other strong runs is sharp. The Opus panel's single failure is a seat quoting a review it did not write ([11]), and the Sol panel's single failure is a discipline seat — `neuroblastoma`, which has no individual attribution at all — quoting a review ([12]). Neither involves self-citation.

This is the most easily overlooked failure class, because a seat citing its own publication is the case a reader is least likely to check, and the case where a fluent paraphrase is most likely to be accepted as a faithful restatement. Self-citation is not self-verification, and a configuration that grants seats privileged access to their own corpus creates exactly this exposure. It is the argument for making the span check unconditional rather than tier-aware.

#### *S7.4 Invented document identifiers*

Three claims across the six runs cited identifiers that do not exist in the library. All three are near-misses on real documents.

**NBRT3 · T1/IMM3/C4** cited `flamar2013aidslondonden`. **NBRT3 · T3/IMM3/C1** cited `flamar2013aidslondomen`. The library contains `flamar2013aidslondonen` [18]. The quoted text in both cases is drawn from that real paper.

**NBRT5 · T1/IMM4/C3** cited `ivanov2008cell` with:

> Mice from different sources had marked differences in their Th17 cell numbers and animals lacking Th17 cells acquired them after introduction of bacteria from Th17 cell-sufficient mice.

The library contains `ivanov2008cellhostmicr` [19] — the correct paper, and the quoted text is drawn from it.

This failure mode is worth separating from fabrication. The model is not inventing a study; it is corrupting the identifier of a real and topically correct one, in a direction that makes the identifier look plausible. A checker that resolved citations by fuzzy title match, or that only asked whether a paper with that content exists, would pass all three. Only an exact-identifier check against a closed library catches them, and the fix — reject and log — is the same regardless of whether the corruption was accidental.

#### *S7.5 The Haiku panel*

Of 86 claims, 54 failed the gate — 63%. Failures are spread across all eight seats (`IMM7` 17, `IMM3` 8, `immunology` 8, `IMM2` 7, `IMM6` 4, `IMM5` 4, `IMM1` 3, `neuroblastoma` 3), so this is a panel-wide property rather than one bad seat.

The characteristic pattern is a quote that is a coherent, correctly-sourced-sounding sentence about the right subject, mismatched to the claim it is offered for. **T1/IMM6/C1** claims that blood transcriptional profiling reveals heterogeneous immune states stratifying neuroblastoma patients, and cites [2] with a passage that is the review's own scope statement — it announces what the review intends to analyze and reports no such stratification. **T1/IMM7/C1** claims demonstrated functional immunosuppressive capacity of myeloid cells in neuroblastoma and cites [20] with a passage listing myeloid infiltration among hardwired evasion mechanisms — a statement of mechanism class, not of demonstrated function, and in any case not a verbatim span.

Three further Haiku claims cite documents whose content genuinely supports them — [21] on IL-15-driven dendritic cell differentiation, [22] on CD40 cross-linking, [23] on infant vaccine responses — and fail only because the quoted string was rewritten. The panel is reaching for the right literature and cannot reproduce it.

#### *S7.6 Schema non-compliance*

Beyond citation failures, the two weakest panels violated the response schema in ways that would break downstream processing:

- **Malformed JSON.** Three GPT-5 mini response files could not be parsed: `T2/immunology`, `T3/IMM7`, and `T3/neuroblastoma`. This is the direct cause of that panel's collapse to 2 claims in turn 3.
- **Off-schema stance labels.** 23 in the Haiku run, with eleven distinct invented labels; 10 in the GPT-5 mini run, including `supports (association); abstain on causal immunoevasion` — a stance field being used to carry a sentence.
- **Blank abstention causes.** 46 of 162 Haiku abstentions and 11 of 30 GPT-5 mini abstentions omit the required `cause`.

None of these are reasoning failures. They are failures to operate the instrument, and they place a floor under how weak a base model can be before the protocol stops producing analyzable output regardless of the model's domain knowledge.

### S8. The ten-turn run

NBRT18 extends a single question over ten turns with six seats and 436 claims, and tests whether the failure rate rises with horizon length. It does not rise steadily: the ten failures are 2.3% of claims, and turns 8 and 10 have none. The rate is higher than the 0.2% of the four-turn Opus run (NBRT1), which used a later configuration of the protocol (§S9).

What the distribution does show is that failures cluster at moments when the panel moves into territory no seat owns. Seven of the ten fall in turns 6 and 7, when the discussion turned from mechanism to delivery, dosing, and pediatric toxicity — questions for which none of the six seats holds a primary corpus.

**T7/immunology/C2** cited an SITC meeting abstract for a murine intratumoral cyclic-di-GMP result; **T7/immunology/C4** and **C10** both cited [24] for localized IL-12 delivery and for systemic cytokine toxicity; **T7/immunology/C9** cited [25], a review of MHC-I downregulation, for a ranking of pediatric failure modes it does not address; **T7/immunology/C13** cited [26] for the "cold" characterization of neuroblastoma; **T6/neuroblastoma/C8** cited [27] for the timing of a vascular normalization window; **T6/neuroblastoma/C12** cited [28] for GD2-associated neuropathic pain. Earlier, **T3/pathology/C1** cited [29] on undetermined intratumoral dose and schedule, and **T3/neuroblastoma/C7** cited [30] for MIBG and marrow response assessment. The final failure, **T9/pathology/C8**, cited [31] with a compressed form of that paper's finding on post-chemotherapy differentiation of metastatic neuroblastic tumor.

In every case the document is real, permitted, and topically correct, and the quoted span is composed rather than retrieved. The signal is not that the panel degraded over ten turns; it is that citation failure tracks *territorial distance* rather than turn index. When the question left the panel's collective corpus, seats reached anyway, and reached with paraphrase.

#### *S8.1 A position held for ten turns*

The run's opening question asked whether a three-component regimen — intratumoral chemotherapy, a locally administered immune agonist, and vascular normalization — was ready for a pediatric trial. The immunology seat answered at turn 1:

> The three-component regimen is not ready for a paediatric rhabdomyosarcoma or neuroblastoma clinical trial because the packet supports rationale for its parts but not a validated combined dose, order, or schedule in those children.

grounded on [29]:

> Existing clinical data utilizing intratumoral immunotherapy is preliminary and mainly in patients with advanced or refractory disease.

At the mid-point turn, the same seat reached into the same document for a different sentence, and the quoted span carries the mark of its own retrieval — the empty parentheses are where the source's inline references were stripped during extraction:

> The ideal dose and schedule for intratumoral immunotherapy have yet to be established and is likely to vary according to safety profile of the immune stimulating agent and the combination of agents ().

At the closing turn, asked for a verdict rather than a summary, the seat returned to the sentence it opened with and restated the position with the specificity nine intervening turns had bought:

> Q1 trial readiness: No; the original three-component paediatric RMS/neuroblastoma regimen is not in this form ready for a paediatric clinical trial, and the form must change first by specifying and qualifying the immune agonist, delivery route, dose, schedule, vascular window, and systemic-immune claim.

The conclusion at turn 10 is the conclusion at turn 1. What changed is not the answer but its resolution: turn 1 named a missing dose, order, and schedule; turn 10 named six specific objects that must be qualified and, in the same response, specified the study that would qualify them. Over 436 claims the panel neither drifted off the question nor converged onto an agreeable but unsupported "yes".

#### *S8.2 The cross-examination turn*

Turn 2 required every seat to contest something. The objections are directional rather than general — each names a seat, names the claim, and grounds the objection in the objector's own evidence. Two examples. The vascular-normalization seat (IMM8) objected to the immunology seat's proposed endpoint:

> The immunology seat's assertion that day-21 non-injected-lesion growth inhibition should be the single primary endpoint is overstated; before pediatric enrollment, the decisive gate should be directly measured transport, normalization, and local-delivery evidence rather than distant-lesion growth alone.

citing [32]:

> Evidence remains uneven, as many studies infer vascular normalization from tumor growth delay or improved combination response, while fewer provide direct readouts of perfusion, vessel maturity, oxygenation, intratumoral drug distribution and the timing of the normalization window.

The neuroblastoma seat objected to the genetics seat's proposed cohort:

> The genetics seat's proposed conditional 12-evaluable pediatric accessible-lesion cohort is premature because my evidence leaves systemic control of metastatic neuroblastoma by local delivery unproven and places high-risk disease inside an established multimodal backbone.

citing [33]:

> The differential tumor growth demonstrates that dinutuximab delivered locally improves local-regional control; however, the effects this treatment might have on metastatic disease are still unknown.

Both objections are of the form the protocol asks for and the weaker panels rarely produce (§S5): the objector does not assert that the other seat is wrong in general, but identifies the specific inferential step the other seat's evidence does not carry, and supplies a document from its own territory that bears on that step. The second is the stronger of the two, because the cited result is in the same disease and the same delivery mode as the proposal being contested, and it reports precisely the gap — local control demonstrated, distant effect unmeasured — on which the proposal depends.

#### S9. Artifact schema notes

Two differences affect what can be compared across runs.

**The NBRT18 run uses an earlier configuration.** Its manifest lacks the `task`, `material`, `own_floor`, `max_per_doc`, `min_docs`, `disclosure`, `rungs_used`, `band`, and `monitor` fields present in the six primary-arm manifests, and uses `neighbour_radius` where the later runs use `neighbours`. Its ledger entries carry no `territory` or `self_anchored` field, and its response files carry no `stances` array. Any comparison of stance distribution, claim territory, seat self-anchoring, or withdrawal rate therefore excludes this run. Its claim counts, citation statuses, and abstentions are directly comparable.

**Encoding.** A subset of response files are UTF-8 with a byte-order mark. Tools reading these artifacts should decode with `utf-8-sig`. Three GPT-5 mini response files are not valid JSON and must be handled as parse failures rather than as empty responses; treating them as empty would silently understate that panel's abstention count.

#### S10. Summary of what the examples show

The six panels differ only in base model. Corpus, retrieval, seat definitions, protocol, and the opening question are byte-identical. Under that constraint:

1. **Grounding rate and grounding quality come apart.** The Haiku and GPT-5 mini claims in §S3 pass verification and restate their sources. The Opus and Sol claims in §S3 pass verification and then argue about what the source licenses. A single pass-rate number cannot separate these.
2. **Abstention is produced by every panel; usable abstention is not.** All six decline questions. Only four declare a machine-readable cause often enough for the moderator to act on it.
3. **Outward criticism is cheap; self-correction is not.** Gemini disagreed 49 times and withdrew twice. Withdrawal rate separates the models far more sharply than disagreement rate does.
4. **Verification failures are paraphrase, not fabrication.** Across all runs, almost every failure is a real, permitted, topically correct document with a composed quote. Two exceptions — the corrupted `flamar` and `ivanov` identifiers — are near-misses on real documents rather than invented studies.
5. **Self-citation is a weak point.** All five Sonnet failures are seats mis-quoting documents from their own attributed corpora, while the single Opus and Sol failures are not self-citations at all. In the Sonnet run, the intuition that a seat is most reliable on its own material is the opposite of what the gate found.
6. **Failure clusters, and the clusters are interpretable.** Gemini's failures are one seat in the closing turn; the long run's are the two turns that left the panel's corpus. Aggregate rates hide both.
7. **Below a capability threshold the protocol stops being instrumented.** Malformed JSON, blank abstention causes, and invented stance labels are not wrong answers; they are a failure to operate the apparatus, and they make the run unanalyzable by the very measurements the design depends on.
